# The genomic landscape of adaptation to a new host plant

**DOI:** 10.1101/2023.04.17.537225

**Authors:** Rachel A. Steward, Kalle J. Nilsson, Jesús Ortega Giménez, Zachary J. Nolen, Chao Yan, Yajuan Huang, Julio Ayala López, Anna Runemark

**Affiliations:** Department of Biology, Lund University, Lund, Sweden; Cavanilles Institute of Biodiversity and Evolutionary Biology, Universidad de Valencia, Paterna, Spain; Division of Theoretical Ecology and Evolution, Universität Bern, Bern, Switzerland

**Author notes:** Equal contributions (Rachel A. Steward, Kalle J. Nilsson, Jesús Ortega Giménez).

**Keywords:** adaptive divergence, host plant adaptation, inversion, parallelism, genomic selection, introgression

## Abstract

Adaptation to novel ecological niches is known to be rapid. However, how the loci underlying ecological divergence are coupled to traits reproductively isolating populations, ultimately enabling the formation of persistent species, remains a consequential question in speciation research. Here, we investigated the genomic differences underpinning colonization of a new niche and formation of two partly sympatric host races of *Tephritis conura* peacock flies. We took advantage of two independent sympatric zones west and east of the Baltic Sea, where host plant specialists using the thistle species *Cirsium heterophyllum* and *C. oleraceum* co-occur, and address what regions of the genome maintain the host races in parallel. Using genome-wide association, differentiation and divergence statistics, we identified a large, highly divergent region associated with host use among western and eastern populations. Within this region, we identified unique haplotypes associated with each host race, indicative of a large inversion, adding to the growing body of evidence that structural changes to the genome are important for adaptations to persist in the face of gene flow. We further showed strong signatures of selection in this region, especially in populations of the derived *C. oleraceum* specialist host race. The region also had reduced introgression, especially in western populations, while the rest of the genome showed signs of extensive gene flow. Genes within highly differentiated windows within the putative inversion were not only enriched for functions involved in host adaptation, including phenology and metabolic responses to different metabolites in the two host plants, but also enriched for gametogenesis, fertilization and embryological development, all of which suggest sequence divergence could have large consequences on reproductive isolation between the host races. In conclusion, this study suggests that structural changes in the genome may facilitate the formation of persistent host races, and ultimately speciation, in face of gene flow.

## Introduction

How biological variation and novel adaptation arise are fundamental and unresolved questions in our understanding of the origins of biodiversity. Ecological adaptation to novel environments and food sources are known to result in rapid phenotypic and genetic divergence (Bush, 1969; Boag and Grant, 1981; Feder *et al*., 1988; Filchak *et al*., 2000; Herrel *et al*., 2008; Nosil, 2012), but to what extent this ecological adaptation leads to persistent speciation is contested (Schluter, 2009; Anderson and Weir, 2022; Anderson *et al*., 2023). Both the ecological context, including the consistency (Hendry, 2009; Bolnick, 2011) and multidimensionality (Rice and Hostert 1993; Nosil and Sandoval 2008; Nosil et al. 2009; Chevin et al. 2014)(Rice and Hostert, 1993; Nosil and Sandoval, 2008; Nosil and Harmon, 2009; Chevin *et al*., 2014) of the divergent niches, and the genomic underpinnings of adaptation to those niches (Chevin *et al*., 2014) may influence the likelihood of divergent lineages becoming distinct species over time. Whether ecological adaptations and reproductive isolation are linked, at either the phenotypic or genomic scale, is important to the speciation process. Without reproductive isolation, ecological shifts can lead to collapses of formerly isolated populations, resulting in increased gene flow and breakdown of the speciation process (Gow *et al*., 2006; Taylor *et al*., 2006; Lackey and Boughman, 2017). Thus, a challenge for evolutionary biologists is to identify the genomic architecture that enables persistent species to form (Kulmuni *et al*., 2020).

One way to uncover the genomic architecture that enables speciation is to assess sequence differences that are robust to gene flow between populations in sympatry (Dittmar *et al*., 2016; Ferris *et al*., 2017). Gene flow can be limited in regions of the genome harboring genes that are involved in reproductive isolation, and differences in these regions are likely contributing to the speciation process (Ellegren *et al*., 2012; Coughlan and Matute, 2020; Schluter and Rieseberg, 2022). However, interpreting which of the differences between existing species *caused* speciation may be challenging, as additional genes involved in reproductive isolation may have accumulated since the species diverged (Cruickshank and Hahn, 2014; Ellegren and Galtier, 2016). Investigating the genomic differences between recently diverged taxa that are maintained in independent contact zones can help resolve this challenge (Schluter and Rieseberg, 2022). Regions that repeatedly differ among differentially adapted taxa are likely to play a role in speciation (Johannesson *et al*., 2010; Meier *et al*., 2018; Rivas *et al*., 2018; Bohutínská *et al*., 2021; Koch *et al*., 2022). Extensive theoretical and empirical evidence has shown that genomic architecture coupling coadapted loci, including inversions, is particularly important for persistence of differential adaptation in the face of gene flow (Feder, Roethele*, et al.*, 2003; Kirkpatrick and Barton, 2006; Wellenreuther and Bernatchez, 2018; Faria, Johannesson*, et al.*, 2019; Berdan *et al*., 2022; Schaal *et al*., 2022; Lucek *et al*., 2023). Yet, the extent to which linkage of ecological adaptation and reproductive isolation loci is facilitated by pleiotropic effects of the same genes, coupling of genes through structural variation, or co-inheritance of uncoupled genomic regions upheld by correlational selection remains an empirical challenge for speciation researchers.

Herbivorous insects and the plants they interact with are excellent models for studying the genomic basis of ecological adaptation and speciation. These are two of the most speciose groups of eukaryotes (Mora *et al*., 2011; Christenhusz and Byng, 2016), and specialization of phytophagous insects and their hosts is hypothesized to drive this divergence (Vidal and Murphy, 2018). The specificity and multidimensionality of the niches constituted by the host plants (Hardy and Otto, 2014) and strong dependency on host plants for reproduction and survival (e.g. assortative mating based on host recognition; Bush, 1969) may increase the chance of ecological speciation in herbivorous insects (Hardy and Otto, 2014). Changes in host repertoire, either through host range expansions or host shifts, have previously been associated with chemosensory genes underlying butterfly host choice (van Schooten *et al*., 2020), genes involved in oral secretions in aphids (Nouhaud *et al*., 2018; Boulain *et al*., 2019; Shih *et al*., 2023), genes coding for enzymes in the digestive systems of both butterflies and aphids (Nallu *et al*., 2018; Singh *et al*., 2020; Shih *et al*., 2023), and genes underlying phenological shifts in *Rhagoletis* flies (Feder *et al*., 1988; Feder, Berlocher*, et al.*, 2003; Egan *et al*., 2015; Hood *et al*., 2020; Inskeep *et al*., 2022). However, how the genetic changes underlying ecological adaptation are coupled to reproductive isolation at the genomic level (Butlin and Smadja, 2018), enabling co-existence and long-term persistence of differentially adaptive ecotypes or host races (cf. Dres and Mallet, 2002), remains a challenge to resolve. It is possible the time required for stable existence of host races is determined by the accumulation of genomic architecture coupling host adaptation and reproductive isolation loci. This is the case in birds, where sister taxa with inversions co-exist at shorter divergence times (Hooper *et al*., 2019).

Here, we use the Tephritid fly *Tephritis conura* to uncover the genomic basis of ecological divergence and reproductive isolation promoting stable coexistence of two differentially adapted host races. These host races infest the ancestral host thistle *Cirsium heterophyllum* and the derived host thistle *C. oleraceum,* respectively (Romstock-Volkl, 1997). They differ in phenology and host plant preference (Romstock-Volkl, 1997; Diegisser, Johannesen*, et al.*, 2006), and have host-specific larval performance and survival (Diegisser *et al*., 2008) resulting in relatively strong reproductive isolation (Seitz and Komma, 1984; Romstock-Volkl, 1997; Diegisser, Johannesen*, et al.*, 2006; Diegisser, Seitz*, et al.*, 2006). Moreover, mitochondrial networks and allozyme markers suggest they are genetically differentiated (Diegisser, Johannesen*, et al.*, 2006; Diegisser, Seitz*, et al.*, 2006). Taken together, these findings suggest that there is strong divergent ecological selection acting on these host races. This study system also features independent contact zones where the host races are found in sympatry both west and east of the Baltic Sea (Nilsson *et al*., 2022), enabling us to determine the genomic regions that differ between host races in parallel, and hence are likely to confer ecological adaptation. We show that a large, contiguous region of the ancestral dipteran X chromosome is the basis of genomic differentiation between *T. conura* host races. This region has unique haplotypes for each host race and is highly resistant to introgression, consistent with an inversion. Not only does the putative inversion exhibit signatures of positive selection, especially in the derived host race, but it also couples genes that are enriched for functions critical to host adaptation, mating and reproduction. Together, these results suggest that structural variation has enabled colonization of a novel host and facilitates host race persistence in sympatry.

## Methods

*Tephritis conura* (Loew, 1844; Diptera) is a true fruit fly in the family Tephritidae. Most species in this family are specialists, eating one or a few plants within the same family or genus (Aluja and Norrbom, 2001). Larvae feed on fruits, seeds, flowers, stems or plant roots (Christenson and Foote, 1960; Headrick and Goeden, 1998). Within Tephritidae, several examples of host races occur in the genera *Eurosta* (Craig *et al*., 1993), *Rhagoletis* (Bush, 1969; Feder *et al*., 1988) and *Tephritis* (Diegisser *et al*., 2004; Diegisser, Seitz*, et al.*, 2006). *Tephritis conura* host plant specialization has been documented in populations in continental Europe, following a host shift from *C. heterophyllum* (referred to as CH flies from here on) to the derived host plant *C. oleraceum* (referred to as CO flies; Romstock-Volkl, 1997; Diegisser, Johannesen*, et al.*, 2006). The host races are nearly indistinguishable morphologically, except for the ovipositor length, which is shorter for CO females (Romstock-Volkl, 1997; Diegisser *et al*., 2007; Nilsson *et al*., 2022). Sampling adult flies on their host plants or as larvae in infested buds also facilitates discrimination.

To uncover the genomic regions involved in host plant adaptation in *T. conura*, we assembled and annotated a reference genome and used whole genome resequencing of four populations from each host race. For both host races, *T. conura* larvae inside infested buds were collected during June and July 2018. We sampled buds from eight populations, four for each host race, including sites in allopatric populations of *C. oleraceum* in Germany and Lithuania, allopatric populations of *C. heterophyllum* in northern Sweden and Finland, and sympatric populations of both host races in southern Sweden and Estonia (Fig. 1; Table S1). This design enables us to identify the genomic regions that consistently differ between host races, as the sympatric regions sampled on each side of the Baltic Sea constitute independent contact zones. The pupa used for the reference genome was sampled from a CH individual from the CHST population (Fig. 1D).

**Figure 1.**
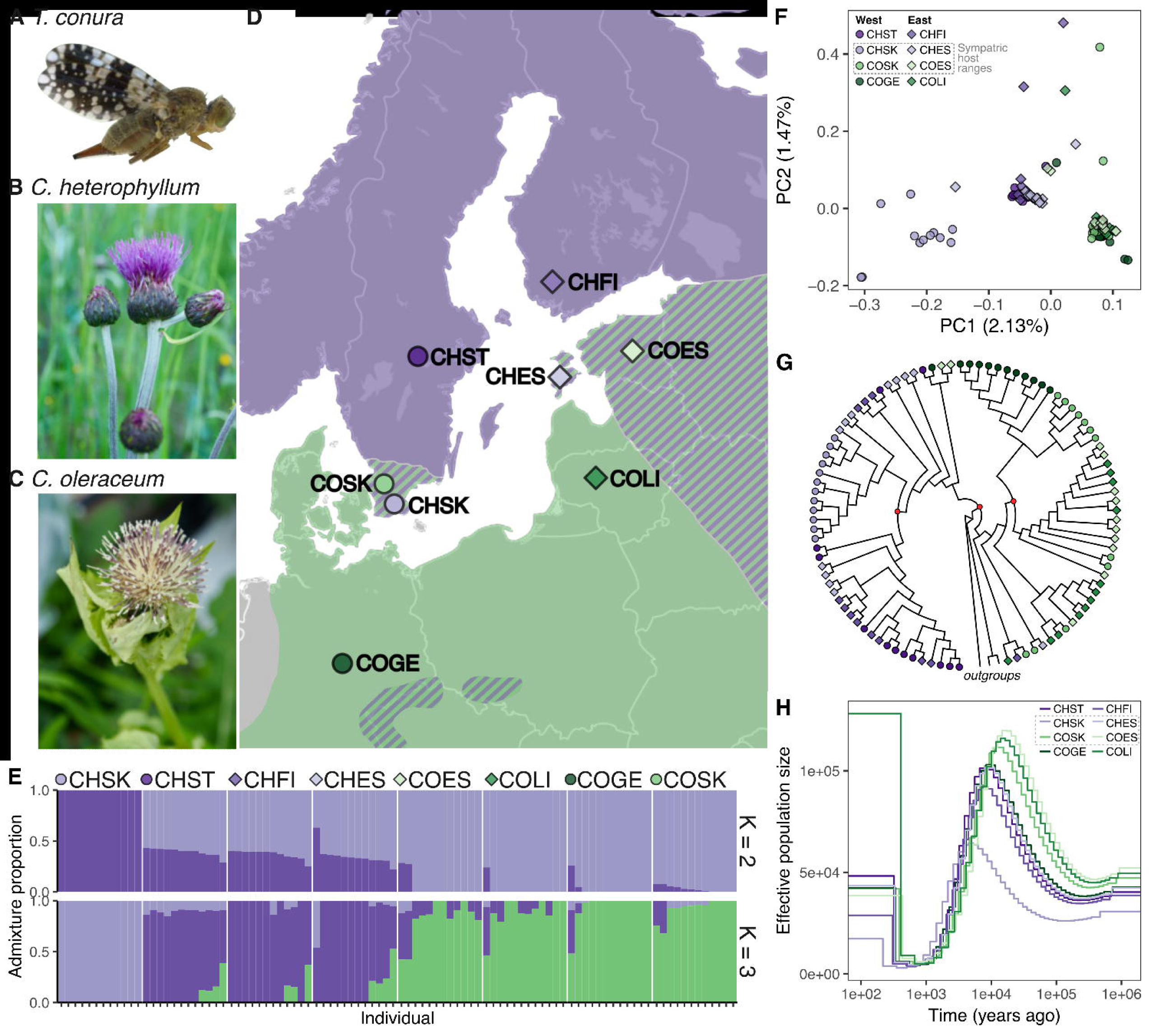
Genomic divergence of Tephritis conura. (A; female pictured) host races specializing on Cirsium heterophyllum (CH; B) or C. oleraceum (CO; C) thistle buds. (D) Flies were sampled from populations of the CH (purple points) or CO (green points) host race along transects east (diamonds) and west (circles) of the Baltic Sea. Transects traversed regionds of northern Europe where *C. heterophyllum* exists in allopatry (purple background), *C. oleraceum* exists in allopatry (green background), or both host plants are found in sympatry (striped background). (E) Admixture proportions estimated for K=2 and K=3 (see Fig. S6A for K4-9). (F) Principal component (PC) analysis of genetic differences among populations separates CH (purple) and CO (green) individuals on the first PC axis (see Fig. S6B for comparison of the second and third PC axes). (G) ASTRAL consensus cladogram of flies used in the study. Terminal nodes are colored by population and internal nodes with strong support (posterior probability > 0.8) are colored in red. For branch lengths and posterior probabilities of all nodes, see Fig. S7. (H) MSMC analysis of demographic history using eight individuals with the most similar coverage from each population and excluding highly divergent or repetitive regions.

Thistle buds containing *T. conura* larvae were stored individually in netted cups at 21°C in a laboratory at Lund University, Sweden. We sampled adult flies three days after eclosion. We considered all flies emerging from a given bud as potential siblings and therefore sampled a single adult male from each bud to avoid including related individuals. Adult flies and the pupae were euthanized through snap freezing and stored at −80°C.

### Genome assembly and improvement

DNA from a single *T. conura* pupa was extracted and sequenced by SciLifeLabs, Solna, Sweden following their in-house protocol for PacBio sequencing. The pupa was from the allopatric CH population from Grythyttan, Sweden (aka CHST). Briefly, the DNA was sequenced using PacBio long read technology and sequences were error-corrected using SMRT Link analysis. Sequencing resulted in 4,173,684 reads with a total of 74.3 Gbp and an N50 read length of 17.8 kb. PacBio sequences were assembled with hifiasm 0.7-dirty-r255 (Cheng *et al*., 2021), which was run using default parameters, followed by purge_dups (v. 1.2.5; Guan *et al*., 2020) to identify and remove heterozygous duplication (e.g. haplotigs) in the resulting assemblies.

We used RepeatModeler (v. 2.0.3; Flynn *et al*., 2020) and RepeatMasker (v. 4.1.2; Smit *et al*., 2015) to identify and soft mask repetitive elements in the *T. conura* reference assembly. For RepeatModeler, we specified the ‘rmblast’ engine and ran the LTR structural pipeline. We ran the-gccalc option in RepeatMasker to account for variable GC content. To put the *T. conura* results in context we accessed repeat summaries for nine of the Tephritid species with published genomes (Sproul *et al*., 2023). While Sproul *et al*. (2023) performed additional repeat screens, we used the output from their first round of masking, in which they used RepeatModeler2.0 (search engine “ncbi”) and RepeatMasker4.1.0 to generate custom repeat libraries.

Contigs were arranged into chromosome-level scaffolds using HiC sequencing data from a second *T. conura* individual. Following the Arima HiC pipeline (v. 02; https://github.com/ArimaGenomics/mapping_pipeline; accessed 06/2023), HiC reads were mapped to the hifiasm assembly with bwa-mem (v.7.17-r1188; Li and Durbin, 2009), paired and filtered (mapQ > 10) using python scripts from the Arima pipeline, and duplicates were marked and removed using Picard tools MarkDuplicates (v2.10.3; Picard toolkit, 2019). The mapped reads were then used to scaffold existing contigs using YaHS (Zhou *et al*., 2022), specifying a minimum contig length of 50 kb and requiring the use of existing contigs (i.e., hifi-asm contigs were not broken by YAHS while scaffolding). The resulting contact map was assessed visually using Juicebox (https://github.com/aidenlab/juiceboxgui; Fig. S1A). Due to the low coverage of our HiC data, we performed downstream analyses at the contig level but used the scaffolding output to cluster contigs within hypothetical linkage groups. We assessed the HiC scaffolds by comparing synteny with the published genome of a close relative, *Rhagoletis pomonella*. We hardmasked repeats and aligned the scaffolded assembly to the *R. pomonella* assembly (GCF_013731165.1_Rhpom_1.0; accessed 11/2022; length = 1223 Mbp, scaffolds = 32,060, scaffold N50 = 72 Mbp) using minimap2 (asm20; Li, 2021) and visualized synteny using circlize (v. 0.4.15; Gu *et al*., 2014). We used a similar approach to compare *T. conura* scaffolds against a *Drosophila melanogaster* assembly (dmel-all-chromosome-r6.48, accessed from FlyBase 01/2023; https://ftp.flybase.org/releases/FB2022_05).

The reference assembly was scanned with Kraken2 (Wood *et al*., 2019) to identify bacterial, viral and fungal sequences. Contigs with a contamination proportion of greater than 10% were removed (150 contigs; 47,038,732 bp), a threshold that was chosen based on a large gap that occurred between 2 and 10%. The ancestral dipteran X chromosome was identified by finding contigs with an excess of hits (at least one hit per million bp, approximately top 5% of contigs) when blasted (tblastn; v.2.11.0; Camacho *et al*., 2009) with a set of *Drosophila melanogaster* X-linked proteins, as the the *D. melanogaste*r X chromosome is homologous to the ancestral dipteran X (Vicoso and Bachtrog, 2013, 2015). This resulted in 102 putative X-enriched contigs, 80.3% of which were clustered on a single HiC scaffold: scaffold_3. We used a similar approach to identify scaffold_7 as homologous with *D. melanogaster* chromosome 4 (aka Muller element F), which was previously identified as the Z chromosome in congener *T. californica* (Vicoso and Bachtrog, 2015).

### Gene prediction and annotation

We annotated the assembly using the BRAKER3 pipeline (Gabriel *et al*., 2024). We generated expressed sequence predictions using RNAseq reads from flash-frozen flies from various life stages, sexes and populations (see Steward *et al*., 2024 for details about extraction and sequencing). Reads were mapped to the genome using the splice-aware aligner HISAT2 (Kim *et al*., 2019). Protein hints were generated from the Arthropoda ODB_11 database (Kuznetsov *et al*., 2022). We used the default BRAKER3 pipeline, but a large proportion of single exon genes motivated us to re-merge the protein-based and RNA-based annotations with TSEBRA, reducing the support necessary for introns from 1 to 0.25. The result was a highly complete annotation (BUSCO v. 5.3.1; Manni *et al*., 2021; 98.0% complete (S: 73.6%, D: 24.4%), 0.8% fragmented, 1.2% missing, diptera_ODB10, n = 3285), with a total of 25,175 genes and 30,632 transcripts. We functionally annotated the longest isoform of the predicted genes with eggNOG emmaper (v. 2.1.5; Huerta-Cepas *et al*., 2018) using default settings. Functional annotations were returned for 17,123 genes, including gene ontology (GO) terms for 10438 of these.

### DNA extraction and sequencing

We extracted DNA for whole genome sequencing from entire flies using Qiagen DNeasy Blood and Tissue Kit (Qiagen Corp., Valencia, CA), following the standard protocol with some small modifications (Supplementary Methods 1). Frozen flies were briefly thawed and ground with plastic pestles in the tissue lysis buffer. Samples were incubated with proteinase K (3.5h, 56°C), and then RNAse was added and the pellet eluted using a 56μl of EB buffer. Library preparation and whole genome resequencing were performed by SciLifeLab (NGI-Sweden, Solna, Sweden) using Illumina TruSeq PCR-free prep kit with an insert size of 350 bp. Sequencing of 2×150 bp paired-end reads was carried out on 8 Illumina HiSeqX lanes.

### Mapping, variant calling and allele frequencies

We trimmed raw whole genome sequencing data using fastp (v. 0.23.2; Chen *et al*., 2018), including removing poly-G tails common on patterned flow cells (-g). We aligned the trimmed reads to the reference genome using bwa mem (v. 0.7.17; Li, 2013), then merged parallel sequencing runs per sample, sorting and indexing the alignments using samtools (v. 1.10; Danecek *et al*., 2021). PCR duplicates were identified and removed using MarkDuplicates in Picard tools (v. 2.23.4; Picard toolkit, 2019). Genome-wide mean coverage was calculated with samtools (Fig. S2). Raw sequence data processing and mapping, along with several latter genomic analyses were performed using a custom Snakemake (Mölder *et al*., 2021).

To account for low coverage, we utilized genotype likelihood-based methods for analyses when possible. To accommodate both genotype likelihood and genotype call-based analyses, we produced both (1) a Beagle file containing genotype likelihoods for the 96 *T. conura* individuals produced using ANGSD (v. 0.940; Korneliussen *et al*., 2014) and (2) a VCF containing genotype calls for the 96 individuals as well as three outgroup individuals produced using bcftools (Danecek *et al*., 2021). The filters and steps to produce these two files are described in Supplementary Methods 2. The final Beagle file and VCF contained 10,421,531 and 32,991,264 SNPs, respectively.

Per population, we generated site allele frequency indices (SAF) with ANGSD using the *T. conura* reference to polarize the ancestral state and relying on the same filters as described for the Beagle file, excluding those used for SNP calling (Supplementary Methods 2). We produced folded 1D and 2D site frequency spectra (SFS) for each population and population pair using the SAF files in realSFS (Nielsen *et al*., 2012).

### Population clustering

To explore and visualize genetic clustering of the individuals, we used a principal component analysis performed in PCAngsd (v. 0.982; Skotte *et al*., 2013) and then formally assessed clustering using NGSAdmix (v. 33; Skotte *et al*., 2013) for 1-10 clusters (K). To prepare input files for these tools, we estimated pairwise linkage disequilibrium between all SNPs within 100 kb of each other using ngsLD (Fox *et al*., 2019). We then used the included prune_ngsLD.py script to produce a list of linkage pruned SNPs, excluding edges from the initial graph between sites with a distance greater than 50 kb and with an *r^2^* of less than 0.1 (--max_dist 50000 -- min_weight 0.1). We then subset our Beagle file to these linkage pruned sites and used it as input for both PCAngsd and NGSAdmix. We also performed PCAs using PCAngsd for each contig, and used PC1 to evaluate patterns of genetic clustering across the genome. Contigs with fewer than 500 sites in the pruned Beagle file were excluded. For the clustering analysis, we performed between 10-100 replicate optimizations for each K with a maximum of 4000 iterations per replicate. Replication was halted when the replicates converged or 100 replicates had completed, considering replicates as converging when the three highest likelihood replicates came within 2 log-likelihood units of each other, as in Pečnerová et al. (2021). We selected the best K as the highest level of K achieving convergence.

### Coalescent-based phylogenetic analyses

We reconstructed the phylogenetic relationship between samples using a coalescent-based approach as implemented in ASTRAL-MP (Yin *et al*., 2019), which takes into account the discordance between gene trees and the true species tree expected in recently diverged taxa. For all *T. conura* samples and the three outgroup samples we produced consensus FASTA sequences for all 2,419 complete single-copy Dipteran BUSCO genes present in our assembly using ANGSD, calling the consensus base with the highest effective sequencing depth (-doFasta 3), excluding positions with an individual sequencing depth less than 3 or greater than 100. For each gene, we inferred the maximum likelihood gene tree using IQ-TREE2 (v. 2.2.6; Minh *et al*., 2020), testing and selecting the best fitting substitution model and performing 1000 ultrafast bootstrap replicates to assess node confidence. We collapsed nodes with bootstrap support of less than 10% into polytomies using Newick Utilities (v. 1.6; Junier and Zdobnov, 2010) as collapsing low confidence nodes in gene trees can improve species tree estimation accuracy (Zhang *et al*., 2018). Finally, we inferred the coalescent phylogenetic relationships from BUSCO gene trees with sequencing data for all samples (2,418/2,419 genes) using ASTRAL-MP v. 5.15.5.

### Demographic history

Demographic history was estimated for each population using MSMC2 (v. 2.1.2; Schiffels and Durbin, 2014). We generated the individual VCF and mask bed file by using bcftools (v. 1.14; Li, 2011) and bamCaller.py script from msmc-tools (https://github.com/stschiff/msmc-tools; accessed 09/2023). Sites with extremely low or high coverage, defined as 0.5x and 2x mean coverage respectively, were masked before using Samtools (v. 1.16) to compute the average sample coverage and generate a BED file per-sample. We merged the VCF and mask files (msmc-tools, generate_multihetsep.py script) and used the resulting file as input for MSMC2. To scale the result, we used a mutation rate of 3.46[×[10^-9^ mutations per site per generation, which is derived from estimates in *D. melanogaster* (Keightley *et al*., 2009) and a mean generational interval of one year (Romstock-Volkl, 1997). Because this software is sensitive to read coverage, neutrality, and repetitive content, we compared the MSMC2 output for three sets of samples (the full set of samples, the individuals with highest coverage in each of the eight populations, and the individuals with coverage closest to the global mean in each of the eight populations) at three levels of VCF filtering (the VCF output from bcftools + bamCaller.py script described above; the VCF excluding all repetitive content, and the VCF excluding all outlier windows identified by population branch structure analysis described below).

### Genome-wide association analysis with BayPass

Genome-wide scans for association with ecotype were performed with BayPass (v. 2.4; Gautier 2015). We converted the filtered VCF to allele frequencies for each population with bcftools (v. 1.19; Danecek *et al*., 2021) and removed invariant sites using an in-house script. The dataset was further filtered to account for linkage by subsampling every 20th variant across each contig into 20 unique sets of SNPs. We analyzed each set of SNPs first under the BayPass core model using the default options, then under the auxiliary model using a Markov Chain Monte Carlo (MCMC) algorithm and host race as a covariate. Three independent runs (using the option-seed) were performed for each of the 20 SNP sets. Support for host race association was evaluated using the mean Bayes Factor of these three runs. Results from the 20 independent runs were combined into a single dataset for all sites, and reported in deciban units, with 10db corresponding to 10:1 odds, 20db to 100:1 odds, etc. The mean Bayes Factor was further summarized across 50kb windows. Because several windows were highly repetitive, we removed windows containing 500 or fewer SNPs (1% of the window size) from downstream analyses and visualizations.

### Population differentiation and divergence

To measure population differentiation, *F*_ST_ was calculated for each site, first by generating the site frequency spectrum (2D-SFS) for each population pair, based on which we calculated per-site *F*_ST_ using a Bhatia estimator (Bhatia *et al*., 2013) as implemented in ANGSD (realSFS *F*_ST_). Average pairwise *F*_ST_ was estimated genome-wide and for nonoverlapping 50kb windows (realSFS stats2). We used population branch statistics (PBS) to assess differentiation between sympatric populations of the two host races when corrected for divergence between populations within host races. This accounts for heterogeneity in background differentiation between populations, as differentiation within host races should reflect genetic drift and the recombination rate landscape whereas differentiation between host races should also reflect changes in nucleotide diversity resulting from selection and introgression. We calculated PBS for each site (realSFS stats2; Fig. S3) for four population triads composed of a focal sympatric population, and the sympatric and allopatric populations of the other host race in the same transect.

To measure population divergence, we first re-calculated the 2D-SFS for every 50kb window, then calculated absolute nucleotide divergence (dxy) using a modified script from (https://github.com/marqueda/PopGenCode/blob/master/dxy_wsfs.py; accessed 11/2023 and modified to run in R). We then calculated relative node depth (RND; Fig S3), which quantifies genetic divergence between two populations (here, two sympatric populations) relative to a third population (usually an outgroup, but here an allopatric population). RND is therefore analogous to PBS, as it controls for divergence within host races while testing for divergence between host races. Because several windows were highly repetitive, we removed PBS and RND windows with less than 20% coverage from downstream analyses.

### Outlier analysis and gene ontology term enrichment

To identify genomic regions associated with host use we identified outliers for three different metrics: BayPass (estimated for all eight populations), PBS, and RND (both assessed for a focal sympatric population of each host race in the western and eastern transects, for a total four combinations). Windows that exceeded the genome-wide mean by three standard deviations in at least two of these metrics were identified as outliers. We identified outliers for the CH and CO host races in the western and eastern transects separately, and defined highly differentiated windows as those that were shared between comparisons in the western and eastern transects.

We performed gene set enrichment analysis for gene ontology (GO) terms using the package TopGO (Alexa and Rahnenfuhrer, 2022), only considering terms that were assigned to a minimum of five genes (minimum node size = 5). We tested for enrichment of biological processes, molecular functions and cellular components. Significant enrichment was tested with one-sided Fisher’s exact tests corrected with the parent-child algorithm (Grossmann *et al*., 2007). GO terms were considered significantly enriched with a p-value < 0.05. We further generated a STRING database from the *T. conura* genome annotation (STRG0A87NSB; https://string-db.org/; updated 12/2023) and identified networks within our highly differentiated genes genes using Markov clustering implemented in STRING (inflation = 3). We report the 8 largest networks and associated GO terms, KEGG pathways and enriched reactomes.

### Nucleotide diversity and divergence from neutrality

We evaluated nucleotide diversity (π) and divergence from neutrality, as characterized by Tajima’s D, to understand genome-wide and local differences among populations. ANGSD was used to calculate pairwise nucleotide diversity for each site (thetaD, realSFS saf2theta). These were used to estimate Tajima’s D and pairwise differences across the entire genome and over 50kb non-overlapping windows (thetaStat do_stat). To calculate π, pairwise differences were divided by the total number of sites used in each window. For both metrics, we excluded windows with less than 20% coverage from downstream analyses.

### Selection statistics

To identify SNPs associated with host races and different biogeographic scenarios, we used SelScan (v. 1.3; Szpiech and Hernandez, 2014) to calculate the cross-population extended haplotype homozygosity (XP-EHH). To prepare the data for XP-EHH analysis, SNPs unpruned for linkage disequilibrium were phased with SHAPEIT2 v 2.r837 (Delaneau *et al*., 2013) using default MCMC parameters (7 burn[in MCMC iterations, 8 pruning iterations, and 20 main iterations), conditioning states for haplotype estimation (K = 100), and window size set at 0.5 Mb as recommended for whole genome sequence data. We also used an effective population size (Ne) of 494, based on estimates calculated with SNeP (v. 1.1; Barbato *et al*., 2015). Here, we used the mean across populations, as all estimates were very similar. XP-EHH was calculated in windows of size 100 kb in each direction from core SNPs, allowing decay curves to extend up to 1 Mb from the core, and SNPs with MAF < 0.05 were excluded from consideration as a core SNP. As we lacked a fine[scale genetic map for *T. conura*, we assumed a constant recombination rate of 1.73 cM/Mb as found for the related species *Bactrocera cucurbitae*. Scores were then normalized within contigs with the norm version of Selscan v. 1.3.0.

The XP-EHH method identifies genomic regions that underwent a selective sweep in one population but remained variable in the second population. We calculated XP-EHH for six population pairs to test for differences in selection between host races in sympatry and allopatry, and to test for differences between western and eastern transects. For each pair, we identified putative locality[specific sweeps by selecting those SNPs in the top 1% of the genome-wide distribution. Genes under selection were detected by comparing the SNPs detected by the XP-EHH test against a gene annotation for *T. conura* with bedtools window 2 kb upstream and downstream of these genes (v. 2.29.2; Quinlan and Hall, 2010). We also used a sliding window approach, estimating the average normalized XP-EHH score across all SNPs in 50 kb nonoverlapping windows. More negative values indicate a stronger signature of positive selection in one population in a pair while more positive values indicate a stronger signature of positive selection in the other population.

### Introgression statistics

We investigated evidence of introgression between populations by estimating genome-wide Patterson’s D (Green *et al*., 2010; Durand *et al*., 2011) and *f_4_*-ratios (Patterson *et al*., 2012) for all possible population trios, as implemented in Dsuite’s DTrios function (v0.5-r52; Malinsky *et al*., 2021). To assess variation in introgression across the genome in sympatric populations, we investigated *f_dM_* in windows across the genome for eight focal trios, as implemented in the script fourPopWindows.py (https://github.com/simonhmartin/genomics_general; accessed 02/2024). We selected this statistic as it is designed to detect introgression in localized regions of the genome and is suitable for detecting introgression between both P1 and P3, as well as P2 and P3 (Malinsky *et al*., 2015). Briefly, for each of the four sympatric populations as P2, we estimated *f_dM_* in non-overlapping 50 kb windows, assigning the closest allopatric population of the same host plant to P1, with P3 being either the closest allopatric or sympatric population of the alternate host plant. For all analyses we utilized a *Tephritis hyoscyami* individual (sample ID P18705_127) as the outgroup.

## Results

Our final assembly of the *T. conura* genome was 1.99 Gbp in 2,713 contigs (N50 = 1.75 Mbp). HiC scaffolding with YaHS arranged these contigs into seven large (>100Mbp) linkage groups comprising over 1.77 Gbp of the sequence genome (N50 = 323 Mbp). HiC scaffolds were highly syntenic with *Rhagoletis pomonella* linkage groups (Fig. S1), and linkage group (LG3) was syntenic with the *Drosophila melanogaster* X chromosome, which is homologous to the ancestral dipteran X (Vicoso and Bachtrog, 2013, 2015). Benchmarking with Dipteran single copy orthologs found 97.0% completeness with low duplication (2.4%), making it not only the longest but among the most complete Tephritidae genomes available to date (Fig. S4). It was also the most repeat rich genome by more than 20%, driven by an expansion of long terminal repeats that make up over 22% of the genomic content (Fig. S5; Table S2). This history of genomic expansion and high repetitive content, which increase the opportunity for both structural change and chromatin state differences, suggests that *T. conura* has the potential for extensive and rapid genomic divergence driven by structural variants.

### Genomically divergent host races

Using eight continental *T. conura* populations sampled from regions of host plant allopatry and sympatry east and west of the Baltic Sea (Fig. 1D), we found considerable evidence that the *C. heterophyllum* (CH) and *C. oleraceum* (CO) host races represent distinct lineages. The two host races separated under the best supported model in an Admixture clustering analysis (K = 3; Fig. 1E; see Fig. S6A for remaining K). In a genome-wide principal component (PC) analysis, individuals are generally separated by host race on the first PC axis, explaining 2.13% of the variance (Fig. 1F; see Fig. S6B for PC2 and PC3). The samples also split into two distinct branches in a consensus tree constructed using ASTRAL from 3285 single copy ortholog gene trees (posterior probability = 1, Fig. 1G, see Fig. S7 for scaled branch lengths and posterior probabilities for all nodes). These results were supported by pairwise genome-wide *F*_ST_ and dxy comparisons, which showed the largest differentiation and divergence occurred between populations of different host races (Fig. S8, S9). The western sympatric CH population (CHSK) appears to be the most distinct lineage, separating from both CO individuals and the remaining CH individuals along PC1, and forming a separate cluster under the best supported model in the clustering analysis (K = 3). This population also formed a distinct cluster in the ASTRAL consensus tree constructed from 3285 single copy ortholog gene trees (Fig. 1G).

Despite clear evidence of divergence between the two host races, we also found signs of recent introgression or incomplete lineage sorting among the sampled flies, especially in eastern populations. In the consensus tree, there were five individuals (3 CO, 2 CH) that were not assigned to the same major branch as the rest of their host race (Fig 1G). Combined with the clustering analysis, which showed considerable admixture in some individuals (Fig. 1E), and the PCA, in which several individuals of both host races were highly divergent on the second PC axis (1.47% of the variance; Fig 1F), these results suggest some level of admixture in these individuals.

Although it was previously hypothesized that the derived host race arose just before the onset of the last glacial maximum c. 18,000 years ago (Diegisser, Seitz*, et al.*, 2006), an MSMC2 analysis suggests host races diverged much earlier, between 0.5 - 1 million years ago (Fig. 1H). These results broadly confirm findings from PCA and admixture analyses. In particular, the highly divergent CHSK population again appears as a distinct lineage, perhaps having diverged from the other CH populations much earlier than the estimated divergence between host races. However, similar patterns can arise when populations have been affected by recent bottlenecks, and we are as yet unable to disentangle these alternative explanations.

### Genomic regions underlying host race formation

To identify genomic regions associated with host race divergence, we tested for allele frequency associations with host race across all eight populations using BayPass. We further evaluated genomic differentiation and genomic divergence among population triads east and west of the Baltic Sea using population branch statistics (PBS) and relative node depth (RND). PBS leverages pairwise *F*_ST_ comparisons among three populations (triads) to assess differentiation and signatures of selection, while RND uses dxy in an analogous framework to PBS to compare divergence and introgression (Fig. S3, S10-S12).

We found strong evidence for a large, contiguous region of linkage group 3 (LG3), spanning 104 Mb, 89 contigs and containing 1269 genes, being highly genomically divergent between the host races (Fig. 2). This region was consistently supported in tests for whole-genome association with host race across all populations, as well as PBS and RND comparisons in all triads (Fig. 2A-C, S13). In each case, we identified outlier windows for each statistic as those falling more than 3 standard deviations outside the mean, and considered outlier windows with support from at least two metrics as highly differentiated between the two host races for CH and CO triads (west: Fig. 2D, east: Fig. 2E). Overall, highly differentiated windows were very consistent among host race comparisons and transects, with 214 windows shared across all four comparisons out of 289 total windows (Fig. 2F). All of these 214 windows were located within the contiguous region on LG3.

**Figure 2.**
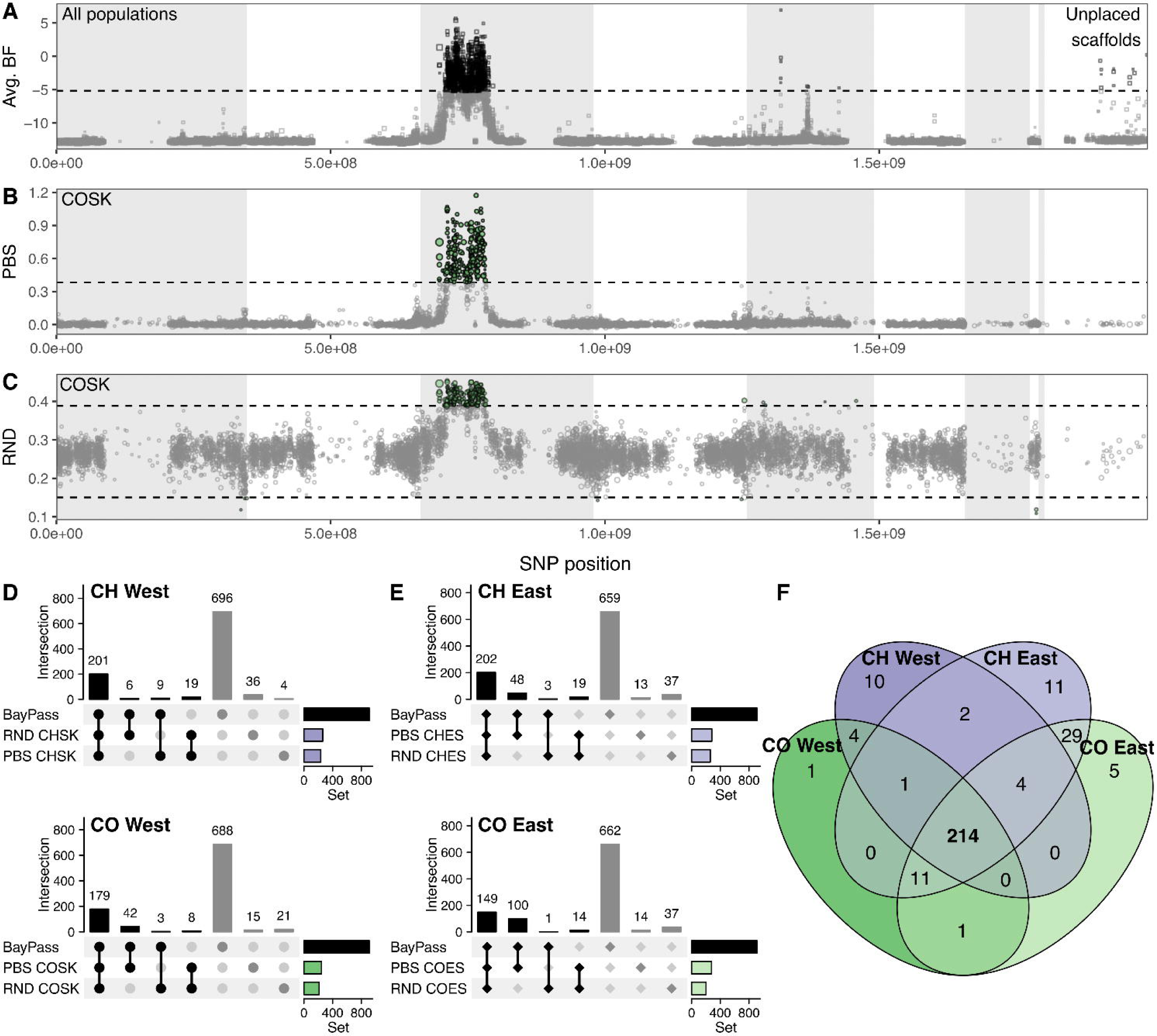
Highly divergent genomic windows between host races. Based on (**A**) BayPass analysis for association with host race using all 8 populations (**B**) population branch statistics (PBS), and (**C**) relative node depth (RND) for COSK, the sympatric CO population in the west, as a representative (see Fig. S13 for remaining triads). Statistics were evaluated over 50 kb non-overlapping windows. Windows were considered outliers when greater than three standard deviations away from the mean (dashed line). (**D**) Overlap between outlier sets within CH and CO comparisons within western and eastern transects. Windows were considered highly differentiated if they were outliers in at least two of the three metrics tested. (**H**) Overlap between all highly differentiated windows.

This dense accumulation of highly differentiated windows on LG3 suggests a large region with reduced recombination, possibly under strong selection in one or both host races, as might be expected for an adaptive inversion. To evaluate this hypothesis, we used PCAngsd to generate local PCAs for each contig in our assembly (Fig. 3A). Within the highly differentiated region, we found two distinct clusters on PC1 for each of the host races (Fig. 3C), consistent with two inversion haplotypes. There was considerably higher variance among CH individuals on PC1, but it is unclear whether this is due to recent introgression into the CH haplotype or ancestral variation among CH populations predating the formation of the inversion and divergence of the CO host race. Interestingly, for contigs outside of the putative inversion, there was little evidence of population structure (Fig. 3B, D), suggesting that gene flow is high throughout the rest of the genome. The local PCA revealed several individuals (4 CO, 3 CH from all except the western sympatric populations CHSK and COSK) with intermediate PC1 values throughout the entire putative inversion, which may indicate that these are hybrid individuals. However, due to the high variance among CH individuals, it is not clear whether these individuals are truly heterozygous for the CH/CO haplotypes.

**Figure 3.**
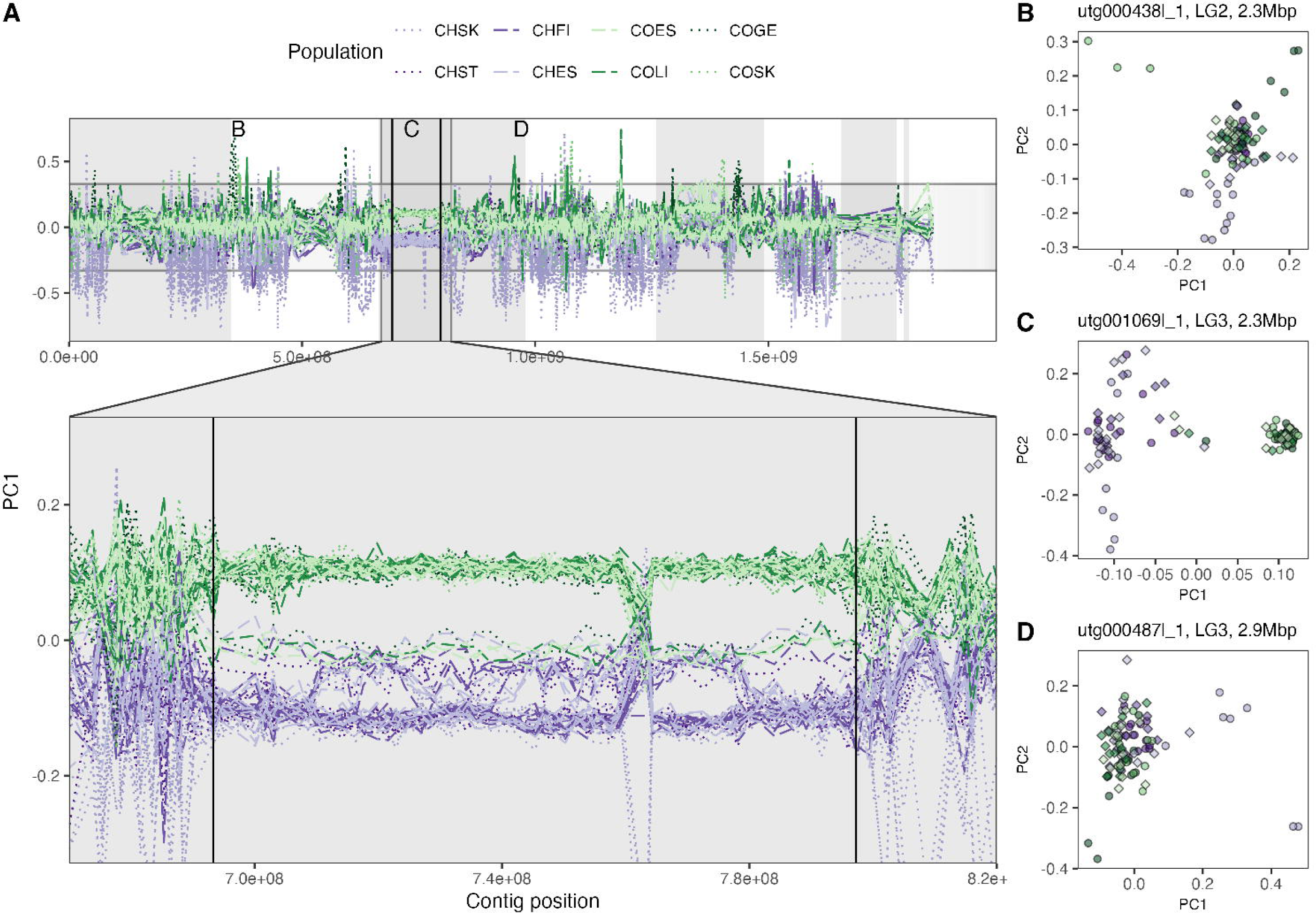
Host race-specific haplotypes within the highly differentiated region suggest a large inversion. (**A**) PC1 across ordered contigs in the genome for all 96 individuals. Zoom panel highlights the putative inversion, with potential break points indicated by black lines. Contigs were included if more than 200 filtered variants were used to calculate the PCA and are ordered according to HiC scaffolds (Fig. S1). Hypothetical linkage groups are shown with alternating gray and white bands. The right-most white band represents unplaced scaffolds. The convergence of haplotypes at 760 Mbp is likely a scaffolding error. (**B**-**D**) PCA of two contigs outside and one inside the putative inversion. Locations of contigs in panels B-D are indicated in A.

### Strong selection on the derived host race (CO) within the inversion

To understand the selective pressures acting on the putative inversion, we tested for evidence for selective sweeps, and whether these are stronger in the derived CO host race. To do this, we assessed Tajima’s D, nucleotide diversity (π) and extended haplotype homozygosity (XP-EHH; Fig. 4). When considering all windows across the genome, π within highly differentiated windows was lower than in non-differentiated windows for all populations (Fig. 4A-B, S14; Table S3). This effect was consistently larger in CO compared to CH populations (Table S3), suggesting selection has disproportionately decreased nucleotide diversity in the putative inversion in the CO host race. Tajima’s D was similar among populations (Fig. 4C-D, S15; Table S4). The exception was CHSK, which had a much higher Tajima’s D, adding to the evidence from Admixture and MSMC analyses suggesting that this population has experienced a recent bottleneck. While Tajima’s D tended to be lower in highly differentiated windows compared to non-differentiated windows, especially in allopatric populations, Tajima’s D was higher in highly differentiated windows in two sympatric populations (CHSK and COSK).

**Figure 4.**
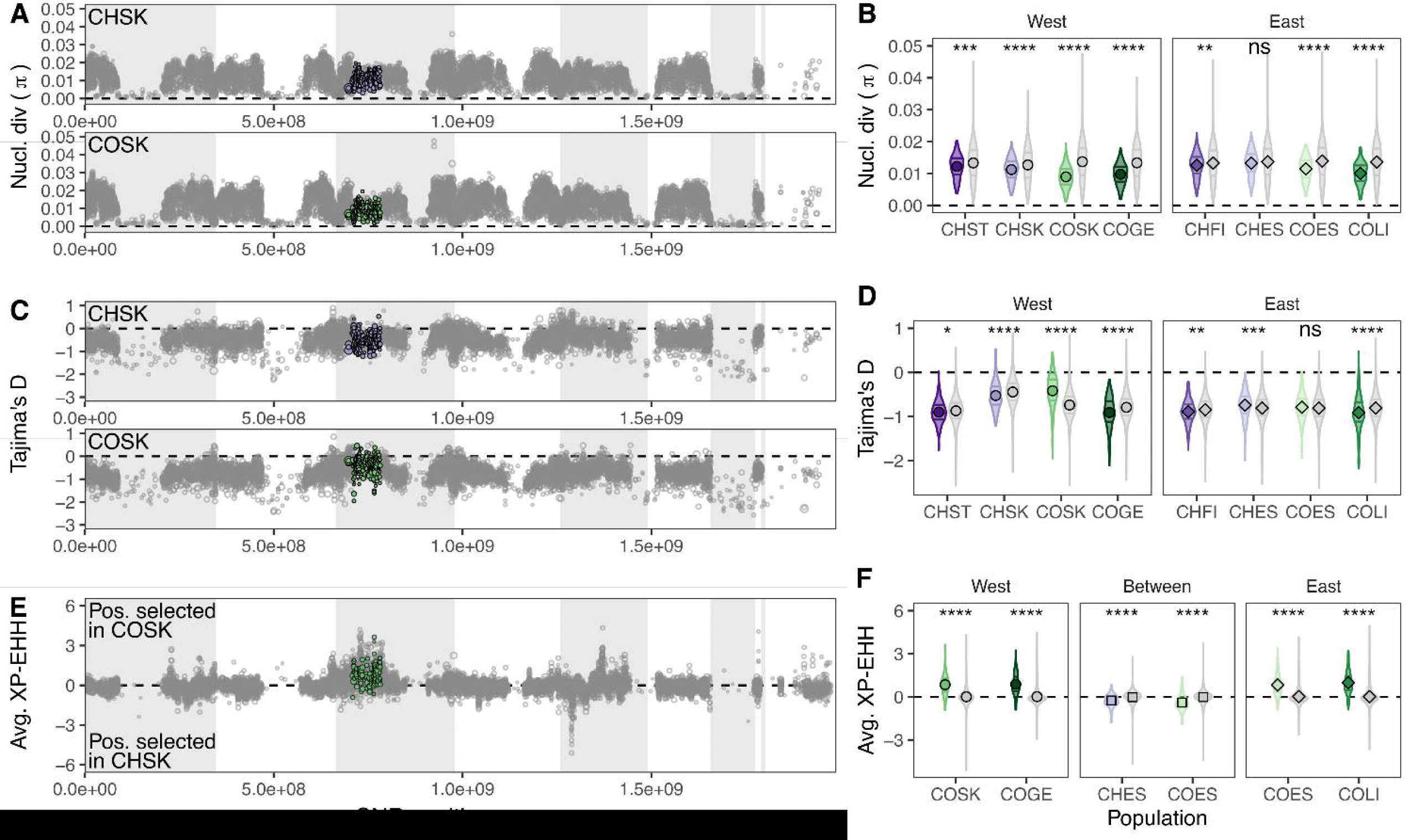
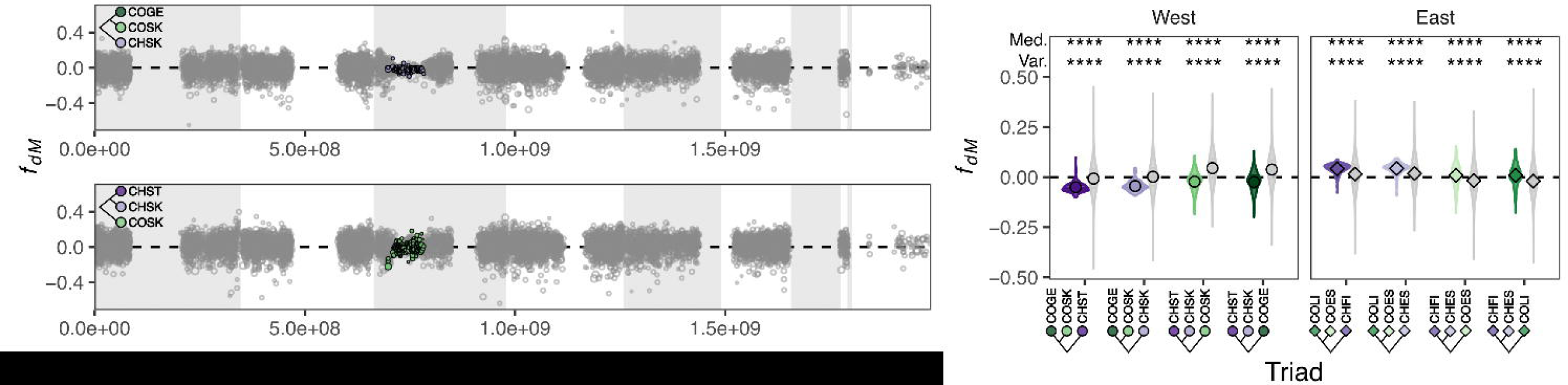
Signatures of selection across the genome. (**A**) Nucleotide diversity (π) in 50 kb windows across the genome for two representative populations (CHSK and COSK, see Fig. S14 for remaining populations). Windows with lower than 20% coverage were excluded. Highly differentiated windows are colored by population. **(B)** π in highly differentiated windows and undifferentiated windows for all populations. Horizontal lines in violin plots show 25th, 50th, and 75th quantiles and points indicate the mean. (**C**) Tajima’s D in 50 kb windows across the genome for two representative populations (CHSK and COSK, see Fig. S15 for remaining populations). (**D**) Tajima’s D in highly differentiated and undifferentiated windows for all populations. **(E)** Average XP-EHH in 50 kb windows across the genome for a representative comparison (COSK vs. CHSK, see Fig. S16 for remaining populations). **(F)** Average XP-EHH in highly differentiated and undifferentiated windows for contrasts between sympatric populations and between allopatric populations of different host races in the west and east, and between western and eastern sympatric populations of the same host race (‘Between’). Medians were compared between highly differentiated windows and undifferentiated windows using Mann-Whitney tests (p < 0.05 *, p < 0.01 **, p < 0.001 ***, p < 0.0001 ****, Table S3-S5).

Extended haplotype homozygosity statistics revealed considerably stronger positive selection in CO populations when contrasted with CH populations (Fig. 4E-F, S16). In all comparisons between CH and CO population pairs, the mean normalized XP-EHH was significantly more positive in highly differentiated windows compared to non-differentiated regions, indicative of stronger positive selection in CO populations (Fig. 4F, Table S5). We found that the difference in XP-EHH was similar in all comparisons between host races (Fig. 4F, Table S5), despite our expectation that positive selection for host adaptation or reproductive isolation should be stronger in sympatric than in allopatric populations (Kautt *et al*., 2020). Meanwhile, in comparisons within host races between contact zones, there was evidence of stronger selection in sympatric populations in the western transect (CHSK and COSK) compared to the eastern transect, although the effect was much smaller than between host race comparisons (Fig. 4F, Table S5).

We further identified significantly selected SNPs in each population for each pairwise contrast using the top 1% of SNPs. Between 42-48% of these SNPs overlapped genes, including the 2,000 bp up and downstream of the coding regions. We found more genes overlapping selected SNPs in the CH host race (Table S6), as SNPs under selection in CO populations were more densely clustered in a smaller number of annotated genes compared to CH populations (Table S6). Furthermore, these selected genes in CO populations were more likely than those in CH populations to overlap with genes in highly differentiated windows or with genes in the putative inversion (Fig. S17), adding evidence that selected SNPs in CO populations fall in a more concentrated, more differentiated set of genes.

### The putative inversion is a region of reduced recombination

Despite extensive evidence for prezygotic reproductive isolation between the *T. conura* host races (Nilsson *et al. in prep*), we detected introgression between the host races in both the eastern and western sympatric regions. D-statistics suggest higher levels of introgression between host races in the eastern than the western region, consistent with this being an older contact zone for these host races (Table S7).

Introgression was not evenly distributed across the genome (Fig. 5A), as illustrated by variation in *f_dM_*in 50 kb windows. Values of *f_dM_* are expected to be positive when introgression occurred between P3 and P2, and negative if it occurred between P3 and P1. *f_dM_* values in highly differentiated windows had a much narrower spread around the median than did undifferentiated windows, suggesting these regions are more resistant to introgression (Fig. 5B, S18; Table S8). Interestingly, introgression in highly differentiated regions differed between western and eastern transects (Fig. 5B, Table S9). In the west, *f_dM_* in highly differentiated regions was consistently lower than undifferentiated regions, suggesting more introgression between allopatric populations of different host races, rather than between sympatric populations of the different host races. Negative values are consistent with greater introgression between the more geographically distant populations, which may reflect historical gene flow. These patterns suggest outlier regions have increased resistance to introgression between sympatric populations. The pattern was opposite in the east, with greater introgression between sympatric populations than allopatric populations within the highly differentiated regions.

**Figure 5.**
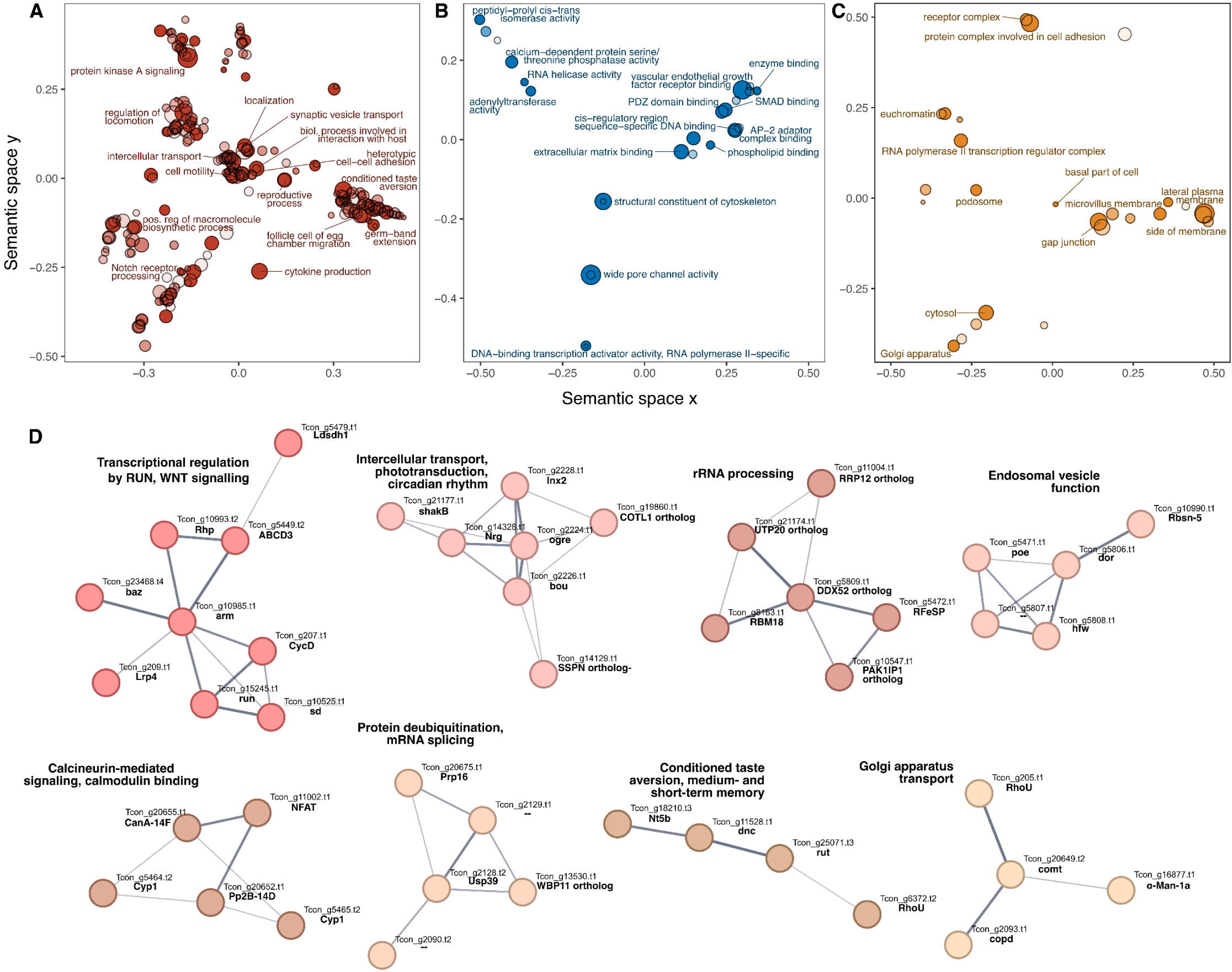
Introgression in across the genome. (**A**) Introgression was compared between triads of *T. conura* populations using *f_dM_*. Triads were composed of two populations of the same host race (P1 = allopatric, P2 = sympatric) and a third population of the other host race (P3). Pictured are *f_dM_*in 50 kb windows for two representative P3 populations, CHSK and COSK. Values close to zero indicate little to no introgression, negative values indicate greater relative introgression between P1 and P3, while positive values indicate greater relative introgression between P2 and P3. (**B**) Introgression in highly differentiated windows (colored according to P3) and undifferentiated windows (gray) for all eight populations. Medians were significantly higher in the west and lower in the east in highly differentiated windows compared to undifferentiated windows (Mann-Whitney, p < 0.0001 ****, Table S9), while variance around the median was consistently significantly lower in highly differentiated windows (Levene’s test; Table S8).

### Highly differentiated windows enriched for host adaptation and reproductive functions

To understand the possible functional consequences of the putative inversion, we performed gene set enrichment analyses of genes overlapping highly differentiated windows (278 genes; Fig. 6A-C; Table S10). Genes within the putative inversion were enriched for numerous functions that may be directly or indirectly related to adaptation to different host plants. Genes involved in both renal development (including malpighian tubules in flies; Bunt *et al*., 2010; Fig. 6A; table S10) and gut development were enriched. Both of these tissues are involved in processing and excreting toxic substances and waste (Gautam *et al*., 2017; Colombani and Andersen, 2020). Notably, response to toxic substances was also enriched, and was associated with glutathione S-transferase and carboxylesterase genes within the highly differentiated region. Both gene families are known to be involved in generalized detoxification of plant metabolites (Low *et al*., 2007; Yu *et al*., 2009; Cruse *et al*., 2023).

The two host plants differ significantly in their budding phenology, and accordingly CH and CO have divergent flight times, even within sympatric host ranges (Romstock-Volkl, 1997). Numerous terms involved in circadian rhythmic processes and diapause were enriched among highly differentiated genes (Table S10). In a network analysis of highly differentiated genes, the second largest network was associated with both phototransduction and circadian rhythm processes and pathways (Fig. 4D). While it has not been thoroughly tested, it appears adult diapause termination is mediated by temperature cues rather than photoperiod (Diegisser, 2005). Accordingly, we also found that thermosensory behavior was significantly enriched in highly differentiated genes.

While both ability to eat the alternate host plant and differing phenologies serve as major reproductive barriers between the host races (Diegisser *et al*., 2008), we also found highly differentiated genes enriched for functions involved in potential intrinsic barriers. We recovered enrichment of biological process terms involved in oogenesis, spermatogenesis, fertilization, embryological development, embryological growth, and mitosis (Table S10). Other processes broadly important to biological development were also highly enriched, including cell signaling and intra- and intercellular transport. Regulation, transcription, post transcriptional modification, and degradation of RNAs, including mRNAs and ncRNAs, were also enriched, suggesting large divergence in coding and noncoding gene expression between the two host races. Several of the eight largest networks recovered in the network analysis of highly differentiated genes were associated with transcriptional regulation and post-transcriptional modifications of RNAs (Fig. 6D). Together, these enriched processes and networks suggest that the highly differentiated region contains many genes that could be critical to reproductive isolation, either by limiting gametogenesis, fertilization and embryogenesis, or by disrupting critical biological processes like cell-signaling and gene expression.

We also found that genes encoding mating behavior and especially male courtship behavior were highly enriched (Fig. 6A), especially conditioned taste aversion, which was annotated to, among other genes, the bitter gustatory receptor *Gr33a*. This receptor has previously been found to be sensitive to pheromones in *Drosophila melanogaster*, and is also sensitive to a range of antifeedants (Moon *et al*., 2009) and may mediate either identification of host plants, where peacock fly courtship occurs, and/or courtship behavior itself. Calcineurin-mediated signaling and calmodulin binding were the main functions of a network of six highly differentiated genes (Fig. 6D). Calmodulin is involved in *D. melanogaster* courtship (Sato *et al*., 2019), possibly mediated by its role in odorant binding receptor sensitization (Jain *et al*., 2021)

## Discussion

While the genetic basis underlying ecological adaptation is increasingly well understood (Bomblies and Peichel, 2022), it remains unclear how the genomic regions associated with differential adaptation confer or are coupled to reproductive isolation, enabling the formation of persistent species (Butlin and Smadja, 2018; Kulmuni *et al*., 2020). We document a large inversion that differentiates the ecologically differentially adapted host races of *T. conura*, reducing gene flow and enabling persistent adaptation. The inversion is differentiating the host races in parallel in two independent regions of secondary sympatric existence, as expected under parallel selection (Johannesson, 2001; Stuart *et al*., 2017), supporting its role in ecological adaptation.

Inversions have been increasingly appreciated as key to enabling co-existence of ecologically divergent lineages in sympatry (Hooper and Price, 2017; Faria, Chaube*, et al.*, 2019). And yet, how the inversions enabling the formation of stable morphs, divergence or speciation originate remains a consequential question (Faria, Johannesson*, et al.*, 2019; Lucek *et al*., 2023). We document recent and extensive expansions of long terminal repeats, constituting over 20% of the genome in *T. conura*, rendering genome size in *T. conura* 55% bigger than that of the closely related species *Rhagoletis pomonella*. While this expansion and its effects on genomic architecture is biologically interesting, analyzing the genomic content of the repetitive regions is intractable, and they have therefore been excluded from the population genomic analyses. Importantly, this repeat expansion has likely facilitated a number of structural changes, including the inversion enabling host race divergence, and producing a more contiguous assembly required for determining the exact nature of the break points and thus origin of the inversion is an exciting venue for future research.

The inversion spans 89 contigs (104 Mb) scaffolded to LG3, covering 33% of the linkage group, and shows consistent and high divergence between the host races. This region of LG3 is syntenic with contiguous regions in the genome of *R. pomonella*. Individuals sampled from the different host races consistently have native host plant specific inversion haplotypes, with only 7 out of 96 individuals potentially having heterozygous inversion haplotype, and none being homozygous for the alternative inversion haplotype. Jointly, the discrete nature of the divergent genomic region and its strong association with host race suggest that this inversion is key for enabling the formation of two host races of *T. conura*. Recent research has shown that inversions are important for maintaining both discrete morphs within species (Jones *et al*., 2012; Küpper *et al*., 2016; Tuttle *et al*., 2016; Faria, Chaube*, et al.*, 2019), facilitate speciation (Wellenreuther and Bernatchez, 2018; Faria, Johannesson*, et al.*, 2019), and enable coexistence with sister taxa after a shorter period of time (Hooper and Price, 2017).

Our findings suggest that the inversion may be important for coupling (cf. Butlin and Smadja, 2018) ecologically co-adapted traits in *T. conura* through physically linking genes involved in ecological adaptation and reproductive isolation. Functional enrichment analyses suggest that this inversion has linked genes important to multiple ecological traits associated with divergent host plant use, including phenology (Romstock-Volkl, 1997), host preference (Diegisser *et al*., 2008), or physiology including digestion of plant defense chemicals (Diegisser *et al*., 2008) and thus facilitate coexistence between the host races. For instance, genes encoding response to toxic substances and renal- and gut development are enriched, which is expected as the host races suffer very strong extrinsic inviability as larvae when feeding on the wrong host plants (Diegisser *et al*., 2008).

Moreover, genes involved in male courtship behavior and conditioned taste aversion, potentially key for host plant recognition and instrumental for mating to take place on the native host plant as observed in *T. conura* (Romstock-Volkl, 1997) are enriched in the inversion. Genes involved in diapause, relevant to the divergent phenologies of the host races (Romstock-Volkl, 1997) were also enriched. The combination of enrichment of genes involved in performance on the host plants, preference for the host plant and phenology which differs between host plants (Romstock-Volkl, 1997; Diegisser *et al*., 2008) adds to evidence that the inversion has coupled (Butlin and Smadja, 2018; Kulmuni *et al*., 2020; Schluter and Rieseberg, 2022) genes involved in ecological adaptation and those contributing to reproductive isolation.

Another set of functions that were enriched in the inverted outlier region included regulation, transcription, post-transcriptional modification and degradation of mRNA and ncRNA. Further studies assessing the extent to which gene expression is differentiated between the host races are necessary to assess the importance of altered regulation of gene expression in host plant adaptation. Furthermore, while we had a prior expectation of enrichment of genes involved in ecological adaptation to using the different thistle species based on the strong documented selection pressures, the most enriched gene functions were instead oogenesis, spermatogenesis, sperm DNA decondensation and fertilization. The enrichment of these functions is not easily interpreted, as it either could reflect selection for reproductive isolation or potentially merely result from LG3 harboring the inversion being syntenic to the ancestral dipteran X-chromosome and therefore enriched for *Drosophila melanogaster* X-linked proteins. Although LG3 not the sex chromosome in *T.* conura, it has over evolutionary history been subjected to sex chromosome-specific selection different to selection on autosomes due to e.g. the lower effective population sizes of sex chromosomes (Charlesworth *et al*., 2018), their potential lower rates of recombination (Charlesworth, 2017) and the faster X (Mank *et al*., 2009), all expected to increase divergence. An additional intriguing possibility is that lower levels of gene flow could result in the typically more divergent sex chromosomes, with a large role in reproductive isolation, representing the ancestral phylogeny despite substantial introgression homogenizing autosomal regions (Fontaine *et al*., 2015). These processes could complicate the interpretation of genomic regions or genes important for ecological adaptation on these contigs, and disentangling the history of the chromosome and the selection pressures shaping it in detail remains a topic for future research.

Instead, the Z chromosome in Tephritis species is likely homologous to *D. melanogaster* chromosome 4 (aka Muller element F), as previously described by Vicoso and Bachtrog (2015). Using BLAST, we found that LG7 in our assembly is the most homologous to *D. melanogaster* chromosome 4. However, LG7 was highly repetitive in *T. conura*, averaging 91-97% repetitive content in 50 Kb windows (Fig. S5). As a result, windows within this chromosome were excluded from most analyses. Future comparisons of coverage across the genomes of males and females and gene expression data from reproductive tissues will shed further light on the sex chromosome in *T. conura* and its importance for host race formation.

Other major model systems where strong, divergent ecological selection has resulted in the formation of host races or ecotypes include *R. pomonella* apple maggot flies infesting hawthorn and apple, respectively (Feder *et al*., 1988; Filchak *et al*., 2000; Feder, Berlocher*, et al.*, 2003), the parapatric ecomorphs of *Littorina* snails adapted to either wave-exposure or crab predation (Johannesson and Johannesson, 1996; Hollander *et al*., 2005; Faria, Chaube*, et al.*, 2019; Perini *et al*., 2020), and *Timema* walking sticks adapted to be cryptic on different host plants (Nosil *et al*., 2002; Nosil, 2007; Nosil and Sandoval, 2008). In contrast to our findings in *T. conura*, strong selection pressures generally maintain ecologically divergent populations in sympatry despite more incomplete reproductive isolation in these systems. *Rhagoletis* flies are isolated by phenological difference and host preference (Feder, Roethele*, et al.*, 2003; Inskeep *et al*., 2022), but the genomic differences that underlie the differently adapted ecotypes is polygenic and can rapidly be recovered through selection on standing genetic variation within the ancestral hawthorn host race (Egan *et al*., 2015). In contrast, the genomic basis conferring mimicry on different host plants in *Timema* walking consists of few loci of large effects, and the background genomic divergence is lower (Villoutreix *et al*., 2020; Nosil *et al*., 2023). The genomic basis of the *Littorina* ecotypes is similar to that in *T. conura* in that variants within inversions contribute to approximately half of the ecotype divergence (Koch *et al*., 2022). In contrast to *T. conura*, several smaller inversions segregate in *Littorina*, enabling a gradual change in both phenotype and genotype in intermediate selection environments (Morales *et al*., 2019; Stankowski *et al*., 2024).

Compared to those systems, the *T. conura* host races have relatively high levels of genetic divergence and strong reproductive isolation, arising from the combination of preference, phenology and performance in the alternative niche (Diegisser, Johannesen*, et al.*, 2006; Diegisser, Seitz*, et al.*, 2006; Diegisser *et al*., 2007; Diegisser *et al*., 2008; Nilsson *et al*., 2022). Potentially, the adaptation is more multifarious, the genomic architecture has been permissive for coupling i.e. by physical linkage (Butlin and Smadja, 2018; Faria, Johannesson*, et al.*, 2019), or adaptation may have taken place during a longer period of time. In contrast to previous divergence time estimates based on mitochondria, MSMC analyses suggest that host race divergence may be old. *Tephritis conura* CH and CO populations could be older than 500 kya as there is no convergence between the host races in the MSMC plots even at this time scale. Further exploration of the dependency of the MSMC inference of which fractions of the genome are included would be necessary to draw any firm conclusions though. Given the extreme morphological similarity of the host races (Nilsson *et al*., 2022), it may seem surprising that divergence times exceed previous estimates of just before the last glacial maximum. However, our new estimates are consistent with the high level of genomic divergence and limited introgression on LG3 observed in this study. While divergence is limited across the rest of the genome, this could reflect gene flow rather than evolutionary divergence time (Fontaine *et al*., 2015). Furthermore, we believe this timeframe is also in accordance with strong reproductive isolation in this system, evidenced by phenological differences (Romstock-Volkl, 1997), oviposition preference and larval mortality on the new host plant (Diegisser *et al*., 2008). Hence, *T. conura* provides a window into a later stage of ecological speciation continuum (Stankowski and Ravinet, 2021) with the potential to offer interesting insights into the development of reproductive barriers following ecological specialization.

### Conclusions and future directions

In spite of mounting support for inversions having a role in speciation (Jones *et al*., 2012; Küpper *et al*., 2016; Tuttle *et al*., 2016; Hooper and Price, 2017; Wellenreuther and Bernatchez, 2018; Faria, Chaube*, et al.*, 2019; Faria, Johannesson*, et al.*, 2019; Lucek *et al*., 2023), studies that uncover how they contribute to coupling of genes underlying ecological and reproductively isolating traits, enabling the formation of persistent species are scarce (but see Meyer *et al*., 2023; Moan *et al*., 2023). We find compelling evidence for a large inversion enabling host race specific adaptation and progress towards speciation in *T. conura*. The enrichment of both genes underlying the ability to metabolize host race defense chemicals and gene functions putatively important for mate choice, host plant recognition and phenology which constitutes an important reproductive barrier between host races could imply that this is an empirical example of such coupling through strengthening of physical linkage between genes within the inverted region. Future studies should focus on determining the evolutionary origin of the inversion, including if recruitment of genes to the inverted region has taken place, and experimentally assess if host race specific inversion karyotype genes are associated with ecological and reproductive isolation.

The extensive and recent repeat expansion also means that *T. conura* has the potential to become a study system for investigating the role of transposable elements to contribute in reproductive isolation or to serve as a source for novel variation through altering regulation of gene expression and remodeling chromosomes, has recently been recognized (Serrato-Capuchina and Matute, 2018). To which extent recent TE releases has contributed to the potential to adapt to novel niches, and to reproductively isolate the host races of *T. conura* is an interesting venue for future research that could help shed light on the role of TE releases in ecological speciation. Improved long-read assemblies of the host races would be an exciting future venue, enabling new insights into the chromosomal structural variation underlying host race formation and persistence. *Tephritis conura* exemplifies ongoing ecological speciation of two host races and future research on this study system will offer unique insights into changes in the genome enabling the formation of persistent species to arise during adaptation to novel niches.

## Data availability

All code used will be deposited on GitHub upon acceptance. The reference genome will be available from NCBI and all resequencing data will be available at NCBI/ENA upon manuscript acceptance.

## Author contributions

AR conceptualized and funded the study and performed field work. KJN and JOG performed field work, and JOG performed all molecular lab work. Bioinformatic analyses were performed by RAS and ZJN, with contributions from JOG, YH, CY and YH, and oversight from AR. The first draft of the manuscript was written by RAS and AR, with contributions from KJN, ZJN, and JOG. RAS, AR, and KJN made significant revisions to the text for subsequent versions.

## Acknowledgements

This work was funded by a fellowship from the Wenner-Gren foundations, a Swedish Research Council starting grant, and additional grants from the Crafoord Foundation, Erik Philip Sörensens Stiftelse and Carl Tryggers Stiftelse to AR covering field-, sequencing- and salary costs. Grants from The Royal Physiographic Society of Lund to JO contributed to field work (Young Researchers Endowment) and RNAseq (Nilsson-Ehle Endowment) costs. A Lund Djurskyddsfond grant to JO covered additional field work expenses. Authors acknowledge support from the National Genomics Infrastructure in Stockholm funded by Science for Life Laboratory, the Knut and Alice Wallenberg Foundation and the Swedish Research Council, and SNIC/Uppsala Multidisciplinary Center for Advanced Computational Science for assistance with massively parallel sequencing and access to the UPPMAX computational infrastructure. We thank J. Johanneson for support when starting the work on this study system, H. Vogel for help with long-read extractions, E. Kärrnäs and M. Schnuriger for assistance during field work, and S. Jacobsen Ellerstrand for exploration of the sex chromosome.

## Notes

### Competing Interest Statement

The authors have declared no competing interest.

### Summary of Updates

This version of the manuscript has been revised to update results of analyses after re-trimming and mapping the fastq files. We also masked repetitive content in all analyses of differentiation, divergence, selection and introgression. Results are now presented to reflect a new, hypothetical linkage map. We also added a new local PCA analysis to support the conclusion that an inversion differentiates the two host races. These new results did not change our broad conclusions about ecological speciation in Tephritis conura flies.

