## Supplementary Materials for "The genomic landscape of adaptation to a new host plant"

### Supplementary methods

#### Supplementary methods 1: Whole genome extractions

Flies were ground with plastic pestles in 1.5 ml Eppendorf tubes using half the volume recommended of buffer ATL. We used the remaining ATL buffer to rinse the pestle over the Eppendorf tube to avoid any loss of tissue. Samples were incubated with 25 µl of proteinase K for 3.5 h at 56 °C. Then, we added 4 μl of RNAse (Qiagen) and incubated samples at room temperature for 2 min. After centrifuging for 5 minutes, samples were washed with AW2 buffer. We used Qiagen Elution Buffer (EB, Qiagen) rather than AE Buffer to elute the samples. In addition, to increase the concentration of DNA in the final eluate, we used 57 μl, instead of 150 μl of the elution buffer after warming it for 30 min at 56ºC. We carefully deposited the EB buffer onto the membrane and left it incubating at room temperature for 6 minutes in order to increase the final DNA yield. After extraction, we stored DNA samples in Qiagen Elution Buffer (Qiagen) at -80 °C.

#### Supplementary methods 2: Filters and variant calling

For all *T. conura* samples, we produced a variant site Beagle file containing genotype likelihoods and a VCF file containing hard genotype calls for all samples using ANGSD (v. 0.940; Korneliussen et al. 2014). As an outgroup was required for some genotype call based analyses we performed (D-statistics), three outgroup individuals (samples P18705_115 - *Tephritis kogardtauica*, P18705_123 - *Tephritis cometa*, P18705_127 - *Tephritis hyoscyami*) were additionally included in the VCF, but not the Beagle file.

To produce the Beagle file (-doGlf 2), we calculated genotype likelihoods using the GATK model (-GL 2), with the following filters: uniquely mapping reads with a mapping quality of 30 (-uniqueOnly 1 -minMapQ 30), data for half the individuals in the dataset (-minInd 48), and a global sequencing depth greater than four times the total number of individuals and less than two times the mean coverage (11X) times the total number of individuals (-setMinDepth 384 -setMaxDepth 2112). We called variable sites with a SNP p-value of less than 2e-6 and a minor allele frequency of greater than 0.05 inferring major and minor alleles and allele frequencies from the genotype likelihoods (-SNP_pval 2e-6 -minMaf 0.05 -doMajorMinor 1 -doMaf 1; Kim et al. 2011). We additionally only included positions outside of repetitive regions using the -sites option.

We produced the VCF by calling genotypes using the BCFtools multiallelic caller (v.1.18; Danecek et al. 2021) on all samples including the three outgroups, disabling BAQ computation (--no-BAQ) and using only reads with a minimum mapping and base quality of 30 (--min-MQ 30 --min-BQ 30), and excluding repetitive regions (-R). We removed variants with a variant quality of less than 30 using bcftools filter (-e’QUAL < 30’) and set genotypes to missing if they had a read depth lower than 3 or higher than 100 or a genotype quality less than 20 using bcftools +setGT. Finally, we filtered the VCF to include only positions with biallelic SNPs.

Per *T. conura* population, we produced site allele frequency index files (SAFs) and minor allele frequency files (MAFs) using ANGSD. For both, we utilized the same options for estimating genotype likelihoods as for the beagle file (-GL 1 -uniqueOnly 1 -minMapQ 30 -C 50 -baq 1; -sites to exclude repetitive content), adjusting sample size-based filters to match sample sizes for each population (-minInd N/2 -setMinDepth N*4 -setMaxDepth N*2*11). For the SAF files, we polarized allele frequencies using the *T. conura* reference genome. For the MAF files, we limited output to the variable sites present in the dataset wide Beagle file using the -sites option and a sites file produced from the first three columns of the Beagle file. We forced sites to take the major and minor allele identities inferred in the Beagle file (-doMajorMinor 3). This is necessary for the dxy estimation to ensure that sites that are not variable in all populations are included in all MAFs and that frequencies are polarized to the same alleles.

### Supplementary tables

Table S1. Collection locations for sampled populations.

| **Population ID** | **Host Plant** | **Transect** | **Host plant range** | **Location** | **Lat** | **Long** |
| --- | --- | --- | --- | --- | --- | --- |
| COSK | *C. oleraceum* | West | Sympatric | Scania | 55.905000 | 13.413000 |
| COGE | *C. oleraceum* | West | Allopatric | Germany | 51.112800 | 9.597117 |
| COLI | *C. oleraceum* | East | Allopatric | Lithuania | 55.652206 | 21.938014 |
| COES | *C. oleraceum* | East | Sympatric | Estonia | 57.879930 | 26.248673 |
| CHES | *C. heterophyllum* | East | Sympatric | Estonia | 58.436099 | 22.937953 |
| CHFI | *C. heterophyllum* | East | Allopatric | Finland | 61.352485 | 22.056471 |
| CHST | *C. heterophyllum* | West | Allopatric | Sweden | 59.628468 | 14.577117 |
| CHSK | *C. heterophyllum* | West | Sympatric | Scania | 55.864156 | 13.745842 |

Table S2. Gene and repetitive content in the *T. conura* assembly.

| **Genomic feature** | **Subfeature** | **Number of elements** | **Length occupied** | **% of sequence** |
| --- | --- | --- | --- | --- |
| ***Gene content*** | | | | |
| Coding genes | | 27588 | 560418.81 | 28.12 |
| ***Repetitive content*** | | | | |
| SINEs | All | 35584 | 6677374 | 0.34 |
|  | Penelope | 28948 | 11459310 | 0.57 |
| LINES | All | 903575 | 370283439 | 18.58 |
|  | CRE/SLACS | 0 | 0 | 0 |
|  | L2/CR1/Rex | 69742 | 45454845 | 2.28 |
|  | R1/LOA/Jockey | 166317 | 42994323 | 2.16 |
|  | R2/R4/NeSL | 652 | 595640 | 0.03 |
|  | RTE/Bov-B | 308283 | 100788706 | 5.06 |
|  | L1/CIN4 | 463 | 69970 | 0 |
| LTR elements | All | 1245635 | 421195757 | 21.13 |
|  | BEL/Pao | 313664 | 101290113 | 5.08 |
|  | Ty1/Copia | 93378 | 30085145 | 1.51 |
|  | Gypsy/DIRS1 | 597579 | 232357378 | 11.66 |
|  | Retroviral | 4653 | 1417735 | 0.07 |
| DNA transposon | All | 577305 | 186403442 | 9.35 |
|  | hobo-Activator | 235539 | 76509256 | 3.84 |
|  | Tc1-IS630-Pogo | 89322 | 27795795 | 1.39 |
|  | En-Spm | 0 | 0 | 0 |
|  | MuDR-IS905 | 0 | 0 | 0 |
|  | PiggyBac | 15495 | 6361713 | 0.32 |
|  | Tourist/Harbinger | 17379 | 2954184 | 0.15 |
|  | Other (Mirage, P-element, Transib) | 916 | 383021 | 0.02 |
| Rolling circles | All | 310078 | 109191298 | 5.48 |
| Unclassified | All | 2005869 | 45025041 | 22.5 |
| small RNA | All | 34585 | 37515640 | 1.88 |
| Satellites | All | 11798 | 3271725 | 0.16 |
| simple repeats | All | 255225 | 36150051 | 1.81 |
| Low complexity | All | 38363 | 2981959 | 0.15 |

Table S3. Median nucleotide diversity (π) per population and results of Mann-Whitney U tests comparing medians between highly differentiated windows and undifferentiated windows.

| **Transect** | **Pop** | **Host range** | **Median** | **n1** | **n2** | **statistic** | **p** | **Adj. p** | **Effect size** | **Effect mag.** |
| --- | --- | --- | --- | --- | --- | --- | --- | --- | --- | --- |
| West | CHST | Allo. | 0.0137 | 218 | 8968 | 831667 | 1.63E-04 | 2.17E-04 | 0.039 | small |
| West | CHSK | Sym. | 0.0129 | 218 | 7789 | 688322 | 1.81E-06 | 2.90E-06 | 0.053 | small |
| West | COSK | Sym. | 0.0140 | 218 | 9382 | 497110 | 1.37E-38 | 1.10E-37 | 0.133 | small |
| West | COGE | Allo. | 0.0136 | 218 | 9077 | 574845 | 3.38E-26 | 9.01E-26 | 0.110 | small |
| East | CHFI | Allo. | 0.0136 | 217 | 8677 | 832323 | 3.00E-03 | 3.43E-03 | 0.031 | small |
| East | CHES | Sym. | 0.0141 | 217 | 8975 | 898303 | 5.10E-02 | 5.10E-02 | 0.020 | small |
| East | COES | Sym. | 0.0143 | 217 | 9725 | 734278 | 1.67E-14 | 3.34E-14 | 0.077 | small |
| East | COLI | Allo. | 0.0139 | 217 | 9718 | 608935 | 1.56E-26 | 6.24E-26 | 0.107 | small |

Table S4. Median Tajima’s D per population and results of Mann-Whitney U tests comparing medians between highly differentiated windows and undifferentiated windows.

| **Transect** | **Pop.** | **Host range** | **Median** | **n1** | **n2** | **statistic** | **p** | **Adj. p** | **Effect size** | **Effect mag.** |
| --- | --- | --- | --- | --- | --- | --- | --- | --- | --- | --- |
| West | CHST | Allo. | -0.856 | 218 | 8968 | 887296 | 2.00E-02 | 2.29E-02 | 0.024 | small |
| West | CHSK | Sym. | -0.433 | 218 | 7789 | 697069.5 | 6.38E-06 | 1.28E-05 | 0.050 | small |
| West | COSK | Sym. | -0.731 | 218 | 9382 | 1562889 | 1.10E-40 | 8.80E-40 | 0.136 | small |
| West | COGE | Allo. | -0.787 | 218 | 9077 | 809931.5 | 4.57E-06 | 1.22E-05 | 0.048 | small |
| East | CHFI | Allo. | -0.829 | 217 | 8677 | 852696 | 1.80E-02 | 2.29E-02 | 0.025 | small |
| East | CHES | Sym. | -0.792 | 217 | 8975 | 1142586 | 1.24E-05 | 1.98E-05 | 0.046 | small |
| East | COES | Sym. | -0.806 | 217 | 9725 | 1081059 | 5.36E-01 | 5.36E-01 | 0.006 | small |
| East | COLI | Allo. | -0.805 | 217 | 9718 | 837140.5 | 2.00E-07 | 8.00E-07 | 0.052 | small |

Table S5. Median window-averaged extended haplotype homozygosity statistics (XP-EHH) and results of Mann-Whitney U tests comparing medians between highly differentiated (HD) windows and undifferentiated windows. More positive values indicate signatures of positive selection in Pop1 relative to Pop2, and more negative values indicate signatures of positive selection in Pop2 relative to Pop1.

| **Trans.** | **Pop 1** | **Pop 2** | **Host range** | **Win.** | **Median** | **n1** | **n2** | **statistic** | **p** | **Adj. p** | **Eff. size** | **Eff. mag.** |
| --- | --- | --- | --- | --- | --- | --- | --- | --- | --- | --- | --- | --- |
| West | COSK | CHSK | Sym. | 10175 | -0.0327 | 113 | 10024 | 954908 | 3.49E-36 | 5.24E-36 | 0.125 | small |
| West | COGE | CHST | Allo. | 10175 | -0.0669 | 115 | 10022 | 987764 | 1.04E-39 | 2.08E-39 | 0.131 | small |
| Betw. | CHES | CHSK | Sym. | 10167 | 0.0147 | 114 | 10035 | 371274 | 1.10E-10 | 1.10E-10 | 0.064 | small |
| Betw. | COES | COSK | Sym. | 10185 | 0.0051 | 113 | 10050 | 273363 | 2.22E-21 | 2.66E-21 | 0.094 | small |
| East | COES | CHES | Sym. | 10182 | -0.0538 | 133 | 10031 | 1132610 | 1.30E-43 | 3.90E-43 | 0.137 | small |
| East | COLI | CHFI | Allo. | 10175 | -0.0616 | 134 | 10012 | 1170683 | 7.91E-50 | 4.75E-49 | 0.147 | small |

Table S6. Selected SNPs and genes identified with XP-EHH (top 1% per comparison). Selected genes were identified as genes with 2000bp flanking regions that overlapped selected SNPs. Comparisons were designed to test for differences in selection between populations in sympatric host ranges, allopatric host ranges and contact zones.

| **Transect** | **Host range** | **Population pair** | **Selected in** | **Total selected SNPs** | **Selected SNPs in annotated genes** | **Genes containing selected SNPs** | **Selected SNPs per gene** |
| --- | --- | --- | --- | --- | --- | --- | --- |
| West | Sym. | CHSK - COSK | CHSK | 65803 | 30200 | 1026 | 29.4 |
| West | Sym. | CHSK - COSK | COSK | 90779 | 39686 | 571 | 69.5 |
| East | Sym. | CHES - COES | CHES | 75143 | 35026 | 1460 | 24 |
| East | Sym. | CHES - COES | COES | 82737 | 37373 | 634 | 58.9 |
| West | Allo. | CHST - COGE | CHST | 77972 | 35003 | 1466 | 23.9 |
| West | Allo. | CHST - COGE | COGE | 78446 | 35084 | 557 | 63 |
| East | Allo. | CHFI - COLI | CHFI | 65108 | 29765 | 1317 | 22.6 |
| East | Allo. | CHFI - COLI | COLI | 93999 | 39665 | 526 | 75.4 |
| Betw. | Sym. | CHSK - CHES | CHSK | 60937 | 28894 | 1065 | 27.1 |
| Betw. | Sym. | CHSK - CHES | CHES | 91993 | 44281 | 1873 | 23.6 |
| Betw. | Sym. | COSK - COES | COSK | 66883 | 31918 | 1405 | 22.7 |
| Betw | Sym. | COSK - COES | COES | 76716 | 36548 | 1591 | 23 |

Table S7. Genome-wide D-statistics and f4-ratios for all possible trios in the dataset, where P1 and P2 share the greatest number of derived alleles. This necessarily restricts testing to trees where P1 and P2 are from the same host race. Genome-wide estimates for the four trios utilized to calculate *f_dM_* are included among these.

| **P1** | **P2** | **P3** | **Dstatistic** | **Z-score** | **p-value** | **f4-ratio** | **BBAA** | **ABBA** | **BABA** |
| --- | --- | --- | --- | --- | --- | --- | --- | --- | --- |
| CHST | CHSK | COSK | 1.53E-02 | 18.0 | **2.30E-16** | 7.52E-02 | 173346 | 158433 | 153656 |
| CHST | CHSK | COGE | 1.19E-02 | 15.3 | **2.30E-16** | 6.13E-02 | 172558 | 158187 | 154470 |
| CHST | CHFI | COSK | 1.18E-02 | 14.9 | **2.30E-16** | 5.76E-02 | 170853 | 157226 | 153544 |
| CHST | CHES | COES | 1.12E-02 | 11.7 | **2.30E-16** | 6.11E-02 | 167730 | 157691 | 154187 |
| CHST | CHFI | COLI | 1.10E-02 | 14.8 | **2.30E-16** | 5.69E-02 | 170325 | 156407 | 153001 |
| CHST | CHFI | COES | 1.06E-02 | 15.2 | **2.30E-16** | 5.78E-02 | 170403 | 157805 | 154491 |
| CHST | CHES | COSK | 1.01E-02 | 10.8 | **2.30E-16** | 4.89E-02 | 168366 | 156522 | 153394 |
| CHST | CHSK | COES | 1.00E-02 | 13.2 | **2.30E-16** | 5.51E-02 | 173176 | 158046 | 154908 |
| CHST | CHSK | COLI | 1.00E-02 | 12.9 | **2.30E-16** | 5.22E-02 | 173166 | 156582 | 153478 |
| CHST | CHES | COLI | 9.79E-03 | 11.0 | **2.30E-16** | 5.04E-02 | 167791 | 155828 | 152808 |
| CHST | CHFI | COGE | 7.81E-03 | 11.1 | **2.30E-16** | 3.98E-02 | 170103 | 156828 | 154399 |
| COLI | COGE | CHSK | 6.79E-03 | 13.0 | **2.30E-16** | 2.74E-02 | 173392 | 155748 | 153647 |
| COSK | COES | CHES | 6.49E-03 | 6.7 | **2.04E-11** | 4.17E-02 | 173520 | 153856 | 151872 |
| COLI | COES | CHES | 6.43E-03 | 11.1 | **2.30E-16** | 4.10E-02 | 173580 | 153262 | 151302 |
| CHES | CHSK | COGE | 6.28E-03 | 6.3 | **3.03E-10** | 3.35E-02 | 171152 | 157971 | 156000 |
| CHST | CHES | COGE | 5.63E-03 | 6.8 | **1.03E-11** | 2.86E-02 | 167625 | 156016 | 154269 |
| COGE | COES | CHES | 5.55E-03 | 9.5 | **2.30E-16** | 3.60E-02 | 172379 | 154319 | 152615 |
| COSK | COGE | CHST | 5.42E-03 | 6.4 | **1.18E-10** | 3.02E-02 | 175521 | 151894 | 150258 |
| COSK | COES | CHST | 5.34E-03 | 5.3 | **1.30E-07** | 2.97E-02 | 175534 | 152372 | 150753 |
| CHES | CHSK | COSK | 5.27E-03 | 4.8 | **2.00E-06** | 2.74E-02 | 171711 | 157987 | 156332 |
| CHES | CHFI | CHSK | 5.24E-03 | 10.5 | **2.30E-16** | 2.80E-02 | 160222 | 164188 | 162476 |
| COLI | COGE | CHST | 5.00E-03 | 9.4 | **2.30E-16** | 2.79E-02 | 173418 | 151964 | 150450 |
| COLI | COSK | CHSK | 4.95E-03 | 7.2 | **4.56E-13** | 1.99E-02 | 175031 | 154676 | 153153 |
| COLI | COES | CHST | 4.89E-03 | 8.5 | **2.30E-16** | 2.71E-02 | 175641 | 151826 | 150349 |
| COLI | COES | CHSK | 4.84E-03 | 8.2 | **2.81E-16** | 1.95E-02 | 175751 | 155193 | 153697 |
| COLI | COES | CHFI | 4.42E-03 | 8.2 | **2.30E-16** | 2.44E-02 | 174448 | 153912 | 152559 |
| CHFI | CHSK | COGE | 4.08E-03 | 4.1 | **4.37E-05** | 2.21E-02 | 173112 | 158226 | 156940 |
| COSK | COES | CHFI | 3.97E-03 | 4.3 | **1.74E-05** | 2.22E-02 | 174321 | 154452 | 153231 |
| CHES | CHST | CHSK | 3.69E-03 | 5.5 | **3.77E-08** | 1.96E-02 | 158235 | 163040 | 161842 |
| CHFI | CHSK | COSK | 3.50E-03 | 3.2 | **1.28E-03** | 1.85E-02 | 173664 | 158252 | 157147 |
| CHES | CHST | CHFI | 3.14E-03 | 5.1 | **2.95E-07** | 2.38E-02 | 158505 | 160843 | 159837 |
| COGE | COES | CHFI | 2.64E-03 | 4.8 | **1.52E-06** | 1.49E-02 | 173178 | 154907 | 154091 |
| CHES | CHFI | COGE | 2.06E-03 | 3.4 | **7.63E-04** | 1.09E-02 | 168824 | 156543 | 155900 |
| COES | COGE | CHSK | 1.96E-03 | 3.4 | **5.84E-04** | 8.15E-03 | 174089 | 156399 | 155786 |
| COES | COGE | COSK | 1.94E-03 | 4.0 | **7.56E-05** | 1.56E-02 | 155858 | 158121 | 157508 |
| COLI | COGE | CHFI | 1.77E-03 | 3.4 | **8.01E-04** | 9.84E-03 | 172414 | 153371 | 152828 |
| COSK | COGE | CHSK | 1.76E-03 | 2.0 | **4.04E-02** | 7.27E-03 | 175099 | 155297 | 154753 |
| CHES | CHFI | COSK | 1.68E-03 | 2.5 | **1.18E-02** | 8.66E-03 | 169345 | 156708 | 156181 |
| CHST | CHFI | CHSK | 1.55E-03 | 2.7 | **7.30E-03** | 8.42E-03 | 160405 | 163350 | 162846 |
| COSK | COGE | CHFI | 1.20E-03 | 1.6 | 1.12E-01 | 6.71E-03 | 174496 | 153270 | 152902 |
| CHES | CHFI | COLI | 1.17E-03 | 1.8 | 7.14E-02 | 6.41E-03 | 168756 | 155860 | 155496 |
| CHSK | CHES | COES | 1.10E-03 | 1.0 | 3.24E-01 | 6.43E-03 | 171423 | 157856 | 157510 |
| CHSK | CHFI | COLI | 9.18E-04 | 0.9 | 3.76E-01 | 5.11E-03 | 173499 | 156716 | 156429 |
| COLI | COGE | CHES | 8.99E-04 | 1.6 | 9.96E-02 | 5.75E-03 | 171782 | 152082 | 151809 |
| COSK | COGE | CHES | 8.21E-04 | 1.1 | 2.83E-01 | 5.27E-03 | 173923 | 152029 | 151779 |
| CHFI | CHES | COES | 7.14E-04 | 1.0 | 3.40E-01 | 4.16E-03 | 168769 | 157407 | 157183 |
| COSK | COLI | CHST | 5.14E-04 | 0.8 | 4.43E-01 | 2.84E-03 | 175243 | 150148 | 149994 |
| COLI | COSK | CHFI | 5.06E-04 | 0.8 | 4.27E-01 | 2.80E-03 | 174019 | 152428 | 152274 |
| CHSK | CHFI | COES | 4.92E-04 | 0.5 | 6.13E-01 | 2.90E-03 | 173574 | 158106 | 157950 |
| CHES | CHSK | COLI | 3.37E-04 | 0.3 | 7.51E-01 | 1.87E-03 | 171483 | 156109 | 156004 |
| COLI | COGE | COSK | 3.08E-04 | 0.5 | 6.35E-01 | 2.49E-03 | 154413 | 156749 | 156652 |
| COES | COGE | CHST | 1.58E-04 | 0.3 | 7.79E-01 | 9.14E-04 | 174149 | 152630 | 152582 |
| COES | COSK | COLI | 1.28E-04 | 0.1 | 8.94E-01 | 1.16E-03 | 156028 | 156563 | 156523 |
| COES | COSK | CHSK | 1.04E-04 | 0.1 | 9.19E-01 | 4.31E-04 | 175289 | 155423 | 155390 |
| COLI | COES | COGE | 7.47E-05 | 0.1 | 9.06E-01 | 5.85E-04 | 158149 | 155980 | 155957 |
| COSK | COLI | CHES | 3.74E-05 | 0.1 | 9.55E-01 | 2.38E-04 | 173370 | 151256 | 151245 |

Table S8. Levene’s tests for homogeneity of variance between *f_dM_* in highly differentiated windows and undifferentiated windows for each tree tested (P1_P2_P3).

| **Trans.** | **P1_P2_P3** | **n1** | **n2** | **statistic** | **p** | **Adj. p** | **Eff. size** | **Eff. mag.** |
| --- | --- | --- | --- | --- | --- | --- | --- | --- |
| West | COGE_COSK_CHSK | 162 | 9418 | 3.92E+05 | 2.63E-26 | 7.01E-26 | 0.108 | small |
| West | COGE_COSK_CHST | 182 | 10650 | 5.35E+05 | 2.96E-25 | 5.92E-25 | 0.100 | small |
| West | CHST_CHSK_COGE | 176 | 9274 | 3.82E+05 | 1.06E-33 | 4.24E-33 | 0.124 | small |
| West | CHST_CHSK_COSK | 182 | 9531 | 3.75E+05 | 2.27E-39 | 1.82E-38 | 0.133 | small |
| East | COLI_COES_CHES | 223 | 11316 | 1.60E+06 | 3.48E-12 | 3.98E-12 | 0.065 | small |
| East | COLI_COES_CHFI | 220 | 10858 | 1.54E+06 | 2.25E-13 | 3.60E-13 | 0.070 | small |
| East | CHFI_CHES_COES | 233 | 10768 | 1.57E+06 | 7.31E-11 | 7.31E-11 | 0.062 | small |
| East | CHFI_CHES_COLI | 231 | 10633 | 1.56E+06 | 1.74E-12 | 2.32E-12 | 0.068 | small |

Table S9. Median fdM per tree tested (P1_P2_P3) and results of Mann-Whitney U tests comparing medians between highly differentiated windows and undifferentiated windows.

| **Trans.** | **P1_P2_P3** | **Win.** | **Med.** | **n1** | **n2** | **statistic** | **p** | **padj** | **Eff. size** | **Eff. mag.** |
| --- | --- | --- | --- | --- | --- | --- | --- | --- | --- | --- |
| West | COGE_COSK_CHSK | 11078 | 0.018 | 162 | 9418 | 3.92E+05 | 2.63E-26 | 7.01E-26 | 0.108 | small |
| West | COGE_COSK_CHST | 11539 | 0.021 | 182 | 10650 | 5.35E+05 | 2.96E-25 | 5.92E-25 | 0.100 | small |
| West | CHST_CHSK_COGE | 9713 | 0.044 | 176 | 9274 | 3.82E+05 | 1.06E-33 | 4.24E-33 | 0.124 | small |
| West | CHST_CHSK_COSK | 9450 | 0.037 | 182 | 9531 | 3.75E+05 | 2.27E-39 | 1.82E-38 | 0.133 | small |
| East | COLI_COES_CHES | 11001 | -0.018 | 223 | 11316 | 1.60E+06 | 3.48E-12 | 3.98E-12 | 0.065 | small |
| East | COLI_COES_CHFI | 10864 | -0.020 | 220 | 10858 | 1.54E+06 | 2.25E-13 | 3.60E-13 | 0.070 | small |
| East | CHFI_CHES_COES | 10832 | -0.009 | 233 | 10768 | 1.57E+06 | 7.31E-11 | 7.31E-11 | 0.062 | small |
| East | CHFI_CHES_COLI | 9580 | 0.001 | 231 | 10633 | 1.56E+06 | 1.74E-12 | 2.32E-12 | 0.068 | small |

Table S10. Significantly enriched biological processes (BP), molecular functions (MF) and cellular components (CC) in genes (n = 278) overlapping highly differentiated regions that were shared between all host race and transect comparisons (n = 214). Cat = GO category, Annot = annotated in the genome, sig = significant genes in the gene set, Exp. = expected significant genes based on annotation, P (CF) = p-value, classic one-sided Fisher’s exact test, P (PC) = p-value, Fisher’s exact test corrected for parent child relationships.

| **Cat** | **GO.ID** | **Term** | **Annot.** | **Sig.** | **Exp.** | **Rank** | **P (CF)** | **P (PC)** |
| --- | --- | --- | --- | --- | --- | --- | --- | --- |
| BP | GO:0010737 | protein kinase A signaling | 18 | 5 | 0.35 | 1 | 1.90E-05 | 9.80E-05 |
| BP | GO:0040012 | regulation of locomotion | 605 | 27 | 11.83 | 2 | 4.60E-05 | 1.70E-04 |
| BP | GO:0040017 | positive regulation of locomotion | 315 | 18 | 6.16 | 3 | 4.40E-05 | 2.00E-04 |
| BP | GO:0007297 | follicle cell of egg chamber migration | 140 | 11 | 2.74 | 4 | 9.10E-05 | 4.30E-04 |
| BP | GO:0038001 | paracrine signaling | 6 | 3 | 0.12 | 5 | 1.40E-04 | 5.40E-04 |
| BP | GO:0001661 | conditioned taste aversion | 8 | 3 | 0.16 | 6 | 3.80E-04 | 5.50E-04 |
| BP | GO:0040011 | locomotion | 1473 | 46 | 28.8 | 7 | 6.50E-04 | 6.50E-04 |
| BP | GO:0010738 | regulation of protein kinase A signaling | 15 | 4 | 0.29 | 8 | 1.60E-04 | 6.70E-04 |
| BP | GO:0051701 | biological process involved in interaction with host | 91 | 8 | 1.78 | 9 | 3.90E-04 | 7.20E-04 |
| BP | GO:0090183 | regulation of kidney development | 31 | 5 | 0.61 | 10 | 3.00E-04 | 9.00E-04 |
| BP | GO:2000278 | regulation of DNA biosynthetic process | 79 | 7 | 1.54 | 11 | 8.60E-04 | 9.70E-04 |
| BP | GO:0032879 | regulation of localization | 1537 | 49 | 30.05 | 12 | 2.50E-04 | 1.02E-03 |
| BP | GO:0051782 | negative regulation of cell division | 21 | 4 | 0.41 | 13 | 6.50E-04 | 1.08E-03 |
| BP | GO:2000573 | regulation of DNA biosynthetic process | 48 | 6 | 0.94 | 14 | 3.20E-04 | 1.15E-03 |
| BP | GO:0048144 | fibroblast proliferation | 53 | 6 | 1.04 | 15 | 5.50E-04 | 1.29E-03 |
| BP | GO:0051179 | localization | 3989 | 99 | 77.99 | 16 | 1.30E-03 | 1.30E-03 |
| BP | GO:0040040 | thermosensory behavior | 21 | 4 | 0.41 | 17 | 6.50E-04 | 1.47E-03 |
| BP | GO:0010496 | intercellular transport | 12 | 3 | 0.23 | 18 | 1.42E-03 | 1.53E-03 |
| BP | GO:0048145 | regulation of fibroblast proliferation | 52 | 6 | 1.02 | 19 | 5.00E-04 | 1.71E-03 |
| BP | GO:0043032 | positive regulation of macrophage activation | 6 | 2 | 0.12 | 20 | 5.42E-03 | 2.14E-03 |
| BP | GO:1901722 | regulation of cell proliferation involved in kidney development | 11 | 3 | 0.22 | 21 | 1.08E-03 | 2.32E-03 |
| BP | GO:0034113 | heterotypic cell-cell adhesion | 10 | 3 | 0.2 | 22 | 8.00E-04 | 2.32E-03 |
| BP | GO:0010557 | positive regulation of macromolecule biosynthetic process | 1432 | 38 | 28 | 23 | 2.84E-02 | 2.54E-03 |
| BP | GO:0007377 | germ-band extension | 10 | 3 | 0.2 | 24 | 8.00E-04 | 3.17E-03 |
| BP | GO:0051145 | smooth muscle cell differentiation | 30 | 4 | 0.59 | 25 | 2.61E-03 | 3.22E-03 |
| BP | GO:0090184 | positive regulation of kidney development | 23 | 4 | 0.45 | 26 | 9.40E-04 | 3.28E-03 |
| BP | GO:0001816 | cytokine production | 249 | 12 | 4.87 | 27 | 3.55E-03 | 3.40E-03 |
| BP | GO:0033631 | cell-cell adhesion mediated by integrin | 12 | 3 | 0.23 | 28 | 1.42E-03 | 3.47E-03 |
| BP | GO:0014009 | glial cell proliferation | 46 | 5 | 0.9 | 29 | 1.94E-03 | 3.53E-03 |
| BP | GO:0072666 | establishment of protein localization to vacuole | 33 | 3 | 0.65 | 30 | 2.61E-02 | 3.55E-03 |
| BP | GO:0035987 | endodermal cell differentiation | 32 | 4 | 0.63 | 31 | 3.32E-03 | 3.60E-03 |
| BP | GO:0007220 | Notch receptor processing | 15 | 3 | 0.29 | 32 | 2.81E-03 | 3.83E-03 |
| BP | GO:0098742 | cell-cell adhesion via plasma-membrane adhesion molecules | 84 | 7 | 1.64 | 33 | 1.25E-03 | 3.96E-03 |
| BP | GO:2000147 | positive regulation of cell motility | 224 | 12 | 4.38 | 34 | 1.48E-03 | 4.00E-03 |
| BP | GO:0035886 | vascular associated smooth muscle cell differentiation | 14 | 3 | 0.27 | 35 | 2.28E-03 | 4.38E-03 |
| BP | GO:0072111 | cell proliferation involved in kidney development | 15 | 3 | 0.29 | 36 | 2.81E-03 | 4.39E-03 |
| BP | GO:0032774 | RNA biosynthetic process | 2001 | 48 | 39.12 | 37 | 6.74E-02 | 4.40E-03 |
| BP | GO:2000145 | regulation of cell motility | 428 | 18 | 8.37 | 38 | 1.79E-03 | 4.43E-03 |
| BP | GO:0048870 | cell motility | 1000 | 32 | 19.55 | 39 | 3.44E-03 | 4.74E-03 |
| BP | GO:0009891 | positive regulation of biosynthetic process | 1505 | 40 | 29.42 | 40 | 2.40E-02 | 5.84E-03 |
| BP | GO:0008055 | ocellus pigment biosynthetic process | 25 | 3 | 0.49 | 41 | 1.23E-02 | 5.87E-03 |
| BP | GO:0032963 | collagen metabolic process | 38 | 4 | 0.74 | 42 | 6.22E-03 | 6.06E-03 |
| BP | GO:0046058 | cAMP metabolic process | 17 | 3 | 0.33 | 43 | 4.09E-03 | 6.06E-03 |
| BP | GO:0010745 | negative regulation of macrophage derived foam cell differentiation | 7 | 2 | 0.14 | 44 | 7.49E-03 | 6.21E-03 |
| BP | GO:0051764 | actin crosslink formation | 6 | 2 | 0.12 | 45 | 5.42E-03 | 6.31E-03 |
| BP | GO:0046158 | ocellus pigment metabolic process | 25 | 3 | 0.49 | 46 | 1.23E-02 | 6.33E-03 |
| BP | GO:0035162 | embryonic hemopoiesis | 61 | 5 | 1.19 | 47 | 6.65E-03 | 6.34E-03 |
| BP | GO:0007475 | apposition of dorsal and ventral imaginal disc-derived wing surfaces | 24 | 4 | 0.47 | 48 | 1.11E-03 | 6.51E-03 |
| BP | GO:0007540 | sex determination, establishment of X:A ratio | 6 | 2 | 0.12 | 49 | 5.42E-03 | 6.82E-03 |
| BP | GO:0048489 | synaptic vesicle transport | 169 | 9 | 3.3 | 50 | 5.90E-03 | 6.89E-03 |
| BP | GO:0006351 | DNA-templated transcription | 1945 | 45 | 38.03 | 51 | 1.20E-01 | 6.98E-03 |
| BP | GO:0001921 | positive regulation of receptor recycling | 6 | 2 | 0.12 | 52 | 5.42E-03 | 7.15E-03 |
| BP | GO:0036211 | protein modification process | 2199 | 58 | 42.99 | 53 | 6.88E-03 | 7.42E-03 |
| BP | GO:0042325 | regulation of phosphorylation | 804 | 27 | 15.72 | 54 | 3.81E-03 | 7.50E-03 |
| BP | GO:0044336 | canonical Wnt signaling pathway involved in negative regulation of apoptotic process | 5 | 2 | 0.1 | 55 | 3.66E-03 | 8.19E-03 |
| BP | GO:0051294 | establishment of spindle orientation | 77 | 5 | 1.51 | 56 | 1.72E-02 | 8.20E-03 |
| BP | GO:0022414 | reproductive process | 2034 | 54 | 39.77 | 57 | 8.30E-03 | 8.30E-03 |
| BP | GO:0010628 | positive regulation of gene expression | 1327 | 34 | 25.94 | 58 | 5.75E-02 | 8.34E-03 |
| BP | GO:0016926 | protein desumoylation | 7 | 2 | 0.14 | 59 | 7.49E-03 | 8.48E-03 |
| BP | GO:0072109 | glomerular mesangium development | 7 | 2 | 0.14 | 60 | 7.49E-03 | 8.94E-03 |
| BP | GO:0016925 | protein sumoylation | 26 | 3 | 0.51 | 61 | 1.37E-02 | 9.10E-03 |
| BP | GO:0036120 | cellular response to platelet-derived growth factor stimulus | 11 | 3 | 0.22 | 62 | 1.08E-03 | 9.19E-03 |
| BP | GO:0035556 | intracellular signal transduction | 1467 | 42 | 28.68 | 63 | 6.06E-03 | 9.32E-03 |
| BP | GO:0120033 | negative regulation of plasma membrane bounded cell projection assembly | 23 | 3 | 0.45 | 64 | 9.77E-03 | 9.50E-03 |
| BP | GO:0009887 | animal organ morphogenesis | 1199 | 36 | 23.44 | 65 | 5.41E-03 | 9.67E-03 |
| BP | GO:0051302 | regulation of cell division | 171 | 9 | 3.34 | 66 | 6.36E-03 | 9.97E-03 |
| BP | GO:0007440 | foregut morphogenesis | 22 | 3 | 0.43 | 67 | 8.62E-03 | 1.01E-02 |
| BP | GO:0048146 | positive regulation of fibroblast proliferation | 32 | 4 | 0.63 | 68 | 3.32E-03 | 1.02E-02 |
| BP | GO:0007299 | follicle cell of egg chamber-cell adhesion | 6 | 2 | 0.12 | 69 | 5.42E-03 | 1.02E-02 |
| BP | GO:0060439 | trachea morphogenesis | 16 | 3 | 0.31 | 70 | 3.41E-03 | 1.05E-02 |
| BP | GO:1905475 | regulation of protein localization to membrane | 99 | 7 | 1.94 | 71 | 3.20E-03 | 1.10E-02 |
| BP | GO:0036119 | response to platelet-derived growth factor | 12 | 3 | 0.23 | 72 | 1.42E-03 | 1.13E-02 |
| BP | GO:0051650 | establishment of vesicle localization | 268 | 13 | 5.24 | 73 | 2.30E-03 | 1.15E-02 |
| BP | GO:0072375 | medium-term memory | 22 | 4 | 0.43 | 74 | 7.90E-04 | 1.18E-02 |
| BP | GO:0140352 | export from cell | 857 | 27 | 16.75 | 75 | 8.84E-03 | 1.19E-02 |
| BP | GO:0032200 | telomere organization | 118 | 6 | 2.31 | 76 | 2.81E-02 | 1.20E-02 |
| BP | GO:0070848 | response to growth factor | 447 | 19 | 8.74 | 77 | 1.18E-03 | 1.21E-02 |
| BP | GO:0001817 | regulation of cytokine production | 217 | 10 | 4.24 | 78 | 1.02E-02 | 1.21E-02 |
| BP | GO:0010669 | epithelial structure maintenance | 31 | 5 | 0.61 | 79 | 3.00E-04 | 1.21E-02 |
| BP | GO:0016082 | synaptic vesicle priming | 28 | 3 | 0.55 | 80 | 1.68E-02 | 1.23E-02 |
| BP | GO:0034661 | ncRNA catabolic process | 30 | 3 | 0.59 | 81 | 2.03E-02 | 1.24E-02 |
| BP | GO:0060341 | regulation of cellular localization | 858 | 28 | 16.77 | 82 | 4.79E-03 | 1.25E-02 |
| BP | GO:0046152 | ommochrome metabolic process | 25 | 3 | 0.49 | 83 | 1.23E-02 | 1.29E-02 |
| BP | GO:0070371 | ERK1 and ERK2 cascade | 107 | 8 | 2.09 | 84 | 1.15E-03 | 1.31E-02 |
| BP | GO:0045935 | positive regulation of nucleobase-containing compound metabolic process | 1186 | 34 | 23.19 | 85 | 1.36E-02 | 1.32E-02 |
| BP | GO:1902018 | negative regulation of cilium assembly | 10 | 2 | 0.2 | 86 | 1.54E-02 | 1.32E-02 |
| BP | GO:0043066 | negative regulation of apoptotic process | 554 | 19 | 10.83 | 87 | 1.20E-02 | 1.34E-02 |
| BP | GO:0072665 | protein localization to vacuole | 36 | 3 | 0.7 | 88 | 3.28E-02 | 1.34E-02 |
| BP | GO:0051094 | positive regulation of developmental process | 908 | 29 | 17.75 | 89 | 5.53E-03 | 1.35E-02 |
| BP | GO:0040013 | negative regulation of locomotion | 167 | 9 | 3.26 | 90 | 5.46E-03 | 1.37E-02 |
| BP | GO:0034698 | response to gonadotropin | 24 | 3 | 0.47 | 91 | 1.10E-02 | 1.37E-02 |
| BP | GO:0090136 | epithelial cell-cell adhesion | 18 | 3 | 0.35 | 92 | 4.83E-03 | 1.38E-02 |
| BP | GO:0072012 | glomerulus vasculature development | 7 | 2 | 0.14 | 93 | 7.49E-03 | 1.42E-02 |
| BP | GO:0030705 | cytoskeleton-dependent intracellular transport | 154 | 9 | 3.01 | 94 | 3.21E-03 | 1.42E-02 |
| BP | GO:0051641 | cellular localization | 2597 | 66 | 50.77 | 95 | 8.68E-03 | 1.43E-02 |
| BP | GO:0019221 | cytokine-mediated signaling pathway | 211 | 10 | 4.13 | 96 | 8.45E-03 | 1.44E-02 |
| BP | GO:0031328 | positive regulation of cellular biosynthetic process | 1496 | 38 | 29.25 | 97 | 5.07E-02 | 1.44E-02 |
| BP | GO:0051174 | regulation of phosphorus metabolic process | 941 | 28 | 18.4 | 98 | 1.59E-02 | 1.45E-02 |
| BP | GO:0006195 | purine nucleotide catabolic process | 25 | 3 | 0.49 | 99 | 1.23E-02 | 1.49E-02 |
| BP | GO:1990314 | cellular response to insulin-like growth factor stimulus | 9 | 2 | 0.18 | 100 | 1.25E-02 | 1.50E-02 |
| BP | GO:0030182 | neuron differentiation | 1438 | 38 | 28.11 | 101 | 3.01E-02 | 1.52E-02 |
| BP | GO:0060216 | definitive hemopoiesis | 29 | 3 | 0.57 | 102 | 1.85E-02 | 1.52E-02 |
| BP | GO:1904375 | regulation of protein localization to cell periphery | 72 | 6 | 1.41 | 103 | 2.77E-03 | 1.54E-02 |
| BP | GO:0090130 | tissue migration | 309 | 13 | 6.04 | 104 | 7.61E-03 | 1.56E-02 |
| BP | GO:0048729 | tissue morphogenesis | 963 | 30 | 18.83 | 105 | 6.79E-03 | 1.60E-02 |
| BP | GO:0001706 | endoderm formation | 42 | 5 | 0.82 | 106 | 1.28E-03 | 1.62E-02 |
| BP | GO:2001141 | regulation of RNA biosynthetic process | 1872 | 47 | 36.6 | 107 | 3.61E-02 | 1.62E-02 |
| BP | GO:0008594 | photoreceptor cell morphogenesis | 21 | 3 | 0.41 | 108 | 7.55E-03 | 1.65E-02 |
| BP | GO:0099111 | microtubule-based transport | 161 | 9 | 3.15 | 109 | 4.31E-03 | 1.67E-02 |
| BP | GO:0010556 | regulation of macromolecule biosynthetic process | 2862 | 60 | 55.95 | 110 | 2.82E-01 | 1.67E-02 |
| BP | GO:0071363 | cellular response to growth factor stimulus | 437 | 19 | 8.54 | 111 | 9.00E-04 | 1.68E-02 |
| BP | GO:0006355 | regulation of DNA-templated transcription | 1819 | 45 | 35.56 | 112 | 4.99E-02 | 1.69E-02 |
| BP | GO:0050679 | positive regulation of epithelial cell proliferation | 85 | 6 | 1.66 | 113 | 6.29E-03 | 1.71E-02 |
| BP | GO:0001704 | formation of primary germ layer | 129 | 7 | 2.52 | 114 | 1.32E-02 | 1.74E-02 |
| BP | GO:0018130 | heterocycle biosynthetic process | 2540 | 59 | 49.66 | 115 | 7.25E-02 | 1.75E-02 |
| BP | GO:0072523 | purine-containing compound catabolic process | 45 | 4 | 0.88 | 116 | 1.13E-02 | 1.76E-02 |
| BP | GO:0009636 | response to toxic substance | 545 | 19 | 10.65 | 117 | 1.02E-02 | 1.78E-02 |
| BP | GO:0033690 | positive regulation of osteoblast proliferation | 8 | 2 | 0.16 | 118 | 9.85E-03 | 1.82E-02 |
| BP | GO:0006727 | ommochrome biosynthetic process | 25 | 3 | 0.49 | 119 | 1.23E-02 | 1.82E-02 |
| BP | GO:0001894 | tissue homeostasis | 166 | 9 | 3.25 | 120 | 5.26E-03 | 1.85E-02 |
| BP | GO:0043069 | negative regulation of programmed cell death | 563 | 19 | 11.01 | 121 | 1.41E-02 | 1.90E-02 |
| BP | GO:0002761 | regulation of myeloid leukocyte differentiation | 67 | 4 | 1.31 | 122 | 4.18E-02 | 1.92E-02 |
| BP | GO:0000003 | reproduction | 2124 | 54 | 41.52 | 123 | 1.92E-02 | 1.92E-02 |
| BP | GO:0033314 | mitotic DNA replication checkpoint signaling | 11 | 2 | 0.22 | 124 | 1.86E-02 | 1.93E-02 |
| BP | GO:0051683 | establishment of Golgi localization | 9 | 2 | 0.18 | 125 | 1.25E-02 | 1.96E-02 |
| BP | GO:0007267 | cell-cell signaling | 1112 | 35 | 21.74 | 126 | 2.87E-03 | 1.97E-02 |
| BP | GO:0071897 | DNA biosynthetic process | 149 | 7 | 2.91 | 127 | 2.69E-02 | 1.97E-02 |
| BP | GO:0051291 | protein heterooligomerization | 101 | 4 | 1.97 | 128 | 1.36E-01 | 1.97E-02 |
| BP | GO:0030431 | sleep | 127 | 7 | 2.48 | 129 | 1.22E-02 | 1.97E-02 |
| BP | GO:0001892 | embryonic placenta development | 44 | 4 | 0.86 | 130 | 1.05E-02 | 1.98E-02 |
| BP | GO:0043030 | regulation of macrophage activation | 15 | 2 | 0.29 | 131 | 3.38E-02 | 1.99E-02 |
| BP | GO:0030949 | positive regulation of vascular endothelial growth factor receptor signaling pathway | 10 | 2 | 0.2 | 132 | 1.54E-02 | 2.00E-02 |
| BP | GO:0022008 | neurogenesis | 1654 | 42 | 32.34 | 133 | 4.05E-02 | 2.00E-02 |
| BP | GO:0071559 | response to transforming growth factor beta | 127 | 10 | 2.48 | 134 | 1.90E-04 | 2.00E-02 |
| BP | GO:1901362 | organic cyclic compound biosynthetic process | 2701 | 62 | 52.81 | 135 | 7.97E-02 | 2.01E-02 |
| BP | GO:0051656 | establishment of organelle localization | 516 | 20 | 10.09 | 136 | 2.63E-03 | 2.03E-02 |
| BP | GO:0033993 | response to lipid | 579 | 20 | 11.32 | 137 | 9.33E-03 | 2.04E-02 |
| BP | GO:0048598 | embryonic morphogenesis | 671 | 22 | 13.12 | 138 | 1.16E-02 | 2.05E-02 |
| BP | GO:0048512 | circadian behavior | 149 | 8 | 2.91 | 139 | 8.82E-03 | 2.05E-02 |
| BP | GO:0006623 | protein targeting to vacuole | 27 | 3 | 0.53 | 140 | 1.52E-02 | 2.08E-02 |
| BP | GO:0060562 | epithelial tube morphogenesis | 654 | 22 | 12.79 | 141 | 8.74E-03 | 2.08E-02 |
| BP | GO:0051100 | negative regulation of binding | 101 | 5 | 1.97 | 142 | 4.77E-02 | 2.09E-02 |
| BP | GO:0051054 | positive regulation of DNA metabolic process | 145 | 8 | 2.83 | 143 | 7.53E-03 | 2.09E-02 |
| BP | GO:0035152 | regulation of tube architecture, open tracheal system | 94 | 6 | 1.84 | 144 | 1.02E-02 | 2.13E-02 |
| BP | GO:0040028 | regulation of vulval development | 64 | 5 | 1.25 | 145 | 8.14E-03 | 2.17E-02 |
| BP | GO:0050746 | regulation of lipoprotein metabolic process | 12 | 2 | 0.23 | 146 | 2.21E-02 | 2.18E-02 |
| BP | GO:0060541 | respiratory system development | 366 | 14 | 7.16 | 147 | 1.26E-02 | 2.18E-02 |
| BP | GO:0035099 | hemocyte migration | 19 | 3 | 0.37 | 148 | 5.66E-03 | 2.19E-02 |
| BP | GO:0045743 | positive regulation of fibroblast growth factor receptor signaling pathway | 11 | 2 | 0.22 | 149 | 1.86E-02 | 2.21E-02 |
| BP | GO:0030213 | hyaluronan biosynthetic process | 5 | 2 | 0.1 | 150 | 3.66E-03 | 2.26E-02 |
| BP | GO:0043633 | polyadenylation-dependent RNA catabolic process | 13 | 2 | 0.25 | 151 | 2.57E-02 | 2.32E-02 |
| BP | GO:0010212 | response to ionizing radiation | 128 | 9 | 2.5 | 152 | 8.90E-04 | 2.34E-02 |
| BP | GO:0035107 | appendage morphogenesis | 427 | 15 | 8.35 | 153 | 2.02E-02 | 2.41E-02 |
| BP | GO:0051052 | regulation of DNA metabolic process | 275 | 11 | 5.38 | 154 | 1.92E-02 | 2.41E-02 |
| BP | GO:0006793 | phosphorus metabolic process | 1995 | 50 | 39 | 155 | 3.14E-02 | 2.44E-02 |
| BP | GO:1905332 | positive regulation of morphogenesis of an epithelium | 22 | 3 | 0.43 | 156 | 8.62E-03 | 2.47E-02 |
| BP | GO:0030148 | sphingolipid biosynthetic process | 76 | 4 | 1.49 | 157 | 6.13E-02 | 2.47E-02 |
| BP | GO:0010743 | regulation of macrophage derived foam cell differentiation | 11 | 2 | 0.22 | 158 | 1.86E-02 | 2.47E-02 |
| BP | GO:0000076 | DNA replication checkpoint signaling | 19 | 2 | 0.37 | 159 | 5.23E-02 | 2.52E-02 |
| BP | GO:0033060 | ocellus pigmentation | 25 | 3 | 0.49 | 160 | 1.23E-02 | 2.54E-02 |
| BP | GO:1900373 | positive regulation of purine nucleotide biosynthetic pathway | 27 | 2 | 0.53 | 161 | 9.71E-02 | 2.59E-02 |
| BP | GO:0019438 | aromatic compound biosynthetic process | 2562 | 59 | 50.09 | 162 | 8.31E-02 | 2.59E-02 |
| BP | GO:0033688 | regulation of osteoblast proliferation | 10 | 2 | 0.2 | 163 | 1.54E-02 | 2.60E-02 |
| BP | GO:0070303 | negative regulation of stress-activated protein kinase signaling cascade | 47 | 4 | 0.92 | 164 | 1.31E-02 | 2.61E-02 |
| BP | GO:0061437 | renal system vasculature development | 9 | 2 | 0.18 | 165 | 1.25E-02 | 2.62E-02 |
| BP | GO:0035041 | sperm DNA decondensation | 11 | 2 | 0.22 | 166 | 1.86E-02 | 2.63E-02 |
| BP | GO:0010468 | regulation of gene expression | 2738 | 56 | 53.53 | 167 | 3.71E-01 | 2.63E-02 |
| BP | GO:0051254 | positive regulation of RNA metabolic process | 1079 | 30 | 21.09 | 168 | 2.94E-02 | 2.70E-02 |
| BP | GO:0034097 | response to cytokine | 477 | 17 | 9.33 | 169 | 1.21E-02 | 2.70E-02 |
| BP | GO:0030951 | establishment or maintenance of microtubule cytoskeleton polarity | 56 | 4 | 1.09 | 170 | 2.36E-02 | 2.71E-02 |
| BP | GO:0010811 | positive regulation of cell-substrate adhesion | 47 | 4 | 0.92 | 171 | 1.31E-02 | 2.73E-02 |
| BP | GO:0007163 | establishment or maintenance of cell polarity | 358 | 13 | 7 | 172 | 2.34E-02 | 2.74E-02 |
| BP | GO:0060438 | trachea development | 28 | 3 | 0.55 | 173 | 1.68E-02 | 2.78E-02 |
| BP | GO:0051338 | regulation of transferase activity | 561 | 19 | 10.97 | 174 | 1.36E-02 | 2.80E-02 |
| BP | GO:0001919 | regulation of receptor recycling | 13 | 2 | 0.25 | 175 | 2.57E-02 | 2.82E-02 |
| BP | GO:1900037 | regulation of cellular response to hypoxia | 10 | 2 | 0.2 | 176 | 1.54E-02 | 2.83E-02 |
| BP | GO:1903361 | protein localization to basolateral plasma membrane | 10 | 2 | 0.2 | 177 | 1.54E-02 | 2.93E-02 |
| BP | GO:0001658 | branching involved in ureteric bud morphogenesis | 33 | 3 | 0.65 | 178 | 2.61E-02 | 2.95E-02 |
| BP | GO:0009314 | response to radiation | 490 | 18 | 9.58 | 179 | 7.40E-03 | 2.95E-02 |
| BP | GO:0009791 | post-embryonic development | 1071 | 32 | 20.94 | 180 | 9.44E-03 | 2.96E-02 |
| BP | GO:0098581 | detection of external biotic stimulus | 11 | 2 | 0.22 | 181 | 1.86E-02 | 2.98E-02 |
| BP | GO:0044068 | modulation by symbiont of host cellular process | 34 | 3 | 0.66 | 182 | 2.82E-02 | 3.00E-02 |
| BP | GO:0001675 | acrosome assembly | 12 | 2 | 0.23 | 183 | 2.21E-02 | 3.01E-02 |
| BP | GO:0030335 | positive regulation of cell migration | 218 | 12 | 4.26 | 184 | 1.17E-03 | 3.06E-02 |
| BP | GO:0034330 | cell junction organization | 610 | 20 | 11.93 | 185 | 1.58E-02 | 3.07E-02 |
| BP | GO:0034109 | homotypic cell-cell adhesion | 24 | 3 | 0.47 | 186 | 1.10E-02 | 3.10E-02 |
| BP | GO:0006890 | retrograde vesicle-mediated transport, Golgi to endoplasmic reticulum | 66 | 4 | 1.29 | 187 | 3.99E-02 | 3.10E-02 |
| BP | GO:0046847 | filopodium assembly | 84 | 5 | 1.64 | 188 | 2.42E-02 | 3.11E-02 |
| BP | GO:1902750 | negative regulation of cell cycle G2/M phase transition | 46 | 3 | 0.9 | 189 | 6.05E-02 | 3.12E-02 |
| BP | GO:2001171 | positive regulation of ATP biosynthetic process | 22 | 2 | 0.43 | 190 | 6.80E-02 | 3.13E-02 |
| BP | GO:0090077 | foam cell differentiation | 14 | 2 | 0.27 | 191 | 2.97E-02 | 3.15E-02 |
| BP | GO:1904837 | beta-catenin-TCF complex assembly | 16 | 2 | 0.31 | 192 | 3.81E-02 | 3.15E-02 |
| BP | GO:0022412 | cellular process involved in reproduction in multicellular organism | 1157 | 32 | 22.62 | 193 | 2.64E-02 | 3.17E-02 |
| BP | GO:0016482 | cytosolic transport | 176 | 9 | 3.44 | 194 | 7.63E-03 | 3.18E-02 |
| BP | GO:0051252 | regulation of RNA metabolic process | 2099 | 49 | 41.04 | 195 | 9.42E-02 | 3.20E-02 |
| BP | GO:0006807 | nitrogen compound metabolic process | 6216 | 128 | 121.52 | 196 | 1.85E-01 | 3.21E-02 |
| BP | GO:0042981 | regulation of apoptotic process | 929 | 25 | 18.16 | 197 | 6.20E-02 | 3.23E-02 |
| BP | GO:0048308 | organelle inheritance | 14 | 2 | 0.27 | 198 | 2.97E-02 | 3.33E-02 |
| BP | GO:2000811 | negative regulation of anoikis | 9 | 2 | 0.18 | 199 | 1.25E-02 | 3.33E-02 |
| BP | GO:0007398 | ectoderm development | 47 | 4 | 0.92 | 200 | 1.31E-02 | 3.34E-02 |
| BP | GO:0016192 | vesicle-mediated transport | 1239 | 34 | 24.22 | 201 | 2.46E-02 | 3.34E-02 |
| BP | GO:1904385 | cellular response to angiotensin | 14 | 2 | 0.27 | 202 | 2.97E-02 | 3.42E-02 |
| BP | GO:0050729 | positive regulation of inflammatory response | 35 | 3 | 0.68 | 203 | 3.05E-02 | 3.43E-02 |
| BP | GO:0043412 | macromolecule modification | 2355 | 59 | 46.04 | 204 | 1.87E-02 | 3.43E-02 |
| BP | GO:1902680 | positive regulation of RNA biosynthetic process | 953 | 28 | 18.63 | 205 | 1.85E-02 | 3.46E-02 |
| BP | GO:0021551 | central nervous system morphogenesis | 15 | 2 | 0.29 | 206 | 3.38E-02 | 3.46E-02 |
| BP | GO:0035051 | cardiocyte differentiation | 117 | 6 | 2.29 | 207 | 2.71E-02 | 3.49E-02 |
| BP | GO:0097352 | autophagosome maturation | 44 | 3 | 0.86 | 208 | 5.43E-02 | 3.51E-02 |
| BP | GO:0035039 | male pronucleus assembly | 13 | 2 | 0.25 | 209 | 2.57E-02 | 3.51E-02 |
| BP | GO:0033002 | muscle cell proliferation | 105 | 6 | 2.05 | 210 | 1.68E-02 | 3.53E-02 |
| BP | GO:0048660 | regulation of smooth muscle cell proliferation | 72 | 5 | 1.41 | 211 | 1.32E-02 | 3.54E-02 |
| BP | GO:0060479 | lung cell differentiation | 15 | 2 | 0.29 | 212 | 3.38E-02 | 3.54E-02 |
| BP | GO:0007615 | anesthesia-resistant memory | 17 | 3 | 0.33 | 213 | 4.09E-03 | 3.54E-02 |
| BP | GO:0007043 | cell-cell junction assembly | 133 | 7 | 2.6 | 214 | 1.54E-02 | 3.62E-02 |
| BP | GO:0007369 | gastrulation | 257 | 13 | 5.02 | 215 | 1.59E-03 | 3.63E-02 |
| BP | GO:0051301 | cell division | 490 | 16 | 9.58 | 216 | 3.05E-02 | 3.63E-02 |
| BP | GO:0033687 | osteoblast proliferation | 13 | 2 | 0.25 | 217 | 2.57E-02 | 3.64E-02 |
| BP | GO:0072006 | nephron development | 85 | 5 | 1.66 | 218 | 2.53E-02 | 3.65E-02 |
| BP | GO:0006952 | defense response | 813 | 23 | 15.89 | 219 | 4.58E-02 | 3.67E-02 |
| BP | GO:1901863 | positive regulation of muscle tissue devopment | 42 | 4 | 0.82 | 220 | 8.88E-03 | 3.68E-02 |
| BP | GO:0035282 | segmentation | 340 | 10 | 6.65 | 221 | 1.30E-01 | 3.72E-02 |
| BP | GO:0036151 | phosphatidylcholine acyl-chain remodeling | 7 | 2 | 0.14 | 222 | 7.49E-03 | 3.74E-02 |
| BP | GO:0009886 | post-embryonic animal morphogenesis | 498 | 17 | 9.74 | 223 | 1.79E-02 | 3.77E-02 |
| BP | GO:0051240 | positive regulation of multicellular organismal process | 1070 | 31 | 20.92 | 224 | 1.60E-02 | 3.80E-02 |
| BP | GO:0035160 | maintenance of epithelial integrity, open tracheal system | 12 | 3 | 0.23 | 225 | 1.42E-03 | 3.83E-02 |
| BP | GO:0048636 | positive regulation of muscle organ development | 45 | 4 | 0.88 | 226 | 1.13E-02 | 3.84E-02 |
| BP | GO:1902679 | negative regulation of RNA biosynthetic process | 848 | 24 | 16.58 | 227 | 4.16E-02 | 3.85E-02 |
| BP | GO:0007492 | endoderm development | 72 | 5 | 1.41 | 228 | 1.32E-02 | 3.88E-02 |
| BP | GO:0052652 | cyclic purine nucleotide metabolic process | 37 | 3 | 0.72 | 229 | 3.51E-02 | 3.92E-02 |
| BP | GO:0030574 | collagen catabolic process | 14 | 2 | 0.27 | 230 | 2.97E-02 | 3.92E-02 |
| BP | GO:0032873 | negative regulation of stress-activated MAPK cascade | 47 | 4 | 0.92 | 231 | 1.31E-02 | 3.99E-02 |
| BP | GO:0007610 | behavior | 1180 | 34 | 23.07 | 232 | 1.26E-02 | 4.00E-02 |
| BP | GO:1903358 | regulation of Golgi organization | 12 | 2 | 0.23 | 233 | 2.21E-02 | 4.01E-02 |
| BP | GO:0072073 | kidney epithelium development | 81 | 5 | 1.58 | 234 | 2.10E-02 | 4.01E-02 |
| BP | GO:0060947 | cardiac vascular smooth muscle cell differentiation | 7 | 2 | 0.14 | 235 | 7.49E-03 | 4.06E-02 |
| BP | GO:0007034 | vacuolar transport | 100 | 6 | 1.96 | 236 | 1.35E-02 | 4.08E-02 |
| BP | GO:0060491 | regulation of cell projection assembly | 154 | 7 | 3.01 | 237 | 3.14E-02 | 4.09E-02 |
| BP | GO:0007049 | cell cycle | 1395 | 37 | 27.27 | 238 | 3.07E-02 | 4.09E-02 |
| BP | GO:0090160 | Golgi to lysosome transport | 7 | 2 | 0.14 | 239 | 7.49E-03 | 4.10E-02 |
| BP | GO:0048738 | cardiac muscle tissue development | 132 | 8 | 2.58 | 240 | 4.31E-03 | 4.11E-02 |
| BP | GO:0001656 | metanephros development | 53 | 5 | 1.04 | 241 | 3.64E-03 | 4.15E-02 |
| BP | GO:0031346 | positive regulation of cell projection organization | 335 | 12 | 6.55 | 242 | 3.15E-02 | 4.20E-02 |
| BP | GO:0044238 | primary metabolic process | 6434 | 131 | 125.79 | 243 | 2.37E-01 | 4.20E-02 |
| BP | GO:0010563 | negative regulation of phosphorus metabolic process | 425 | 14 | 8.31 | 244 | 3.91E-02 | 4.20E-02 |
| BP | GO:0033209 | tumor necrosis factor-mediated signaling pathway | 33 | 4 | 0.65 | 245 | 3.72E-03 | 4.21E-02 |
| BP | GO:0021695 | cerebellar cortex development | 36 | 3 | 0.7 | 246 | 3.28E-02 | 4.23E-02 |
| BP | GO:0061326 | renal tubule development | 118 | 6 | 2.31 | 247 | 2.81E-02 | 4.26E-02 |
| BP | GO:0022402 | cell cycle process | 1178 | 32 | 23.03 | 248 | 3.29E-02 | 4.27E-02 |
| BP | GO:0046903 | secretion | 912 | 30 | 17.83 | 249 | 3.10E-03 | 4.27E-02 |
| BP | GO:0140029 | exocytic process | 66 | 4 | 1.29 | 250 | 3.99E-02 | 4.28E-02 |
| BP | GO:0010972 | negative regulation of G2/M transition of mitotic cell cycle | 41 | 3 | 0.8 | 251 | 4.56E-02 | 4.33E-02 |
| BP | GO:1990778 | protein localization to cell periphery | 198 | 9 | 3.87 | 252 | 1.57E-02 | 4.36E-02 |
| BP | GO:0009954 | proximal/distal pattern formation | 45 | 3 | 0.88 | 253 | 5.74E-02 | 4.37E-02 |
| BP | GO:0015718 | monocarboxylic acid transport | 105 | 5 | 2.05 | 254 | 5.47E-02 | 4.39E-02 |
| BP | GO:0061005 | cell differentiation involved in kidney development | 40 | 3 | 0.78 | 255 | 4.28E-02 | 4.41E-02 |
| BP | GO:1901343 | negative regulation of vasculature development | 52 | 3 | 1.02 | 256 | 8.11E-02 | 4.43E-02 |
| BP | GO:0061025 | membrane fusion | 153 | 8 | 2.99 | 257 | 1.03E-02 | 4.46E-02 |
| BP | GO:0046395 | carboxylic acid catabolic process | 231 | 8 | 4.52 | 258 | 8.34E-02 | 4.49E-02 |
| BP | GO:0006688 | glycosphingolipid biosynthetic process | 10 | 2 | 0.2 | 259 | 1.54E-02 | 4.53E-02 |
| BP | GO:0030858 | positive regulation of epithelial cell differentiation | 31 | 3 | 0.61 | 260 | 2.21E-02 | 4.54E-02 |
| BP | GO:0048580 | regulation of post-embryonic development | 133 | 7 | 2.6 | 261 | 1.54E-02 | 4.55E-02 |
| BP | GO:0007530 | sex determination | 62 | 4 | 1.21 | 262 | 3.28E-02 | 4.57E-02 |
| BP | GO:0035073 | pupariation | 16 | 2 | 0.31 | 263 | 3.81E-02 | 4.58E-02 |
| BP | GO:0030952 | establishment or maintenance of cytoskeleton polarity | 68 | 4 | 1.33 | 264 | 4.38E-02 | 4.58E-02 |
| BP | GO:0021697 | cerebellar cortex formation | 18 | 2 | 0.35 | 265 | 4.74E-02 | 4.59E-02 |
| BP | GO:0044346 | fibroblast apoptotic process | 15 | 2 | 0.29 | 266 | 3.38E-02 | 4.66E-02 |
| BP | GO:0097205 | renal filtration | 9 | 2 | 0.18 | 267 | 1.25E-02 | 4.67E-02 |
| BP | GO:0018208 | peptidyl-proline modification | 30 | 3 | 0.59 | 268 | 2.03E-02 | 4.67E-02 |
| BP | GO:0050789 | regulation of biological process | 5843 | 126 | 114.23 | 269 | 4.69E-02 | 4.69E-02 |
| BP | GO:0021675 | nerve development | 63 | 4 | 1.23 | 270 | 3.45E-02 | 4.70E-02 |
| BP | GO:0006869 | lipid transport | 232 | 9 | 4.54 | 271 | 3.84E-02 | 4.74E-02 |
| BP | GO:0009888 | tissue development | 1814 | 48 | 35.46 | 272 | 1.42E-02 | 4.75E-02 |
| BP | GO:0019058 | viral life cycle | 122 | 7 | 2.39 | 273 | 9.92E-03 | 4.77E-02 |
| BP | GO:0007626 | locomotory behavior | 380 | 16 | 7.43 | 274 | 3.16E-03 | 4.79E-02 |
| BP | GO:0014823 | response to activity | 64 | 4 | 1.25 | 275 | 3.63E-02 | 4.81E-02 |
| BP | GO:0032501 | multicellular organismal process | 5587 | 121 | 109.23 | 276 | 4.85E-02 | 4.85E-02 |
| BP | GO:0071236 | cellular response to antibiotic | 152 | 7 | 2.97 | 277 | 2.95E-02 | 4.90E-02 |
| BP | GO:0003008 | system process | 1411 | 39 | 27.59 | 278 | 1.45E-02 | 4.91E-02 |
| BP | GO:0061440 | kidney vasculature development | 9 | 2 | 0.18 | 279 | 1.25E-02 | 4.92E-02 |
| BP | GO:0002165 | instar larval or pupal development | 625 | 20 | 12.22 | 280 | 2.01E-02 | 4.94E-02 |
| BP | GO:0007154 | cell communication | 3339 | 78 | 65.28 | 281 | 3.14E-02 | 4.95E-02 |
| BP | GO:0050790 | regulation of catalytic activity | 1211 | 29 | 23.68 | 282 | 1.43E-01 | 4.97E-02 |
| BP | GO:0009187 | cyclic nucleotide metabolic process | 38 | 3 | 0.74 | 283 | 3.76E-02 | 5.00E-02 |
| MF | GO:0005161 | platelet-derived growth factor receptor binding | 7 | 3 | 0.13 | 1 | 2.10E-04 | 8.80E-04 |
| MF | GO:0022829 | wide pore channel activity | 8 | 3 | 0.15 | 2 | 3.40E-04 | 9.10E-04 |
| MF | GO:0005172 | vascular endothelial growth factor receptor binding | 7 | 3 | 0.13 | 3 | 2.10E-04 | 1.08E-03 |
| MF | GO:0005200 | structural constituent of cytoskeleton | 63 | 4 | 1.18 | 4 | 2.99E-02 | 2.78E-03 |
| MF | GO:0015252 | proton channel activity | 25 | 3 | 0.47 | 5 | 1.09E-02 | 4.04E-03 |
| MF | GO:0019783 | ubiquitin-like protein peptidase activity | 53 | 5 | 0.99 | 6 | 2.99E-03 | 4.88E-03 |
| MF | GO:0008234 | cysteine-type peptidase activity | 87 | 6 | 1.63 | 7 | 5.65E-03 | 8.49E-03 |
| MF | GO:0050840 | extracellular matrix binding | 24 | 3 | 0.45 | 8 | 9.73E-03 | 8.80E-03 |
| MF | GO:0140096 | catalytic activity, acting on a protein | 1283 | 37 | 23.99 | 9 | 3.96E-03 | 1.01E-02 |
| MF | GO:0001968 | fibronectin binding | 8 | 2 | 0.15 | 10 | 9.04E-03 | 1.03E-02 |
| MF | GO:0035612 | AP-2 adaptor complex binding | 9 | 2 | 0.17 | 11 | 1.15E-02 | 1.15E-02 |
| MF | GO:0046332 | SMAD binding | 46 | 4 | 0.86 | 12 | 1.04E-02 | 1.29E-02 |
| MF | GO:0004723 | calcium-dependent protein serine/threonine phosphatase activity | 6 | 2 | 0.11 | 13 | 4.96E-03 | 1.45E-02 |
| MF | GO:0030165 | PDZ domain binding | 68 | 5 | 1.27 | 14 | 8.68E-03 | 1.82E-02 |
| MF | GO:0004180 | carboxypeptidase activity | 37 | 4 | 0.69 | 15 | 4.81E-03 | 1.83E-02 |
| MF | GO:0032559 | adenyl ribonucleotide binding | 338 | 9 | 6.32 | 16 | 1.82E-01 | 2.14E-02 |
| MF | GO:0070566 | adenylyltransferase activity | 17 | 2 | 0.32 | 17 | 3.93E-02 | 2.64E-02 |
| MF | GO:0016810 | hydrolase activity, acting on carbon-nitrogen (but not peptide) bonds | 106 | 6 | 1.98 | 18 | 1.43E-02 | 3.34E-02 |
| MF | GO:0000987 | cis-regulatory region sequence-specific DNA binding | 257 | 8 | 4.81 | 19 | 1.09E-01 | 3.51E-02 |
| MF | GO:1990782 | protein tyrosine kinase binding | 50 | 4 | 0.93 | 20 | 1.39E-02 | 3.58E-02 |
| MF | GO:0003755 | peptidyl-prolyl cis-trans isomerase activity | 26 | 3 | 0.49 | 21 | 1.22E-02 | 3.62E-02 |
| MF | GO:0016859 | cis-trans isomerase activity | 29 | 3 | 0.54 | 22 | 1.64E-02 | 4.14E-02 |
| MF | GO:0005178 | integrin binding | 67 | 4 | 1.25 | 23 | 3.64E-02 | 4.31E-02 |
| MF | GO:0015026 | coreceptor activity | 22 | 2 | 0.41 | 24 | 6.29E-02 | 4.47E-02 |
| MF | GO:0005515 | protein binding | 4190 | 84 | 78.34 | 25 | 2.07E-01 | 4.57E-02 |
| MF | GO:0022803 | passive transmembrane transporter activity | 308 | 9 | 5.76 | 26 | 1.23E-01 | 4.58E-02 |
| MF | GO:0005543 | phospholipid binding | 208 | 8 | 3.89 | 27 | 4.12E-02 | 4.83E-02 |
| MF | GO:0003712 | transcription coregulator activity | 339 | 11 | 6.34 | 28 | 5.28E-02 | 4.86E-02 |
| MF | GO:0070300 | phosphatidic acid binding | 15 | 2 | 0.28 | 29 | 3.11E-02 | 4.92E-02 |
| CC | GO:0031528 | microvillus membrane | 15 | 3 | 0.29 | 1 | 2.68E-03 | 5.40E-04 |
| CC | GO:0035866 | alphav-beta3 integrin-PKCalpha complex | 5 | 2 | 0.1 | 2 | 3.53E-03 | 2.37E-03 |
| CC | GO:0098636 | protein complex involved in cell adhesion | 5 | 2 | 0.1 | 3 | 3.53E-03 | 2.66E-03 |
| CC | GO:0106068 | SUMO ligase complex | 6 | 2 | 0.12 | 4 | 5.23E-03 | 2.68E-03 |
| CC | GO:0031258 | lamellipodium membrane | 5 | 2 | 0.1 | 5 | 3.53E-03 | 2.70E-03 |
| CC | GO:0030054 | cell junction | 973 | 31 | 18.69 | 6 | 3.18E-03 | 3.27E-03 |
| CC | GO:0005921 | gap junction | 9 | 3 | 0.17 | 7 | 5.40E-04 | 3.43E-03 |
| CC | GO:0031527 | filopodium membrane | 6 | 2 | 0.12 | 8 | 5.23E-03 | 3.43E-03 |
| CC | GO:0090575 | RNA polymerase II transcription regulator complex | 118 | 6 | 2.27 | 9 | 2.60E-02 | 3.63E-03 |
| CC | GO:0008305 | integrin complex | 5 | 2 | 0.1 | 10 | 3.53E-03 | 1.08E-02 |

### Supplementary figures

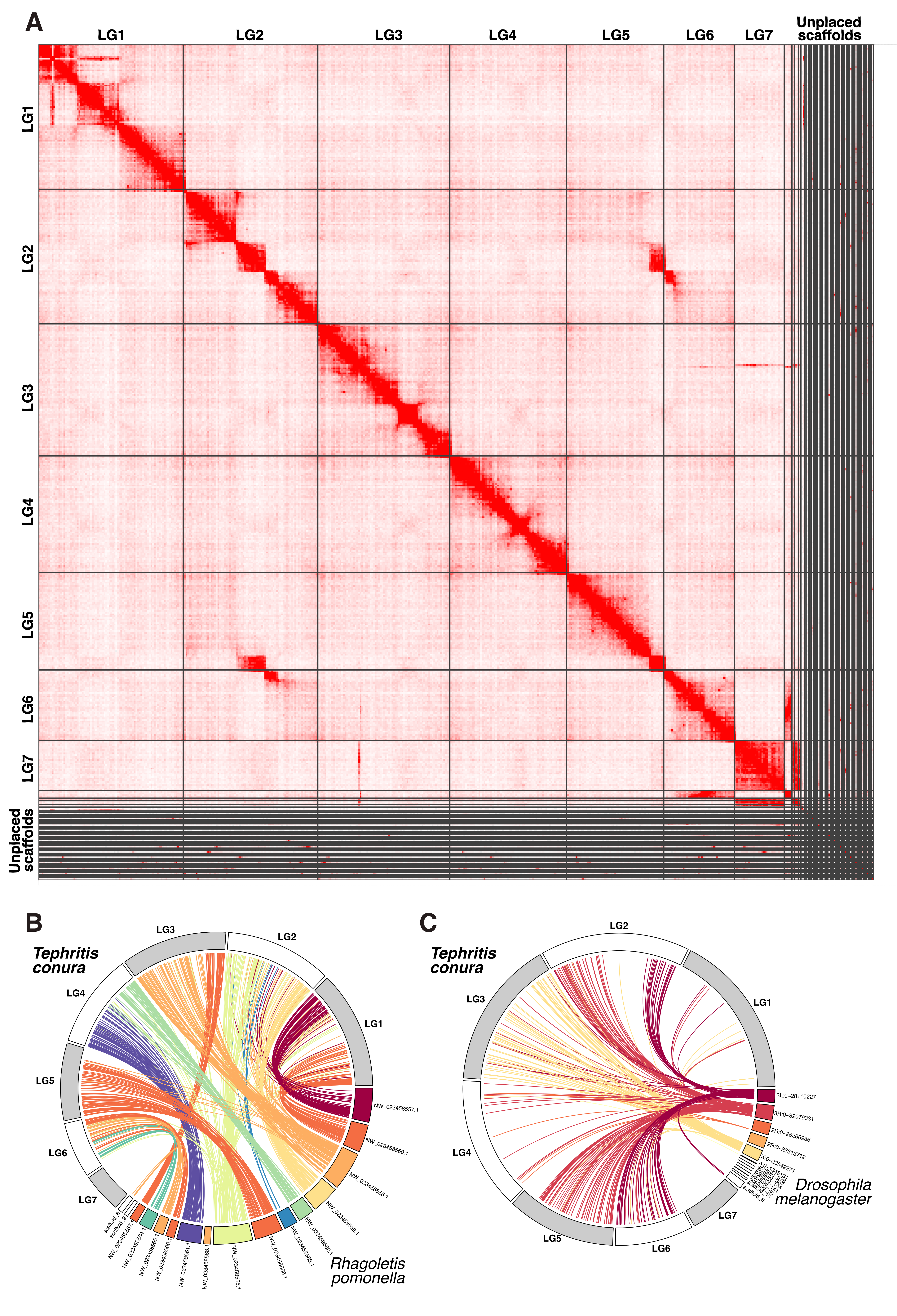

Figure S1. Hypothetical scaffolds of the T. conura genome and synteny with other dipteran genomes. (**A**) HiC contact map of scaffolded contigs. Contigs at least 50 kb long were scaffolded into larger linkage groups using YaHS. Circular synteny maps with (**B**) *Rhageletis pomonella* and (**C**) *Drosophila melanogaster.* The repetitive content in the *T. conura* genome was hardmasked prior to alignment with minimap. Linkage group (LG) 3 is syntenic with the *D. melanogaster* X chromosome, which is homologous with the ancestral dipteran X chromosome. LG7 did not align well with either the *R. pomonella* or the *D. melanogaster* genomes likely due to the high repetitive content of this putative Z chromosome. *T. conura* linkage groups 1-7 are shown in alternating gray and white, and are visualized this way in all following Manhattan plots.

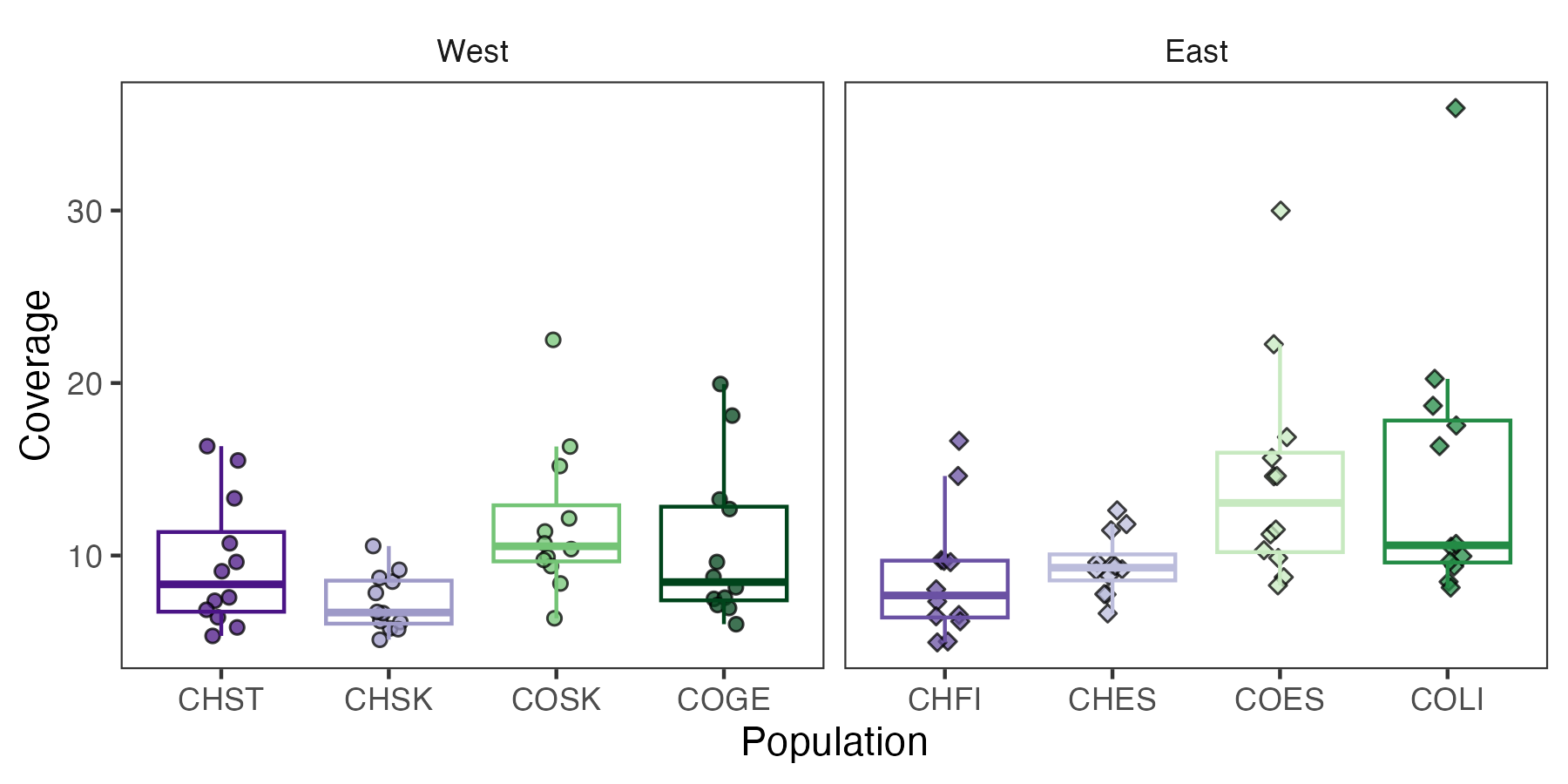

Figure S2. Genome-wide coverage of mapped and deduplicated reads varied among individuals. Points are genome-wide estimates for each individual. Boxplots showing medians (middle line), interquartile ranges (box), and 1.5x the interquartile range (whiskers) for each population. Points and boxplots are colored according to population (see Fig. 1, purple = CH host race, green = CO host race). Coverage was higher in CO host race populations (linear mixed model, log(coverage ~ host race + 1| population), X^2^=13.0, df = 1, p = 3.08x10^-4^).

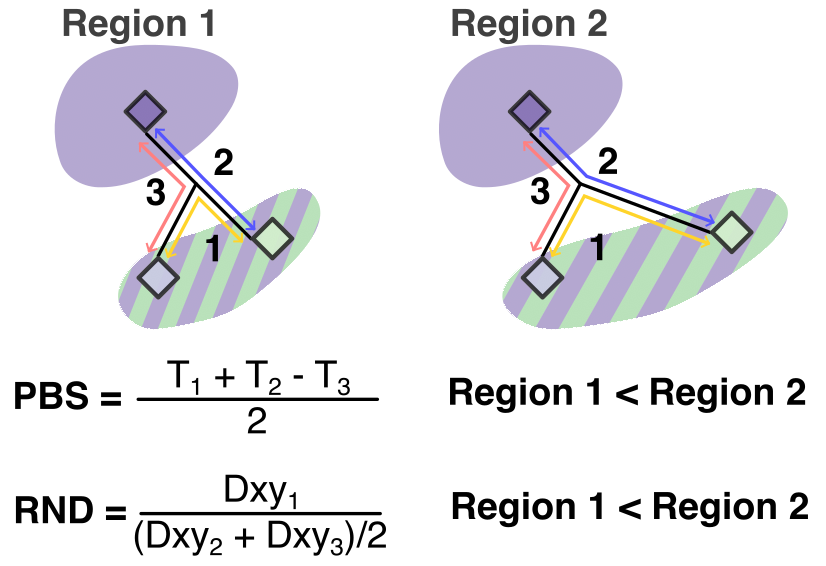

Figure S3. Schematic illustrating calculation of population branch statistics (PBS) and relative node depth (RND). PBS and RND quantify differentiation and divergence, respectively, between triads of populations. A population’s PBS describes genetic differentiation (where *T* = - log(1 – *F_ST_*)) at a locus or a cross the genome since divergence from two other populations. RND quantifies genetic divergence between two populations, relative to a third population (usually an outgroup, but here an allopatric population), and can be indicative of regions of reduced introgression. Here, the focal population is one in which host plants are in sympatry (CHSK, COSK, CHES, COES), contrasted with the sympatric (striped background) and allopatric (solid background) populations of the opposite host race within each transect. In the example pictured COES is compared to CHES and CHFI to identify regions of the genome where CHES is more differentiated and/or divergent. In these regions, PBS and RND will be higher than the remainder of the genome.

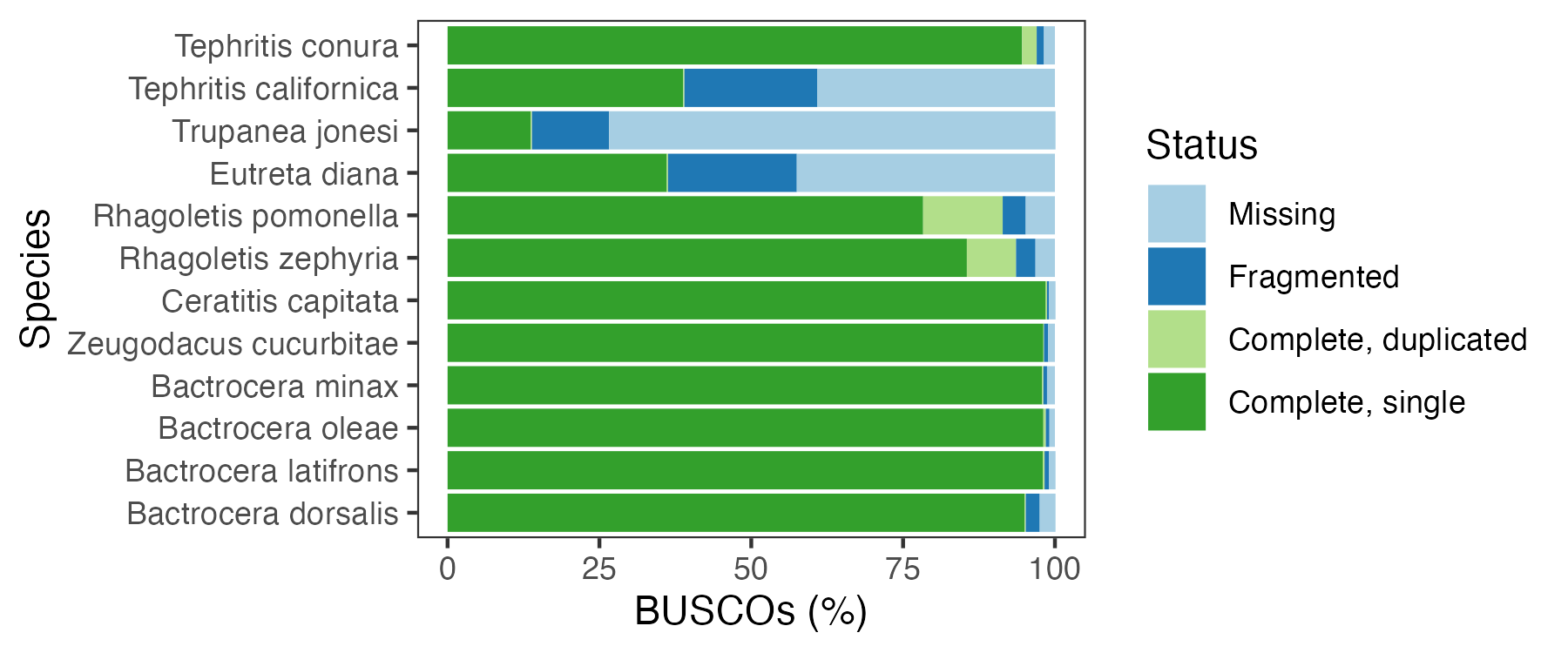
Figure S4. Benchmarking of single copy orthologs (BUSCOs) for published Tephritidae genomes available from BlobToolKit (<https://blobtoolkit.genomehubs.org/view/Tephritidae>; accessed 3/2023). Genomes were compared against nucleotide sequences of SCOs in the Diptera ODB10 database (n = 3286; Kriventseva *et al.*, 2019).

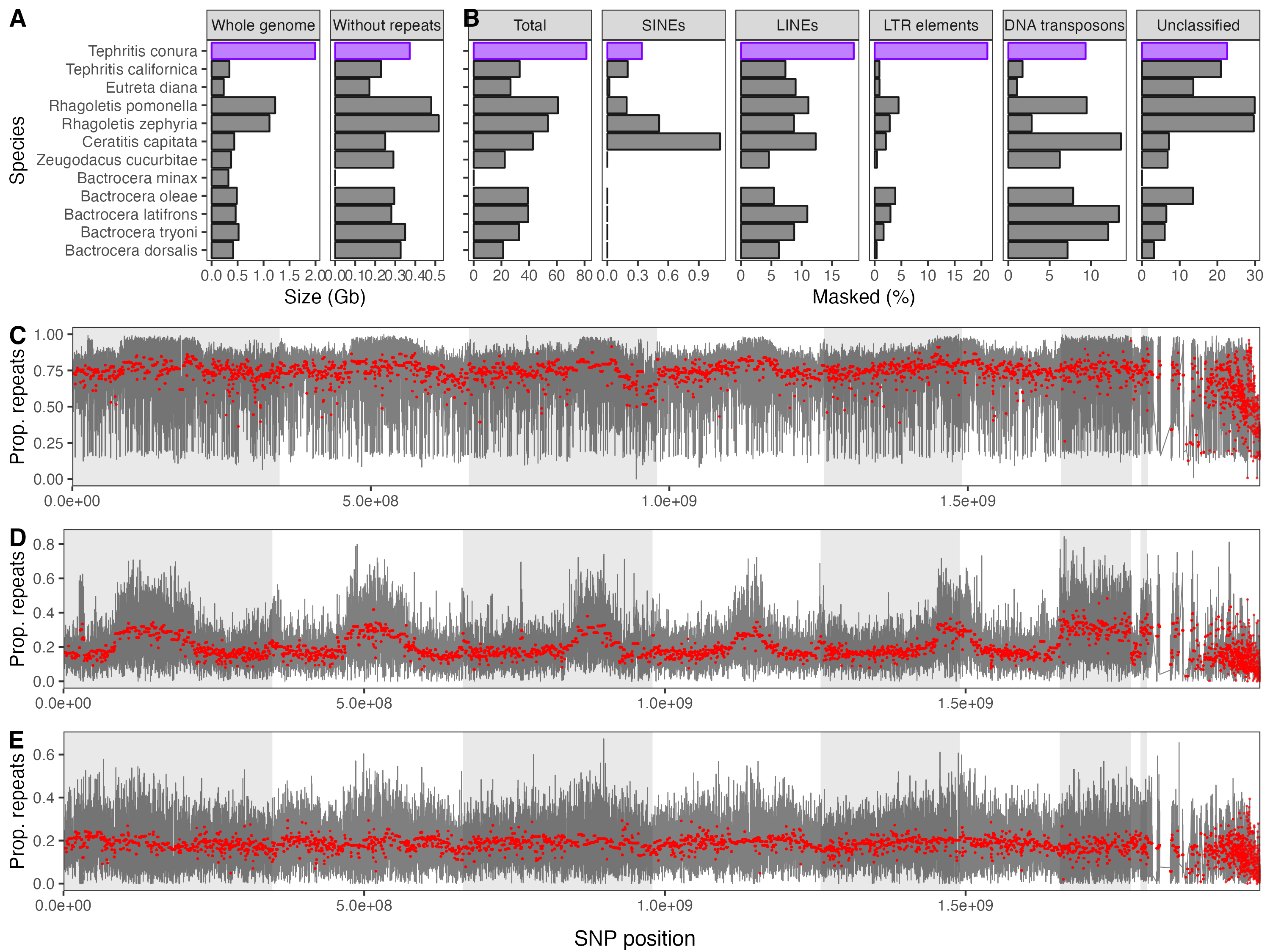

Figure S5. Genome size and repetitive content of *T. conura* and other published Tephritidae genomes. (**A**) Genome sizes with and without repetitive content. (**B**) Proportion of whole genome made up of all repeats (total), short interspersed nuclear elements (SINES), long interspersed nuclear elements (SINES), long terminal repeat (LTR) retrotransposons, DNA transposons and unclassified repeats. LTRs and LINEs are the main classes that are overrepresented in *T. conura* compared to other tephritids. Proportion of (**C**) total, (**D**) LTR retrotransposon and (**E**) LINE repetitive content in 50kb windows (gray lines) and full contigs (red dots) across the *T. conura* genome. Repeats were identified in the *T. conura* genome (CH host race) using Repeatmodeler and Repeatmasker. For all other genomes, we used repetitive content identified by Sproul et al. (2023) available from Heckenhauer et al. 2022 (<https://doi.org/10.6084/m9.figshare.c.6024905.v1>; accessed 3/2023), which did not include data for *Bactrocera minax*.

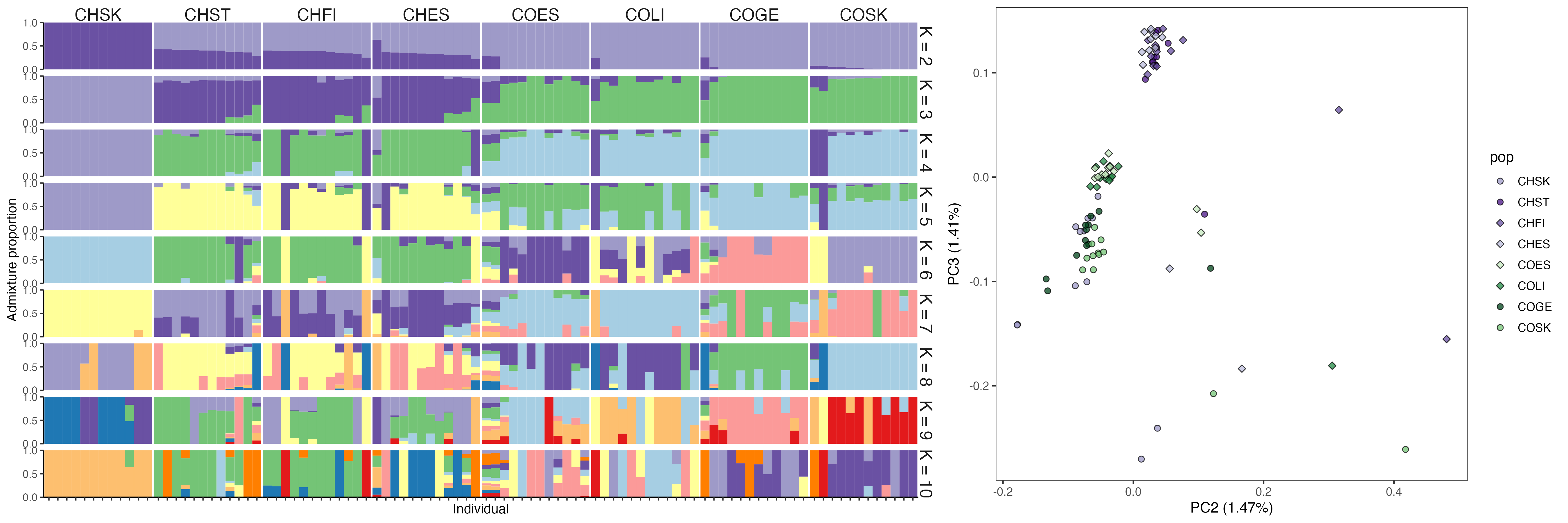

Figure S6. Genomic divergence among individuals (A) Admixture plots for K2-9. (**B**) Second and third principal components of whole genome differentiation between populations.

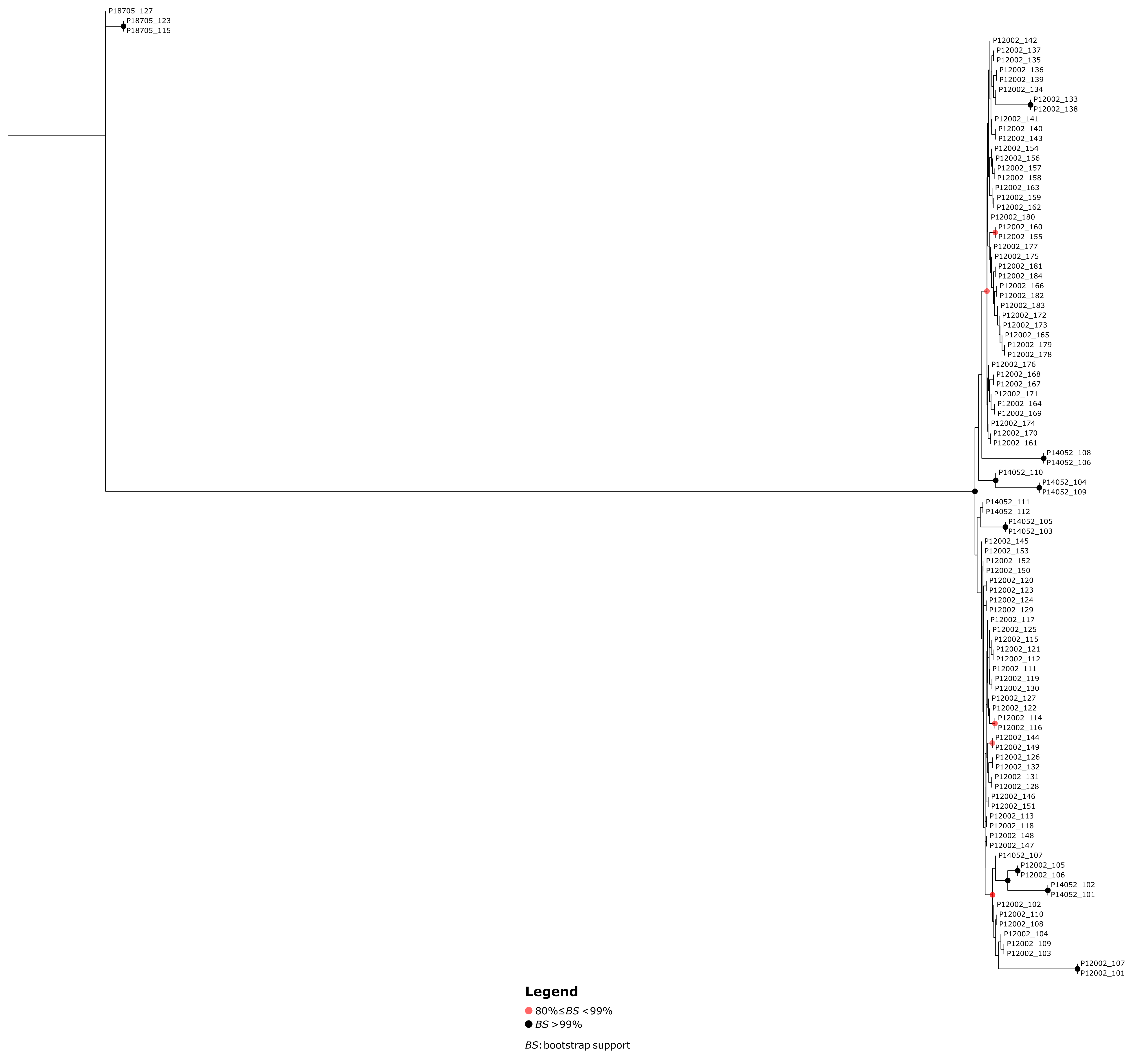

Figure S7. ASTRAL consensus tree of flies used in the study. Terminal nodes are colored by population and internal node support (posterior probability) is indicated and colored on gradient from blue (low support) to red (high support). Tree visualized using Tree Viewer (v. 2.2.0; <https://treeviewer.org/>).

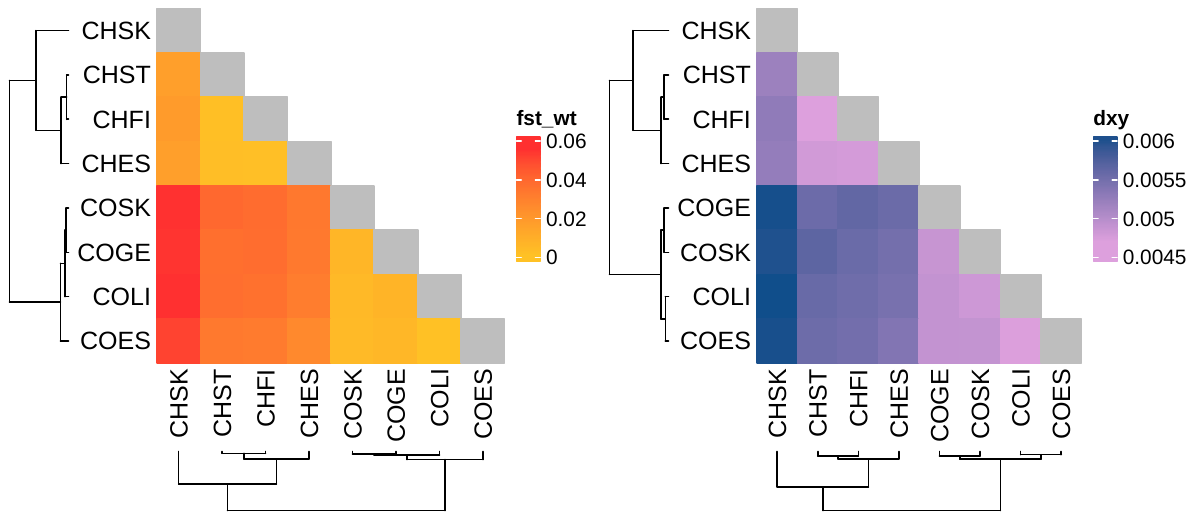

Figure S8. Genome-wide weighted ***F*_ST_** (Bhatia estimator) and dxy between all population pairs. Both statistics are most similar within host races (lighter colors). The sympatric CH population in the west (CHSK) is consistently the most different for both within and between host race comparisons.

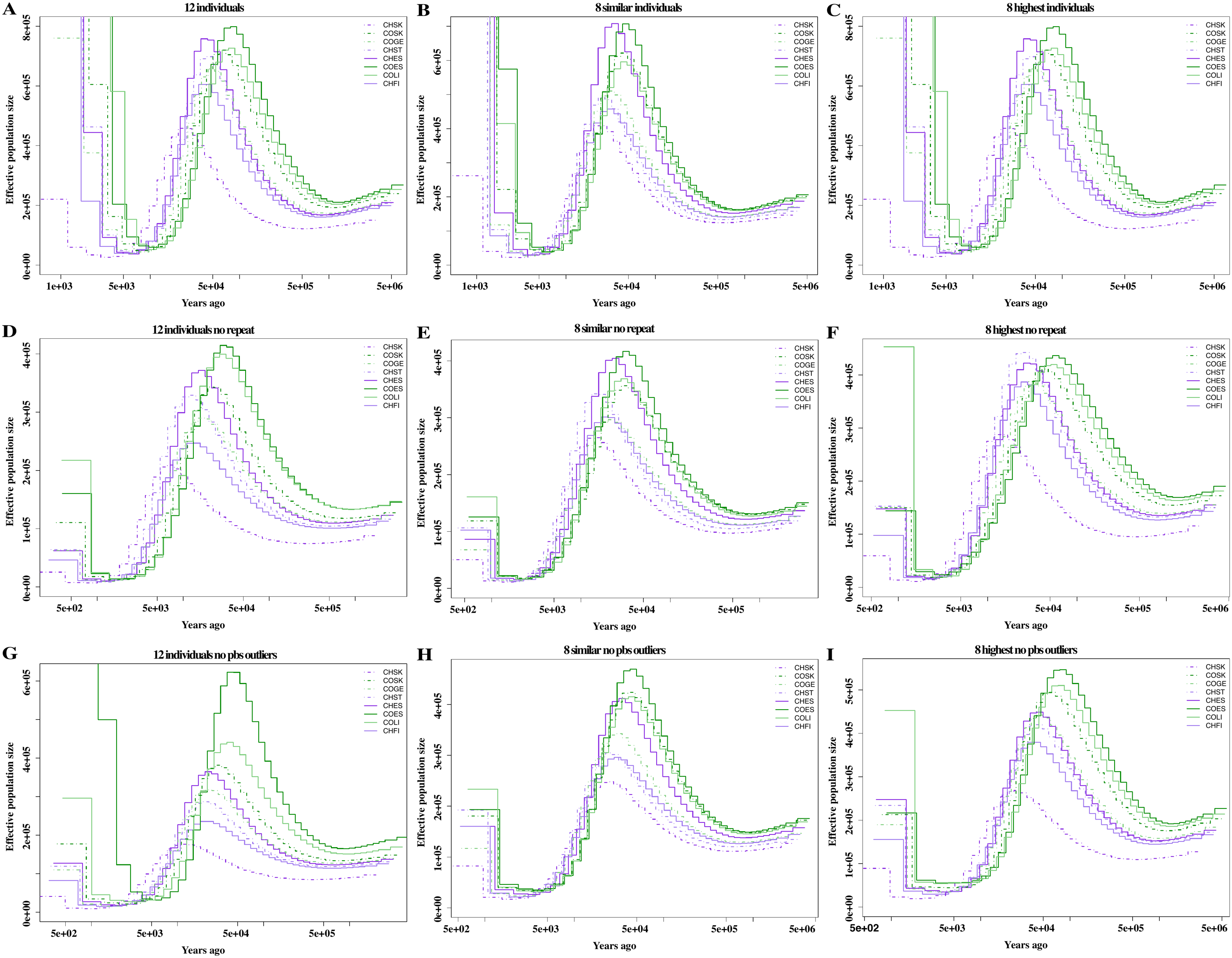

Figure S9. Effective population size estimates using MSMC2 compared between (**A**-**C**) full VCF, (**D**-**F**) VCF filtered for repetitive content and (**G**-**I**) VCF filtered for non-neutral sites (defined here as PBS outlier windows). Analyses were run on all 12 haplotypes per population, 8 haplotypes per population with the most similar coverage, and 8 haplotypes per population with the most similar coverage.

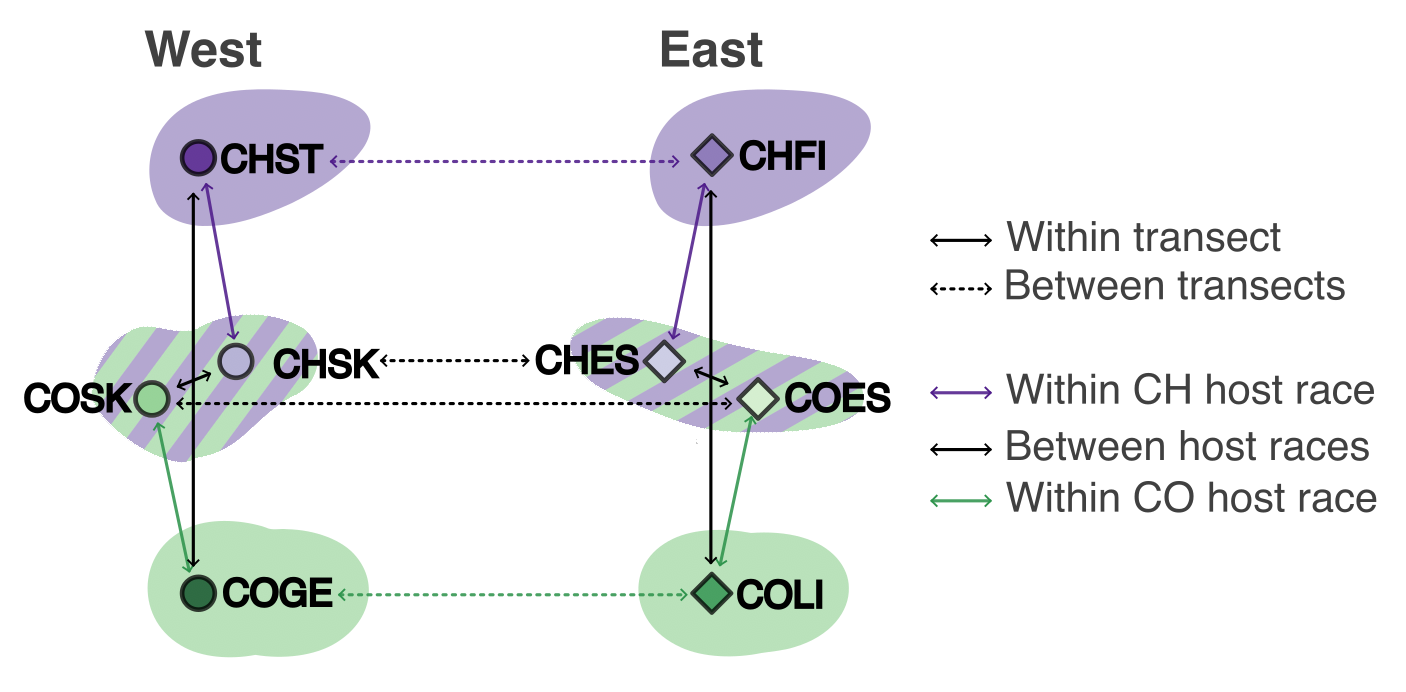

Figure S10. Subset of population pairs used for *F*_ST_ and dxy comparisons.

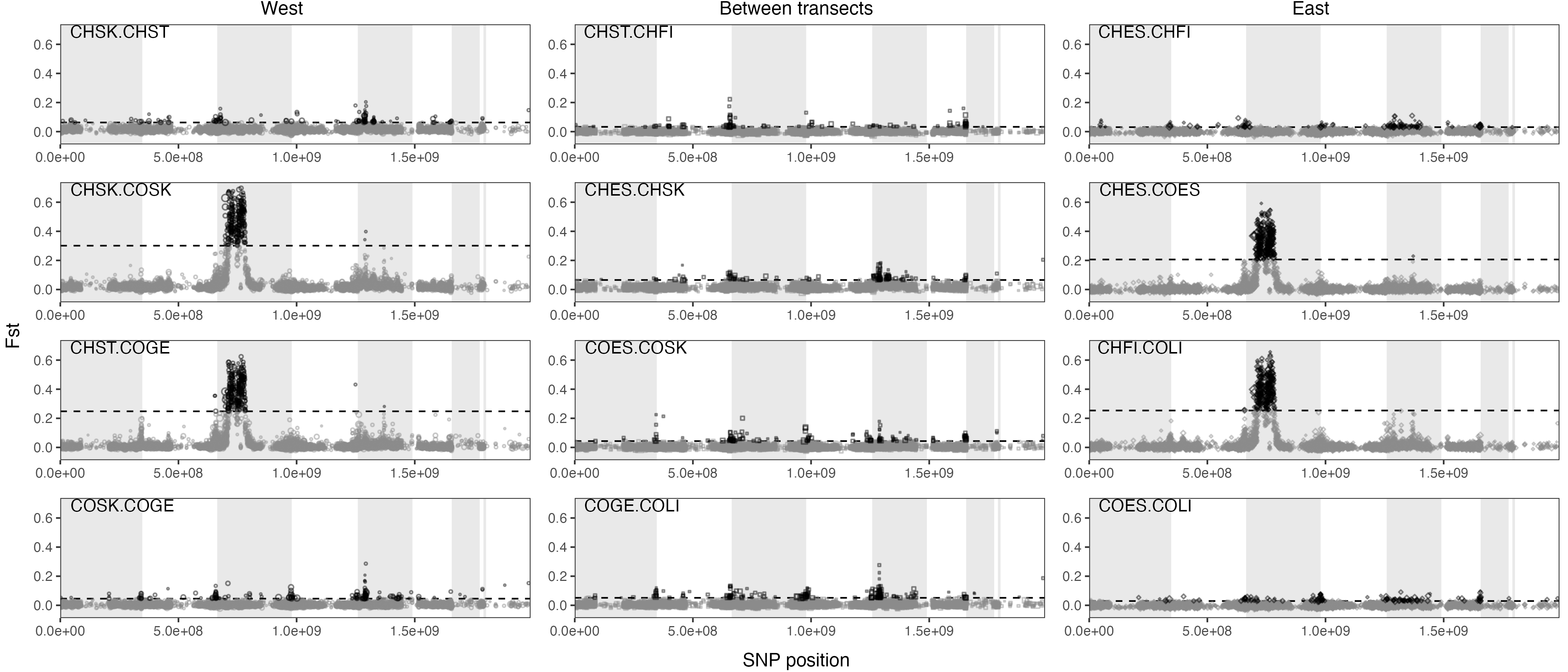

Figure S11. Differentiation (*F*_ST_) between pairs of *T. conura* populations in the west, east and between transects (see Fig. S10 for a schematic of focal comparisons). *F*_ST_ was calculated over nonoverlapping 50 kb windows. Dashed lines indicate three standard deviations from the mean and points falling above this threshold are shown in black. Contigs were ordered according to HiC scaffolds (Fig. S1) and hypothetical linkage groups are shown with alternating gray and white bands. The right-most white band represents unplaced scaffolds.

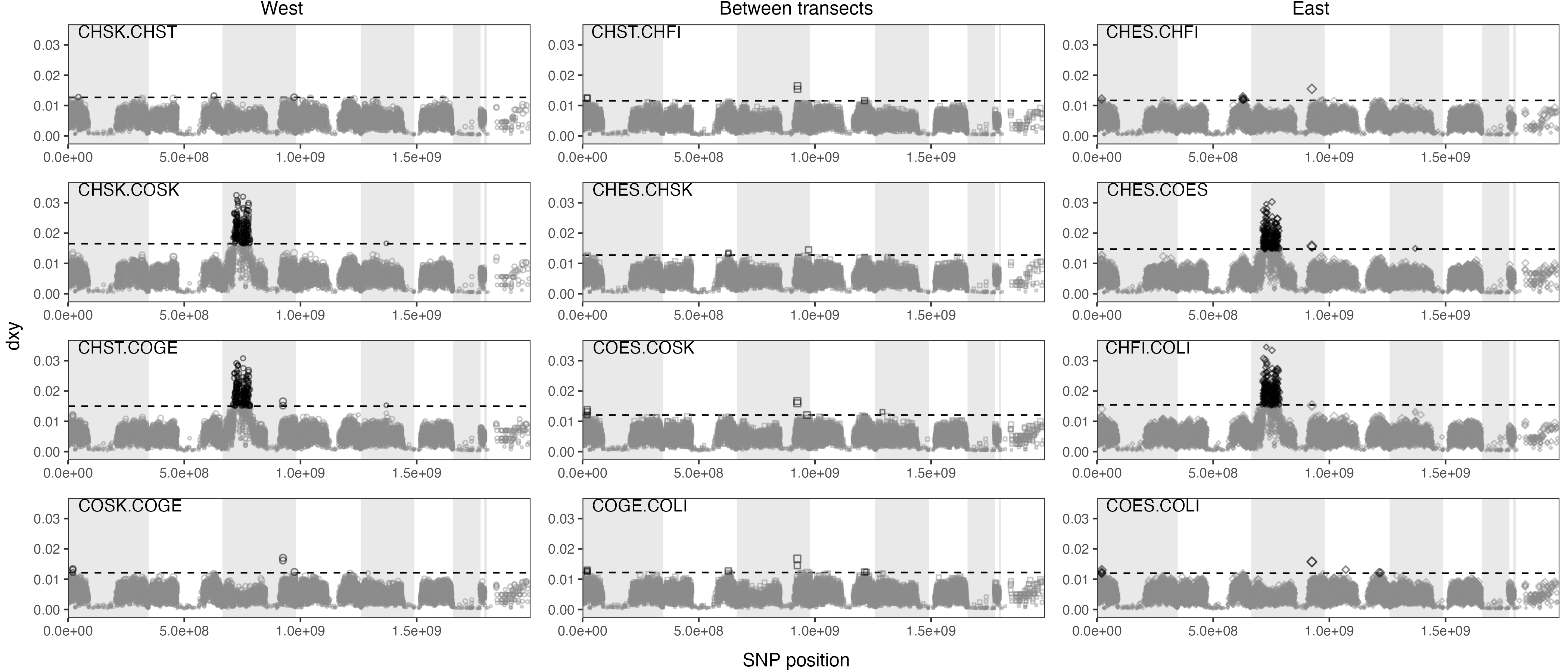

Figure S12. Absolute divergence (dxy) between pairs of T. conura populations in the west, east and between transects (see Fig. S10 for a schematic of focal comparisons). dxy was calculated over nonoverlapping 50 kb windows. Dashed lines indicate three standard deviations from the mean and points falling above this threshold are shown in black. Contigs were ordered according to HiC scaffolds (Fig. S1) and hypothetical linkage groups are shown with alternating gray and white bands. The right-most white band represents unplaced scaffolds.

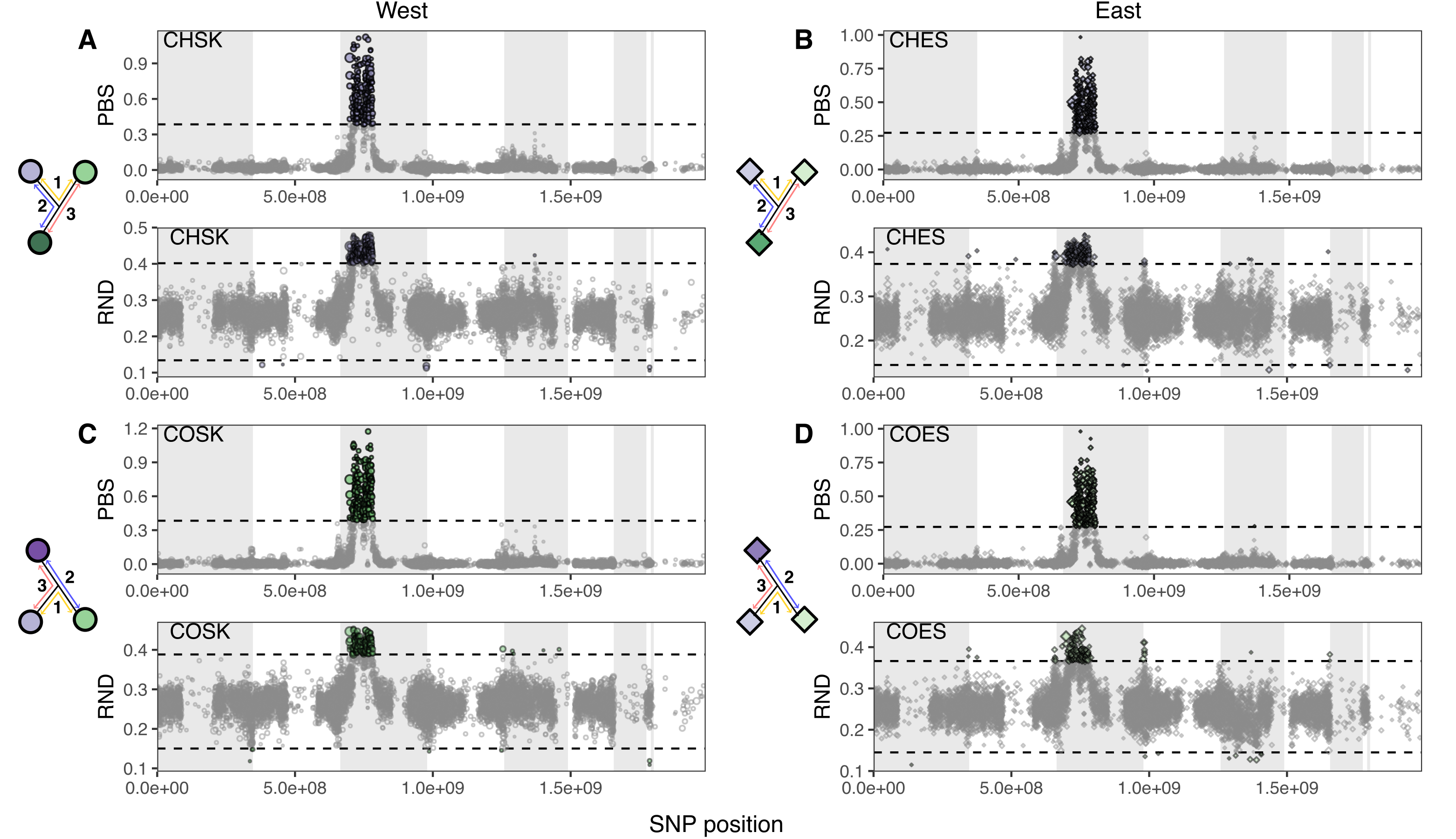

Figure S13. Population branch statistics (PBS) and relative node depth (RND) in 50 kb windows across the genome for *T. conura* population triads in the west and east. Windows with lower than 20% coverage were excluded. Triads (shown at left) were composed of a focal sympatric population of one host race, compared to the sympatric and allopatric populations of the other host race. Outlier windows are colored by population. Contigs were ordered according to HiC scaffolds (Fig. S1) and hypothetical linkage groups are shown with alternating gray and white bands. The right-most white band represents unplaced scaffolds.

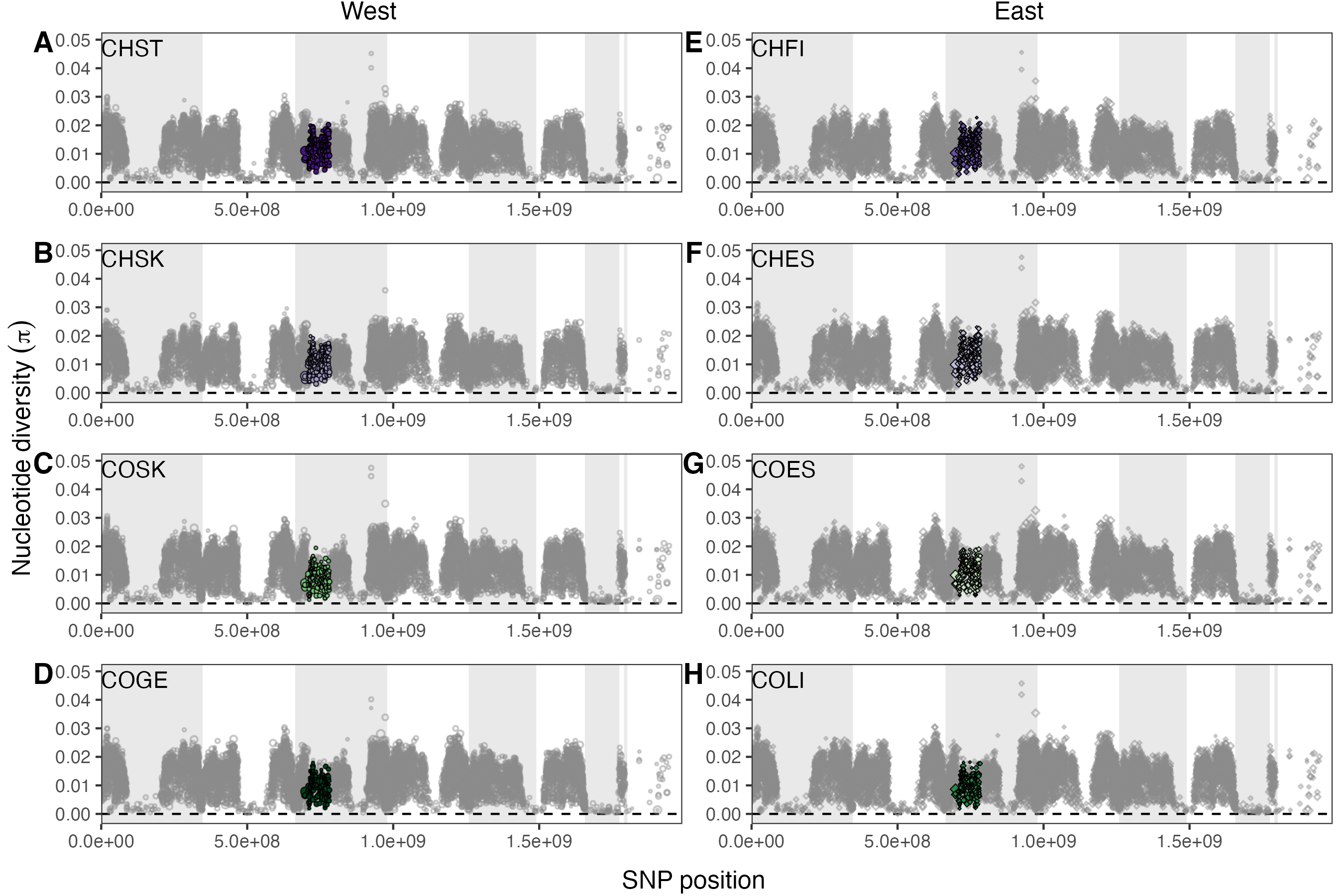

Figure S14. Nucleotide diversity (π) in 50 kb windows across the genome for *T. conura* populations in the west and east. Windows with lower than 20% coverage were excluded. Highly differentiated windows are colored by population. Contigs were ordered according to HiC scaffolds (Fig. S1) and hypothetical linkage groups are shown with alternating gray and white bands. The right-most white band represents unplaced scaffolds.

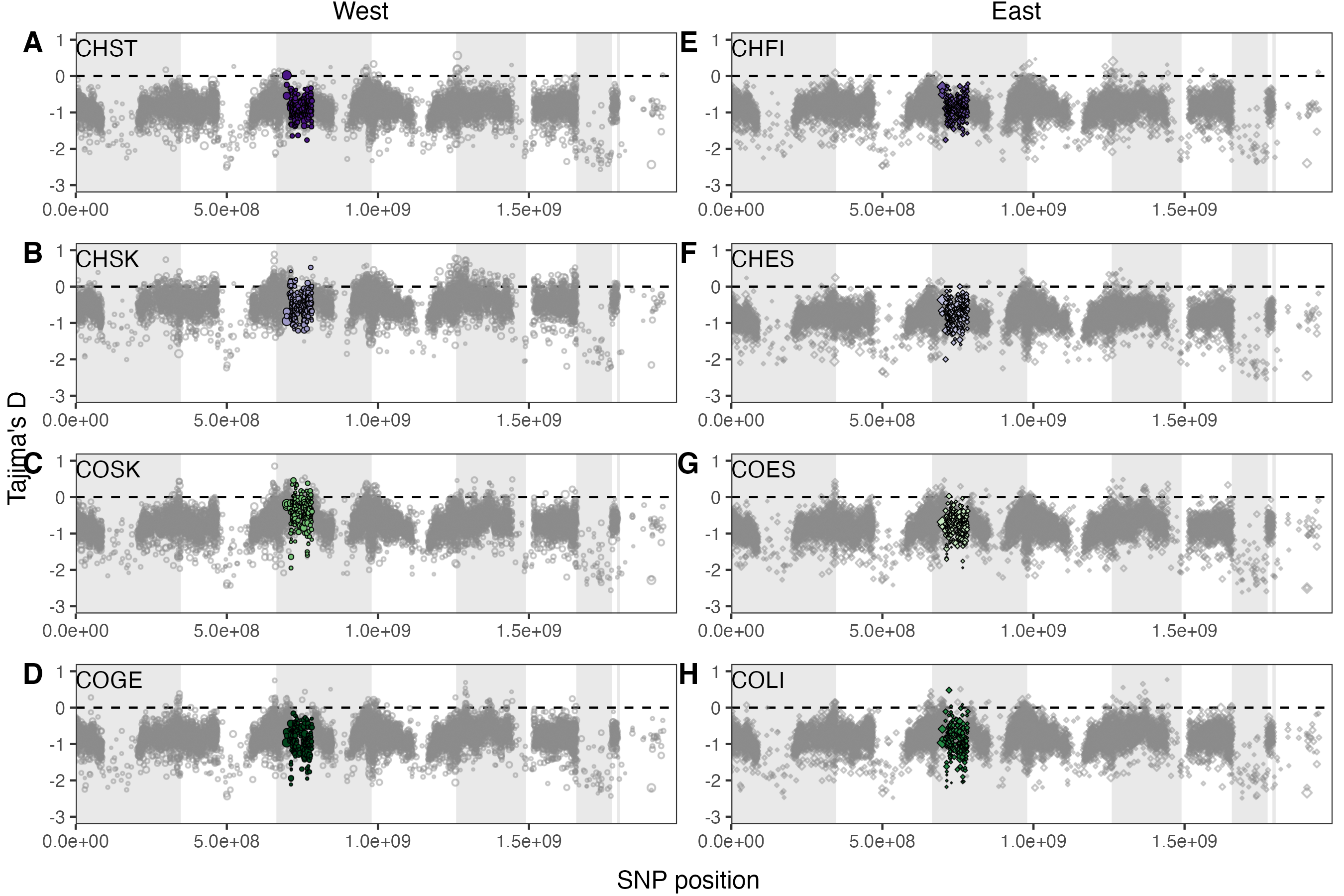

Figure S15. Tajima’s D in 50 kb windows across the genome for *T. conura* populations in the west and east. Windows with lower than 20% coverage were excluded. Highly differentiated windows are colored by population. Contigs were ordered according to HiC scaffolds (Fig. S1) and hypothetical linkage groups are shown with alternating gray and white bands. The right-most white band represents unplaced scaffolds.

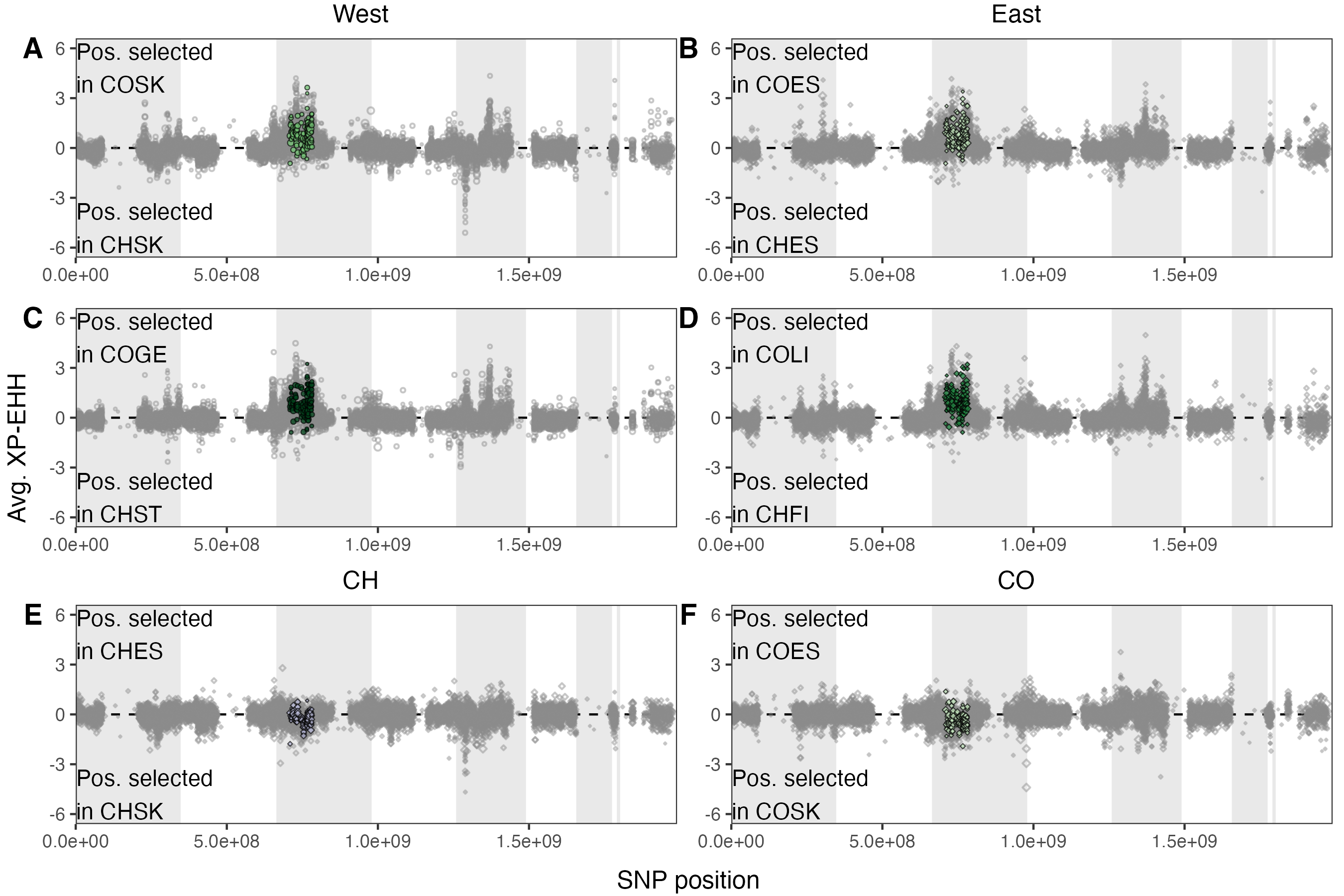

Figure S16. Tests for positive selection in 50 kb windows across the genome for pairwise contrasts of *T. conura* populations. Extended haplotype homozygosity (XP-EHH) was calculated between host races in (**A**-**B**) sympatry and (**C**-**D**) allopatry, and (**E**-**F**) within host races between eastern and western contact zones. More positive values indicate greater signatures of positive selection in one population, while more negative values indicate greater positive selection in the population. Windows with fewer than 500 SNPs (1% of the window) were excluded. Highly differentiated windows are colored by the population in which positive values indicate positive selection. Contigs were ordered according to HiC scaffolds (Fig. S1) and hypothetical linkage groups are shown with alternating gray and white bands. The right-most white band represents unplaced scaffolds.

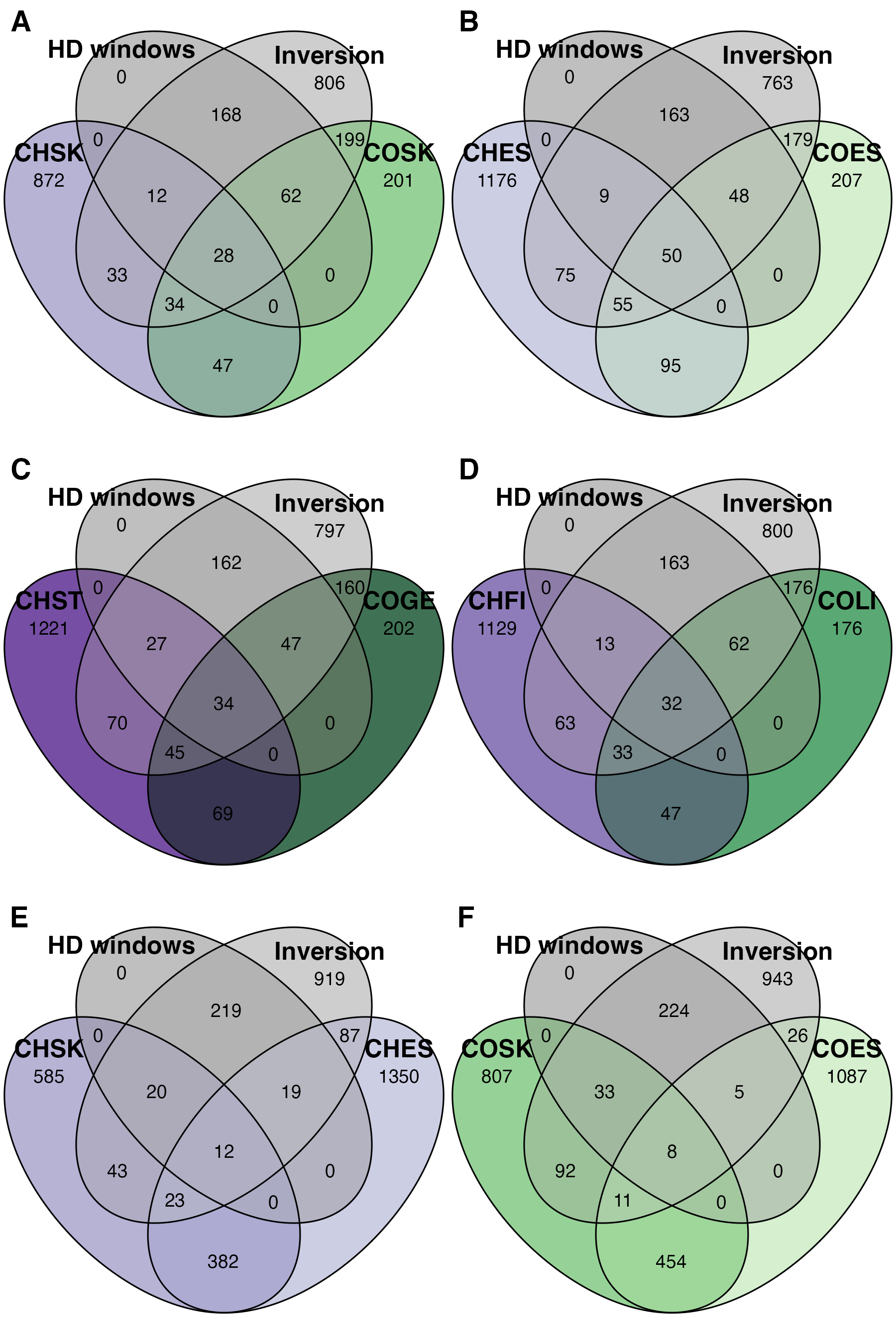

Figure S17. Overlap between genes in highly differentiated (HD) windows, genes in the inversion, and selected genes in pairwise contrasts. HD windows were identified as those overlapping outlier windows from the BayPass, PBS, and RND analyses, n = 309 genes. Inversion genes are those falling within the putative inversion region (Fig. 3), n = 1342 genes. All HD windows fall within the inversion. Selected genes were those containing XP-EHH outlier SNPs in pairwise contrasts between host races in sympatry or allopatry, or within host races between contact zones.

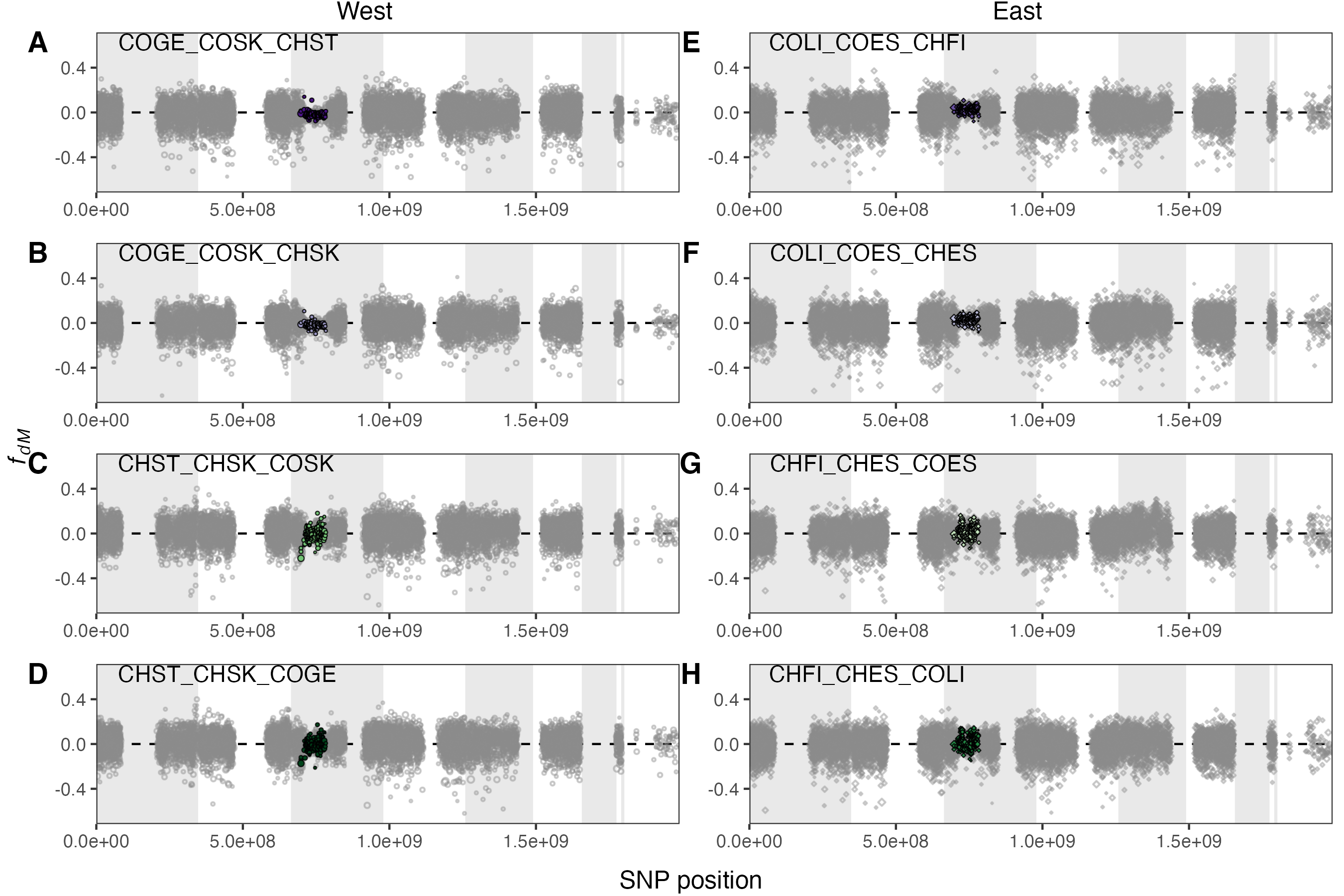

Figure S18. *f*_dM_ in 50 kb windows across the genome for *T. conura* trees (P1_P2_P3) in the west and east. Windows with lower than 500 sites (1% of window) excluded. Highly differentiated windows are colored by P3. Contigs were ordered according to HiC scaffolds (Fig. S1) and hypothetical linkage groups are shown with alternating gray and white bands. The right-most white band represents unplaced scaffolds.
